# Gentle rhodamines for live-cell fluorescence microscopy

**DOI:** 10.1101/2024.02.06.579089

**Authors:** Tianyan Liu, Julian Kompa, Jing Ling, Nicolas Lardon, Yuan Zhang, Jingting Chen, Luc Reymond, Peng Chen, Mai Tran, Zhongtian Yang, Haolin Zhang, Yitong Liu, Stefan Pitsch, Peng Zou, Lu Wang, Kai Johnsson, Zhixing Chen

## Abstract

Rhodamines have been continuously optimized in brightness, biocompatibility, and colors to fulfill the demands of modern bioimaging. However, the problem of phototoxicity caused by the excited fluorophore under long-term illumination has been largely neglected, hampering their use in time-lapse imaging. Here we introduce cyclooctatetraene (COT) conjugated rhodamines that span the visible spectrum and exhibit significantly reduced phototoxicity. We identified a general strategy for the generation of Gentle Rhodamines, which preserved their outstanding spectroscopic properties and cell permeability while showing an efficient reduction of singlet-oxygen formation and diminished cellular photodamage. Paradoxically, their photobleaching kinetics do not go hand in hand with reduced phototoxicity. By combining COT-conjugated spirocyclization motifs with targeting moieties, these gentle rhodamines compose a toolkit for time-lapse imaging of mitochondria, DNA, and actin and synergize with covalent and exchangeable HaloTag labeling of cellular proteins with less photodamage than their commonly used precursors. Taken together, the Gentle Rhodamines generally offer alleviated phototoxicity and allow advanced video recording applications, including voltage imaging.

## Introduction

Modern fluorescence microscopy has evolved from 3D imaging of fixed specimens to 4D-recording of subcellular structures or dynamic cellular processes in live cells or animals. Herein, a spatial resolution beyond the diffraction barrier and a temporal resolution of video rates can be achieved^1-3^. However, time-lapse recording at high resolution subjects the live samples to significantly elevated light doses, surpassing orders of magnitude the levels employed in typical one-shot wide-field or confocal imaging experiments.^4^ High excitation light exposure of fluorescent labels is known to compromise the physiological integrity of biological samples^5-7^. This phenomenon, referred to as phototoxicity, is the reversible or irreversible damaging effect of light and fluorophores on living cells or organisms. Phototoxicity mainly originates from reactive oxygen species (ROS), which are generated by the excited states of chromophores^6-8^. A major ROS relevant to phototoxicity in live-cell fluorescence microscopy is singlet oxygen, which is the product derived from the reaction of the excited fluorophore and molecular oxygen^9-11^. The reactive singlet oxygen can oxidize nearby biomacromolecules such as lipids, carbohydrates, and nucleic acids, thereby affecting their physiological functions^11-15^. Such harmful effects accumulate over time and result in abnormal cell metabolism, deformation of organelles and organisms, arrested cell proliferation, and apoptosis^7,16-18^. Therefore, phototoxicity is a universal phenomenon that widely affects live-cell fluorescence imaging practice, rendering it potentially invasive. With the democratization of super-resolution imaging and time-lapse imaging instruments, minimizing phototoxicity is of growing importance to endorse the physiological relevance of the recorded data in bioimaging.

From the perspective of photophysical chemistry, phototoxicity, and photobleaching are generally believed to stem from the excited triplet states (Fig. 1a). Triplet state quenchers (TSQs), such as mercaptoethylamine (MEA) or cyclooctatetraene (COT), have a rich history of serving as protective agents in live-cell fluorescence imaging^11,19-23^. In the past decade, the direct conjugation of such TSQ moiety on a selected dye scaffold has been proposed as a strategy for increasing photostability^19,20^, particularly demonstrated with single-molecule imaging using cyanine dyes. Recently, our laboratory and others repurposed these photophysically sophisticated molecules for live-cell super-resolution imaging of mitochondria and voltage imaging^11,21-23^, where phototoxicity has emerged as a complementary threat to photobleaching. These pioneering works are niche demonstrations tailored for a small number of cellular structures using cyanine or fluorescein dyes, whose charged chemical nature limits their general biological applications.

**Figure 1.**
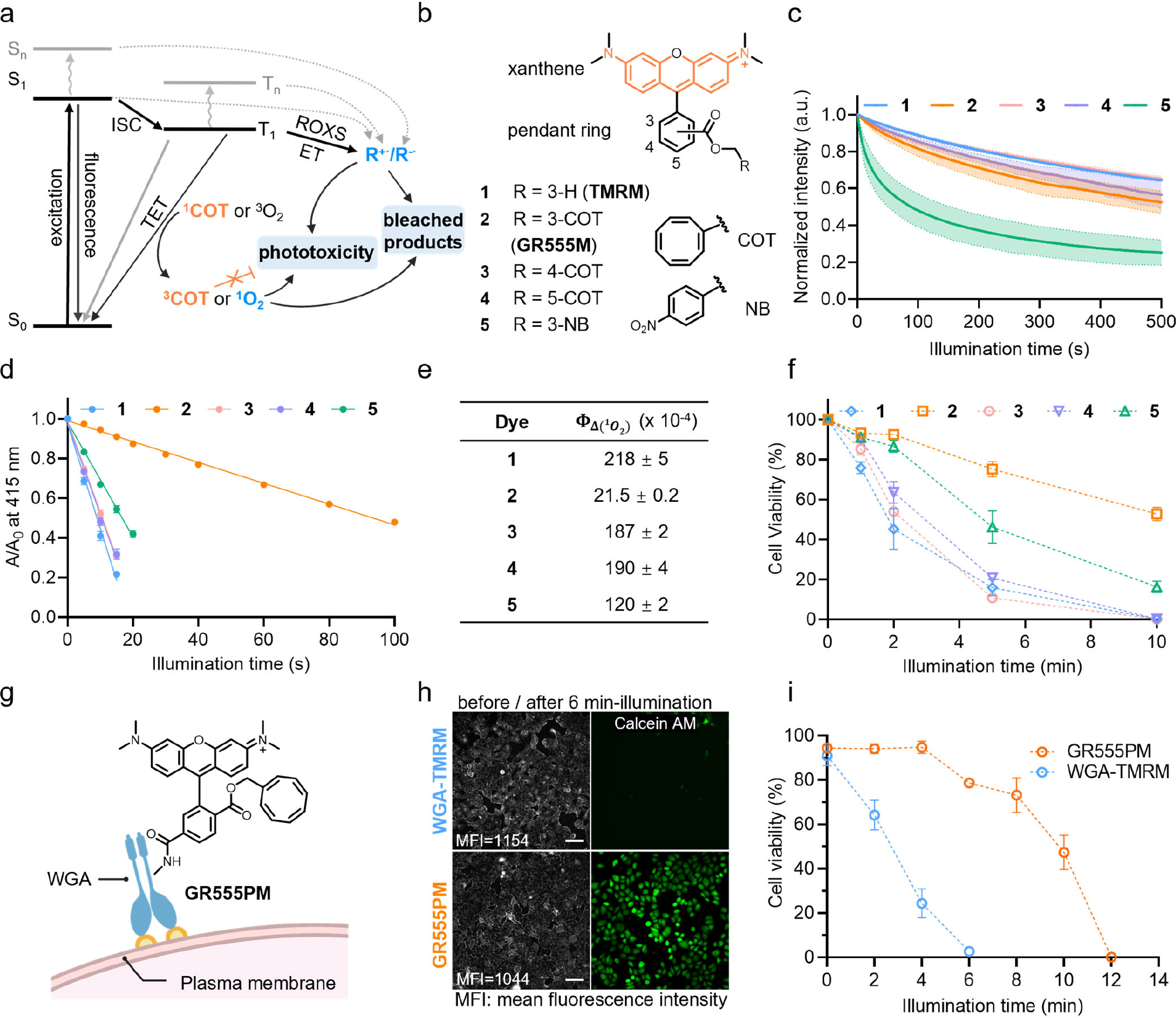
Derivatizing TMR with TSQ alleviates phototoxicity. **a**. Jablonski diagram depicting different ways of relaxation from the exited state S_1_ including photobleaching, phototoxicity, and triplet-state quenching. ISC: Intersystem crossing; TET: Triplet-energy transfer; ROXS: reducing and oxidizing system; ET: Energy transfer. b. Chemical structures of tetramethyl rhodamine-TSQs (TMR-TSQs) probes. **c**. In-vitro photobleaching experiments of compound **1-5**. Normalized photobleaching curves of 2 μM **1-5** embedded in hydrophobic polymethyl methacrylate (PMMA) film under a confocal microscopy. Data points represent averaged fluorescence bleaching curves of three independent replicates. Error bars, showing light-shaded areas, indicate standard deviation. **d**. In-vitro singlet oxygen generation experiments of compound **1-5**. The maximum absorption of DPBF at 415 nm was measured under continuous irradiation with a 520-530 nm LED lamp in the presence of compound **1-5** (the absorbance of each dye at 525 nm= 0.20; concentrations: **1**, 2.2 μM; **2**, 2.0 μM; **3**, 2.5 μM; **4**, 4.8 μM; **5**, 5.8 μM) in air-saturated acetonitrile. TMRE was used as the standard (Φ_Δ_=0.023). Data points represent normalized and averaged DPBF bleaching curves of three independent repeats. Error bars indicate standard deviation **e**. Absolute singlet oxygen quantum yields (Φ_Δ_) of compounds **1-5**. The decay slope of DPBF shown in **1d** is positively correlated with the singlet oxygen quantum yield. Standard deviations of three independent repeats. **f**. Live-cell phototoxicity measurements of compound **1-5** (250 nM, 15 min) in HeLa cells. Cell apoptosis of > 500 cells after yellow LED light illumination (561 nm, 1.4 W/cm^2^) was examined at each time point of the three independent experiments. Error bars indicate standard deviation. **g**. Schematic representation of a low-phototoxic probe, **GR555PM**, for plasma membrane labeling. **h**.Live-cell images of HeLa cells labeled with **WGA-TMRM** (30 μg/ mL, 5 min) or **GR555-PM** (50 μg/mL, 5 min) (gray) and stained with Calcein AM (1μM, 5 min, green) after illumination of 532 nm LED (∼2.6 W/cm^2^) at different illumination time. Scale bars = 100 μm. **i**. Live-cell phototoxicity measurements of **WGA-TMRM** and **GR555PM** in HeLa cells as shown in **1h**. Cell apoptosis of > 500 cells after green LED light illumination (532 nm, 2.6 W/cm^2^) were examined at each time point of the three independent experiments. Error bars indicate standard deviation.

Rhodamine dyes featuring excellent photophysical properties, spectral tunability, and cell permeability, have dominated the field of live-cell fluorescent imaging in recent years. Particularly, silicone rhodamine-(SiR)^24^-carboxyl-based probes (SiR-actin^25^, SiR-tubulin^25^, and SiR-Hoechst^26^) possessing fluorogenicity^24,27^ due to an environmentally-sensitive equilibrium between a fluorescent zwitterion and a non-fluorescent spirolactone, helped to increase the cell-permeability and reduce the background fluorescence in live-cell imaging. In addition, a general strategy tuning this dynamic equilibrium by introducing (sulfon)amide modifications to the 3-carboxylic acid was subsequently established to create multi-color fluorogenic rhodamines for live-cell nanoscopy^28^. To meet the growing demands of fluorescence nanoscopy, the photophysical properties of dyes such as brightness^29,30^, photostability^31-34^, and blinking^35,36^ have been further engineered. Most importantly, rhodamine can be used in conjunction with a variety of labeling techniques for instance self-labeling protein tags (HaloTag^37^, SNAP-tag^38,39^, and TMP-tag^40^), tetrazine for click chemistry^41^, biomolecular ligands^25^ and specific ligands for organelles^25,26^. This synergistic development has also consolidated rhodamine as a mainstream tool for bioimaging. From the phototoxicity perspective, the systematic upgrading of rhodamines towards reduced ROS generation would lead to another breakthrough for imaging tools in the field of 4D fluorescence imaging^18^.

Here, we introduced a general TSQ-conjugation strategy to reduce the phototoxicity of rhodamine derivatives. As confirmed by *in vitro* singlet-oxygen production or protein damage assays, as well as live-cell phototoxicity studies to various sub-cellular compartments, Gentle Rhodamines (GR) are valuable tools for light-intense microscopy applications. Interestingly, the TSQ-rhodamine derivatives do not necessarily bear enhanced photostability, implying alternative photobleaching pathways that are independent of triplet state populations.^42^ This strategy is compatible with a broad range of fluorogenic rhodamine derivatives and popular live-cell labeling strategies such as self-labeling tags or ligands for subcellular structures, bringing out a practical dye palette for general and gentle imaging of mitochondria, plasma membrane, nucleus, cytoskeleton, and proteins of interest in mammalian cells. We demonstrate, that GR probes can be combined with microscopy techniques like time-resolved STED and functional imaging of the membrane potential in cardiomyocytes.

## Results

### Generation of gentle tetramethyl rhodamine through COT conjugation at the 3-carboxyl position

To systematically profile the structure-activity relationship of rhodamine-TSQ conjugates, we selected COT and nitrobenzene as representative TSQs and synthesized their rhodamine derivatives. Cyclooctatetraene-1-menthanol was conjugated to the lower pendant ring of tetramethyl rhodamine (TMR) via the 3-, 4-, or 5-carboxylic acid to yield compound **2-4** and nitrobenzene was coupled to 3-carboxy TMR as compound **5** (Fig. 1b and Supplementary Fig. S1a-e, see Method for details). It has been previously demonstrated that TSQs affect the photophysics of Cy5 in a distance-dependent manner^43^. X-ray crystallography analysis of compound **2** (**GR555M**) highlights the proximity between the xanthene chromophore and the COT moiety (Supplementary Fig. S2). Notably, the COT moiety is conformationally flexible as evident from the two components in the single crystal, further enhancing its effective collision with the chromophore. Subsequently, the TSQ-TMR conjugates, along with the reference compound TMR methyl ester (**TMRM**, compound **1**), were evaluated from three aspects: (i) photostability; (ii) singlet oxygen generation in vitro; and (iii) phototoxicity based on a cell apoptosis assay. First, we measured the photobleaching curves of the five compounds. In organic polymer films (poly methyl methacrylate and vinyl alcohol), mimicking the amphiphilic environment present in cells, compounds **1-4** show high and comparable photostability. Instead, nitrobenzene compound **5** was significantly more susceptible to photobleaching (Fig. 1c and Supplementary Fig. S1f). In aqueous PBS buffer, the COT derivatives are the least photostable likely due to poor hydrophilicity (Supplementary Fig. S3). Overall, the TSQ-conjugated tetramethyl rhodamines give unpredictable trends on photostability. Next, all TSQ-conjugated dyes exhibited reduced singlet oxygen generation (Fig. 1d, e and Table S1), as measured by the singlet oxygen-induced decay of 1,3-diphenylisobenzofuran (DPBF) under the illumination of a green LED lamp (50 mW/cm^2^, 520-530 nm).^44^ The lowest singlet oxygen quantum yield was measured with the TMR derivative bearing COT in the closest proximity (3-carboxy) to the chromophore (compound **2**, Φ_Δ_:2.1 ± 0.1 × 10^−3^), which is 6-fold reduced compared to that of **TMRM** (compound **1**, Φ_Δ_: 2.2 ± 0.4 × 10^−2^, Fig. 1e and Table S1). Finally, we stained HeLa cells with compound **1-5** (250 nM) to assess the phototoxicity through a photo-induced apoptosis assay. The positive charge of these dyes leads to a bright fluorescent signal inside mitochondria, an organelle that is vulnerable to photodamage leading to apoptosis. Illumination of the cells for different time periods was performed in a high-content imager before assessing apoptosis using propidium iodide (PI) stain. The half-lethal light dose for cells stained with compound **1** was reached after 2 min-illumination, while it required 2-5 min-illumination for the TSQ-conjugated compound **3-5** to kill 50% of the cells. Remarkably, for compound **2**, the dose was reached after 10 min-illumination, meaning that such a probe could extend the duration of time-lapse recording by about five times compared to **1** (Fig. 1f). These data corroborate our previous work on COT-conjugated cyanines^11,22^, confirming that the reducing effect of TSQs on phototoxicity can be largely independent on their effect on photostability. Also, we proved that COT is a privileged TSQ for reducing the phototoxicity of fluorophores across different scaffolds like cyanines and rhodamines. In summary, COT-conjugation at 3-carboxyl of TMR can achieve the largest reduction in phototoxicity among the screened isomers and leads to bright and specific staining of mitochondria, wherefore we named this probe **GR555M**.

Having identified **GR555M** as a gentle rhodamine dye, we coupled it to wheat germ agglutinin (WGA) via its succinimidyl ester (**S25**, Supplementary Fig. S4a) yielding **GR555PM**. WGA is a lectin that binds to sialic acid and N-acetylglucosaminyl residues located on the extracellular surface of most mammalian cells^45^ and presents an established strategy for labeling and imaging the plasma-membrane of live cells. **GR555PM** works as a bright fluorescent plasma-membrane marker on live HeLa cells (Fig. 1g, h). We monitored its cellular phototoxicity in comparison to **WGA-TMRM** in a high-content imager by assessing cell viability using Calcein AM stain after continuous imaging of the labeled membranes. Compared to **WGA-TMRM**, our COT-bearing variant **GR555PM** showed lower cellular phototoxicity by a factor of four with a half-lethal dose of 10 min-illumination (Fig. 1i). This assay confirmed the reduced phototoxicity of COT-conjugated rhodamines, at the same time presenting a practical and gentle membrane stain.

### COT-conjugation gives gentle rhodamines with diverse auxochromes at various colors

Next, we extended our design to other commonly used rhodamine derivatives with wavelengths that range from green to far red. Green-emitting Rhodamine 110 and far red-emitting SiR dyes were esterified with COT-alcohol, giving rise to two novel mitochondrial dyes, **7** (**GR510M**) and **9** (**GR650M**), that offer reduced singlet oxygen generation than their methyl-ester counterparts, **6** (**Rho123**) and **8** (**SiRM**) (Fig. 2a-e and Table S1). Compound **7** and **9** differed in their bleaching behaviors relative to their parental compounds (**6** and **8**). These differences in photobleaching rate are environmentally dependent (Supplementary Fig. S5 and S6). Corroborating our studies on tetramethyl rhodamines, no general improvement of the bleaching resistance of COT-conjugated rhodamines with different colors was evident. Moreover, we esterified JF_549_ bearing azetidine auxochromes and Rhodamine 101 bearing julolidine auxochromes with COT alcohol (Compound **11** and **13**, Fig. 2a and Supplementary Fig. S7). Both compounds exhibited drastically reduced singlet-oxygen generation (Figure 2d, e). Overall, COT-conjugation is a general approach to alleviate the phototoxicity of the state-of-the-art rhodamine palette.

**Figure 2.**
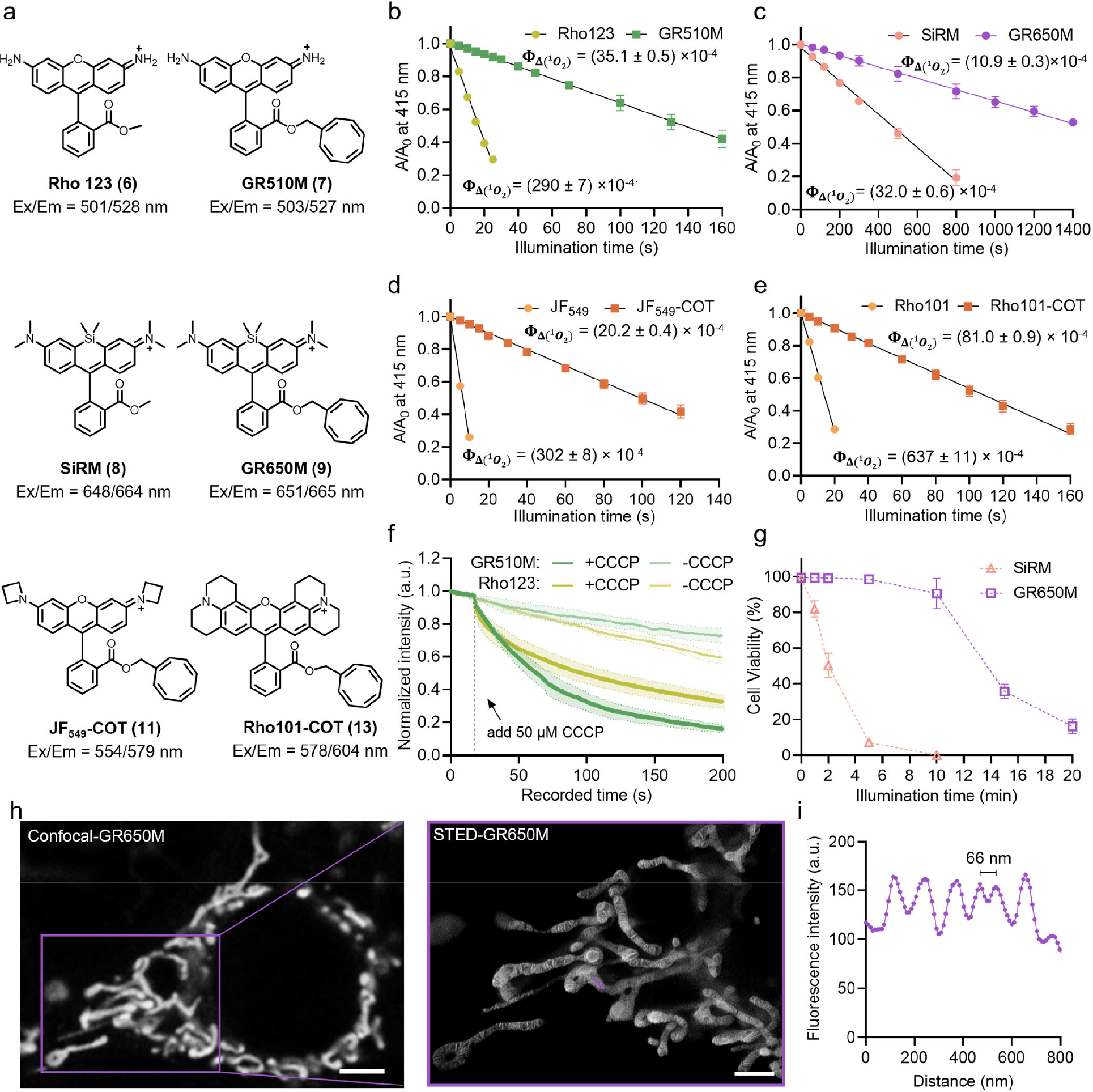
Green and far-red gentle rhodamines for live-cell fluorescence microscopy of mitochondria. **a**. Chemical structures of compounds **6-9, 11**, and **13** and their wavelengths of the maximum absorption and emission peaks. **b**. In-vitro singlet oxygen generation experiments of **Rho123** (**6**) and **GR510M** (**7**). The maximum absorption of DPBF at 415 nm was measured under continuous irradiation with an 520-530 nm LED lamp in the presence of each dye (absorbance at 525 nm= 0.20; concentrations: **6**, 2.0 μM; **7**, 2.2 μM) in air-saturated acetonitrile. The absolute singlet oxygen quantum yields (Φ_Δ_) are given on the graph. Data points represent averaged and normalized DPBF decay curves of three independent repeats. Error bars indicate standard deviation. **c**. In-vitro singlet oxygen generation experiments of **SiRM** (**8**) and **GR650M** (**9**). The maximum absorption of DPBF at 415 nm was measured under continuous irradiation with a 620-630 nm LED lamp in the presence of each dye (absorbance at 625 nm= 0.20; concentrations: **8**, 2.5 μM; **9**, 2.0 μM) in air-saturated acetonitrile. The absolute singlet oxygen quantum yields (Φ_Δ_) are given on the graph. Data points represent averaged and normalized DPBF decay curves of three independent repeats. Error bars indicate standard deviation **d**. In-vitro singlet oxygen generation experiments of **JF**_**549**_ (**10**) and **JF**_**549**_**-COT** (**11**). The maximum absorption of DPBF at 415 nm was measured under continuous irradiation with a 520-530 nm LED lamp in the presence of each dye (absorbance at 525 nm= 0.18; concentrations: **10**, 4 μM; **11**, 1 μM) in air-saturated acetonitrile. The absolute singlet oxygen quantum yields (Φ_Δ_) are given on the graph. Data points represent averaged and normalized DPBF decay curves of three independent repeats. Error bars indicate standard deviation. **e**. In-vitro singlet oxygen generation experiments of **Rho101** (**12**) and **Rho101-COT** (**13**). The maximum absorption of DPBF at 415 nm was measured under continuous irradiation with a 520-530 nm LED lamp in the presence of each dye (absorbance at 525 nm = 0.1; concentrations: **12**, 2.2 μM; **13**, 1 μM) in air-saturated acetonitrile. The absolute singlet oxygen quantum yields (Φ_Δ_) are given on the graph. Data points represent averaged and normalized DPBF decay curves of three independent repeats. Error bars indicate standard deviation. **f**. Normalized fluorescence intensities of time-lapse recordings of COS-7 cells labeled with **Rho123** or **GR510M** (300 nM, 60 min) after the addition of carbonyl cyanide 3-chlorophenylhydrazone (CCCP). Control samples were treated with **Rho123** or **GR510M** without the addition of CCCP. Data points represent averaged fluorescence intensity curves of six cells from two independent biological replicates. Error bars, showing light-shaded areas, indicate standard deviation. **g**. Phototoxicity of **GR650M** and **SiRM** (both 250 nM, 15 min) in HeLa cells, measured by cell apoptosis assay after red LED illumination (650 nm, 1.2 W/cm^2^) at different times. Data points indicate the mean of at least 1,500 individual cells from three independent biological repeats. Error bars indicate standard deviation. **h**. Confocal microscopy (left) and STED nanoscopy (right, zoom in) of live COS-7 cells labeled with **GR650M** (250 nM) for 15 min at 37 °C. Scale bar = 2 μm. (λ_ex_ = 640 nm, λ_STED_ = 775 nm) **i**. Fluorescence intensity line profiles measured as indicated in the magnified view of the purple boxed area in **h**.

We then performed the light-induced apoptosis assay with this set of gentle mitochondrial dyes. Unlike the TMR derivatives which trigger apoptosis under light-illumination, **Rho123** and **GR510M** rapidly escaped from mitochondria before the induction of apoptosis, most likely due to a higher hydrophilicity. We first verified that **GR510M** is a fast-acting mitochondrial membrane potential (MMP) indicator like **Rho123**, by recording of a rapid decrease in fluorescent intensity after the addition of the mitochondrial oxidative phosphorylation uncoupler, carbonyl cyanide 3-chlorophenylhydrazone (CCCP) (Fig. 2f and Supplementary Fig. S8), which means that the drop of **GR510M** or **Rho123** signal could indicate the MMP level and further reflect the mitochondrial health. Therefore, for **Rho123** and **GR510M**, phototoxicity was evaluated by their light-induced decrease in mitochondrial fluorescence. In continuous time-lapse recordings of stained live cells, the MMP of the cells treated with **GR510M** decreased by 50% after 1 min-illumination with a blue LED (488 nm, 1.7 W/cm^2^), whereas the control compound **Rho123** showed a more rapid decrease of MMP with a half-life of only 0.5 min (Supplementary Fig. S9). The *in-vitro* and *in-cellulo* assays collectively proved that COT-conjugation can reduce the phototoxicity of the green **R110** dye by a factor of two. Unlike the **Rho123** derivatives that dissipate from mitochondria instantaneously upon photodamage, the far-red rhodamine **GR650M** and **SiRM** exhibited a slower photo-induced leakage kinetics and induced cell death, likely due to their higher hydrophobicity. Therefore, the photo-induced apoptosis assay can be used to assess the cellular phototoxicity. The half-lethal light dose of the cells stained with **GR650M** was reached after 10-15 min-illumination with a red LED (650 nm, 1.2 W/cm^2^), which is 5-7-fold higher than that of **SiRM** (half-lethal at 2 min illumination) (Fig. 2g). These results demonstrated that the COT-conjugation reduces singlet oxygen generation and cellular phototoxicity of rhodamine derivatives with different colors.

As practical mitochondrial stains, cyanine-COT conjugates generally give stronger fluorescence signals than rhodamines. Yet our selected rhodamine-mito series complements the **PK Mito** probes in the green to yellow spectral range (**GR510M, GR555M**) and, unlike the lipophilic cyanines, **GR510M** enables instantaneous response to inner membrane potential changes for time-lapse functional imaging of mitochondria. Moreover, **GR650M** like other SiR-based probes has far-red emission and a similar quantum yield compared to its parent compound SiRM (Table S2). We demonstrate the compatibility with commercial STED nanoscopy systems equipped with a 775-nm depletion laser by the visualization of the cristae organization of COS-7 cells, which enabled us to distinguish adjacent crista at the spacing of 66 nm (Fig. 2h, i). Overall, rhodamine-COT based mitochondrial dyes supplement and supplant their cyanine counterparts (such as PK Mito dyes^11,22^) for mitochondrial recordings.

### Fluorogenic COT-rhodamines exhibit lower phototoxicity for general organelle imaging

The emerging class of fluorogenic rhodamines, bearing a dynamic equilibrium between a fluorescent zwitterion and a nonfluorescent spirolactone/lactam form, have enabled wash-free imaging of various organelles.^28^ We speculated that the COT-conjugation of rhodamines can be integrated into the spirocyclization motif, in which a COT-sulfonamide group instead of a COT-methanol is introduced to the 3-position, giving a cell-permeable and fluorogenic rhodamine core.

We first derivatized the fluorogenic TMR-COT conjugate with Hoechst at the 5-carboxyl position, yielding **GR555-DNA** (**15**) (Fig. 3a). **GR555-DNA** exhibited an 8-fold fluorescent intensity increase (“turn-on”) upon binding to hairpin-DNA (hpDNA) *in vitro*, exhibiting a higher fluorogenicity than **MaP555-DNA** and a comparable fluorescence quantum yield (Supplementary table S3). Compared to the parent compound **MaP555-DNA** (**14**), the COT-derived counterpart exhibited 8-fold lower singlet oxygen generation under LED illumination (520-530 nm, 50 mW/cm^2^, Fig. 3b and Table S1).

**Figure 3.**
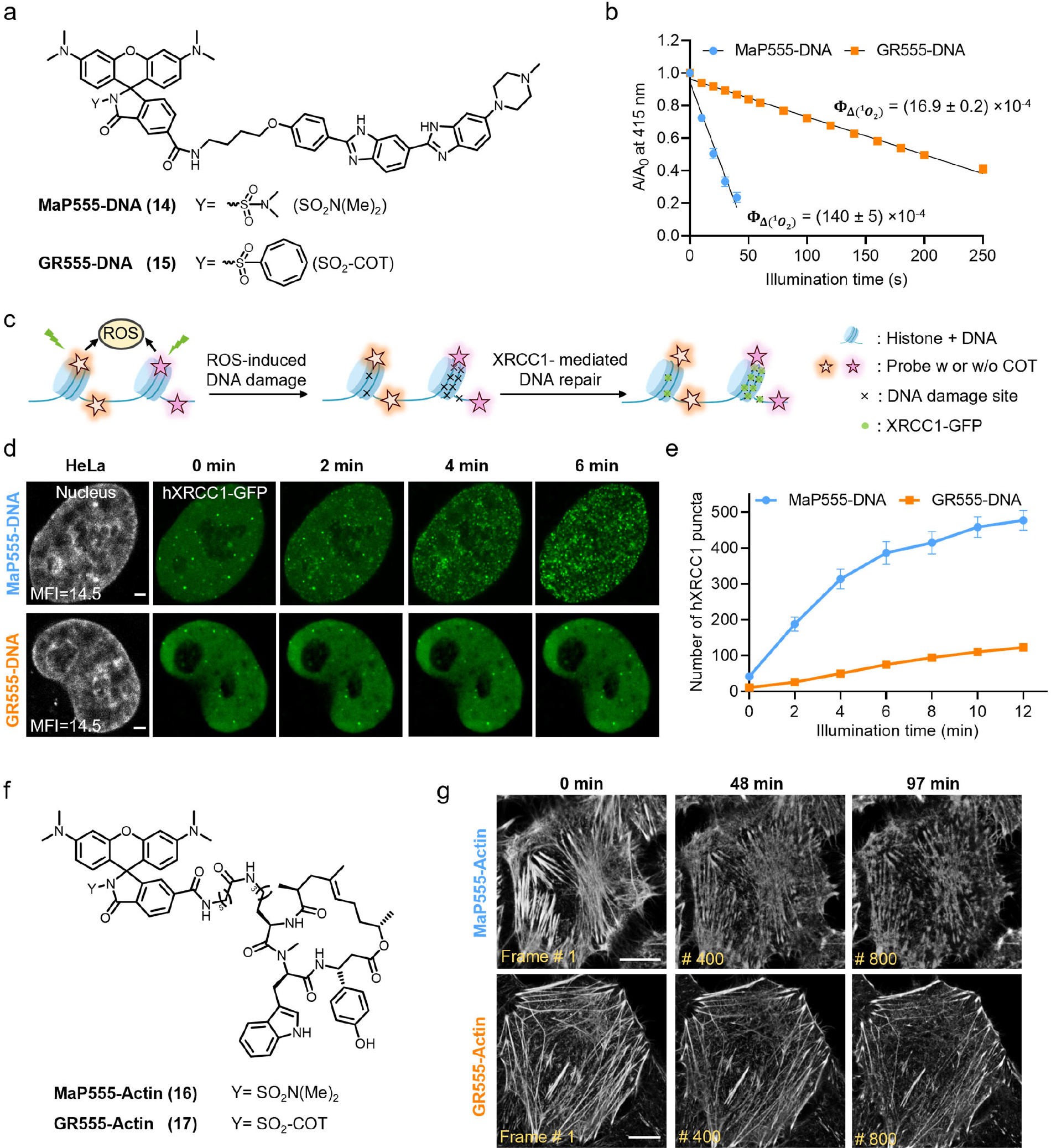
Targetable gentle rhodamines for live-cell imaging of DNA and cytoskeleton. **a**.Chemical structures of **MaP555-DNA** (**14**, blue) and **GR555-DNA** (**15**, orange). **b**. In-vitro singlet oxygen generation experiments of **MaP555-DNA** and **GR555-DNA**. The maximum absorption of DPBF at 415 nm was measured under continues irradiation with an 520-530 nm LED lamp in presence of each dye (absorbance at 525 nm = 0.15; concentrations: **14**, 1 μM; **15**, 1.1 μM) in air-saturated acetonitrile containing 0.1%TFA. The absolute singlet oxygen quantum yields (Φ_Δ_) are given on the graph. Data points represent averaged and normalized DPBF decay curves of three independent repeats. Error bars indicate standard deviation. **c**. Schematic representation of the DNA damage assay based on hXRCC1-GFP. Upon (light-induced) DNA damage, hXRCC1-GFP gets recruited to the damaged site. **d**. Live cell confocal images of HeLa cells expressing hXRCC1-GFP at a frame rate of 2 min/frame. HeLa cells were labeled with **MaP555-DNA** (gray, 200 nM) or **GR555-DNA** (gray, 2 μM) for 60 min at 37 °C. Puncta formation in the time-lapse images of DNA repairing protein hXRCC1 fused with GFP indicates the DNA damage level (green). Scale bars = 2 μm. **e**. Semi-quantitative analysis of cellular phototoxicity of **MaP555-DNA** and **GR555-DNA** of HeLa hXRCC1-GFP cells, as measured by the total number of hXRCC1-GFP puncta from experiments shown in **d**. Data points represent the averaged hXRCC1-GFP number of eleven cells from five independent experiments. Error bars indicate the standard error of the mean. **f**. Chemical structures of actin dyes **MaP555-Actin** (**16**, blue) and **GR555-Actin** (**17**, orange). **g**. Long-term time-lapse confocal recordings of HeLa cells at a frame rate of 7.27 sec/frame. HeLa cells were labeled with **MaP555-Actin** or **GR555-Actin** (both 100 nM with 10 μM verapamil) for 3 h at 37 °C. Cells labeled with **GR555-Actin** showed no shrinkage and fracture of actin filaments during the time of recording. Scale bars = 10 μm.

For live-cell staining, **GR555-DNA** showed nuclear specificity. The new DNA stain displayed a lower cell-permeability than **MaP555-DNA**, presumably due to a larger molecular weight. Yet for HeLa cells staining, labeling with 2 μM **GR555-DNA** or 0.2 μM **MaP555-DNA** for 60 min resulted in similar brightness and signal-to-noise ratios under no-wash conditions (Supplementary Fig. S10a). Notably, the staining conditions of **GR555-DNA**, although slightly more demanding, did not lead to significant cytotoxicity (Supplementary Fig. S11). We then exploited a DNA repair imaging assay to semi-quantitatively characterize the phototoxicity of DNA dyes in live cells (schematic diagram shown in Fig. 3c). X-ray cross-complementing protein 1 (XRCC1) is a scaffolding protein that accumulates at sites of DNA-damage and recruits other proteins involved in DNA repair pathways.^46^ hXRCC1-GFP is evenly distributed in the nucleus of healthy cells, while upon DNA damage it gets recruited to the damaged site and exhibits multiple fluorescent puncta patterns in the nucleus, giving a sensitive assay of DNA damage under stress.^47^ HeLa cells expressing hXRCC1-GFP labeled with **MaP555-DNA** showed a gradual increase in the number of hXRCC1-GFP puncta after 2 min-exposure to a 560 nm pulse laser, and the puncta numbers plateaued after 10 min-exposure. In contrast, HeLa cells labeled with **GR555-DNA** had a lower number of hXRCC1-GFP puncta than those treated with **MaP555-DNA** (Fig. 3d, e). To control the photobleaching factors, we also compared the *in cellulo* photostability of the **GR555-DNA** and **MaP555-DNA**. Neither of the two DNA dyes showed significant fluorescence intensity decay after 12 min of exposure to the 560 nm pulsed laser (Supplementary Fig. S10b). Therefore, we concluded that the COT-conjugation strategy reduced the cellular phototoxicity when such fluorophore is attached to a DNA-targeting moiety and ultimately minimize the DNA damage in live-cell microscopy.

Next, we aimed to develop a low-phototoxicity probe for imaging of the cytoskeleton in live cells. We conjugated **GR555** to a Jasplakinolide derivative binding to F-actin^25^ yielding **GR555-Actin** (compound **17**, Fig. 3f). **GR555-Actin** has a ∼6-fold reduced singlet oxygen generation compared to the reference dye **MaP555-Actin** (compound **16**, Supplementary Fig. S12 and Table S1). In long-term time-lapse confocal imaging, **GR555-Actin** exhibited reduced phototoxicity and enhanced brightness and photostability (Supplementary Fig. S13). Actin filaments of HeLa cells stained with **MaP555-Actin** tend to shrink, accompanied by a large attenuation of the fluorescence intensity, and gradually disintegrated and fractured into short strands after 41-47 min (350-400 frames). In comparison, **GR555-Actin** enabled acquisitions of 118 min (1000 frames) with integral actin filament structures (Fig. 3g and Supplementary Movie S1). Together, these data demonstrate that the COT-conjugated actin probe enables long-term imaging of cellular structures with low phototoxicity.

### Fluorogenic and gentle COT-rhodamines for HaloTag

Self-labeling protein (SLP) tags enable chemogenetic labeling of specific cellular proteins with cell-permeable fluorophores. Compared to the fluorescent proteins (FPs), SLPs offer the mean to select between bright fluorescent probes of different colors and spectroscopic properties, making them the prime method for live-cell nanoscopy. However, the dyes attached to the tags are exposed to the cellular environment and therefore potentially phototoxic. In contrast, the chromophores of FPs are shielded to insulate the sensitization process in ROS generation^48^. To address the phototoxicity of SLP substrates, we coupled our gentle spirocyclic rhodamine dye to the chloroalkane HaloTag Ligand (HTL) to obtain **GR555-HTL** (**19**) (Fig. 4a). In addition, to broaden the spectral range and explore a different fluorophore scaffold, we synthesized the corresponding red-shifted carborhodamine derivative **GR618-HTL** (**21**), as an analogue to its previously published parent dye **MaP618-HTL** (**20**)^28^ (Fig. 4a).

**Figure 4.**
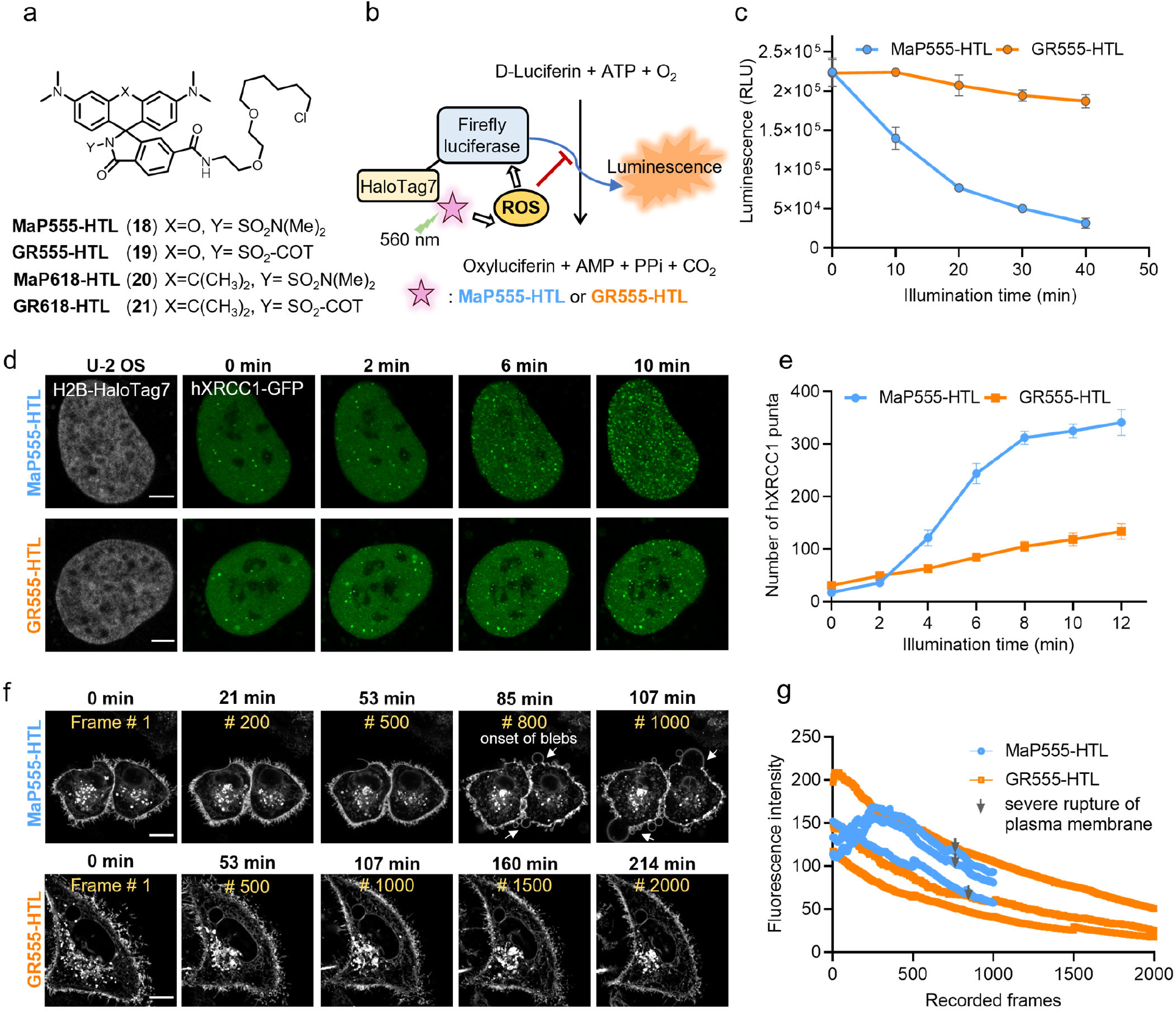
Gentle rhodamines for live-cell imaging of cellular proteins using self-labeling protein tags. **a**.Chemical structures of **MaP555/618, GR555/618** (**18** – **21**) derivatives coupled to the HaloTag Ligand (chloroalkane substrate). **b**. Schematic diagram of the protein damage assay. Firefly luciferase-HaloTag7 (FLuc-HaloTag7) is labeled with **MaP555-HTL** and **GR555-HTL** and the photo-damage under long-term illumination assessed by the luminescence generated by FLuc afterwards. **c**. Light-induced photodamage of FLuc-HaloTag7 labeled with **MaP555-HTL** and **GR555-HTL** *in-vitro*. The fully labeled protein was illuminated and after different time points the protein damage was assessed by D-Luciferin addition and luminescence measurements. Data points represent the averaged luminescence of three independent experiments. Error bars indicate the standard error of the mean. **d**. Live-cell confocal recordings (gray) of U-2 OS cells expressing H2B-HaloTag7 (stable) and DNA repairing protein hXRCC1-GFP (transient) at a frame rate of 2 min/frame. U-2 OS cells were labeled with **MaP555-HTL** or **GR555-HTL** (both 500 nM) for 30 min at 37 °C. Scale bars: 5 μm. **e**. Semi-quantitative analysis of cellular phototoxicity analysis of **GR555-HTL** and **MaP555-HTL** of U-2 OS H2B-HaloTag7 cells, as measured by counting the total number of hXRCC1-GFP puncta. Data points represent the averaged hXRCC1-GFP number of five cells from five independent experiments. Error bars indicate the standard error of the mean. **f**. Long-term time-lapse confocal recordings of HeLa cells expressing HaloTag7-PDGFR^tmb^ at a frame rate of 6.41 sec/frame. HeLa cells were labeled with **MaP555-HTL** or **GR555-HTL** (both 500 nM) for 30 min at 37 °C. Cells labeled with **GR555-HTL** showed no appearance of blebs and intact plasma membrane for the time of recording. Scale bars = 10 μm. **g**. Photobleaching curves of HeLa cells expressing HaloTag7-PDGFR^tmb^ labeled with **MaP555-HTL** or **GR555-HTL** under continuous time-lapse confocal recordings using a 561 nm pulsed laser. The gray arrows indicate the onset of blebbing and membrane disruption. Each curve represents the bleaching curve of an individual HeLa cell.

We compared the propensity of both probes to exist in the spirocyclic form by water-dioxane titrations: **GR555-HTL** probe is less fluorogenic (D_50_: 40) than **MaP555-HTL** (**18**) (D_50_: 66), yet it displays a mild (∼4-fold) fluorescence turn-on and similar brightness when bound to HaloTag7 (Supplementary table S3 and Fig. S14a, b). **GR618-HTL** displayed a great fluorogenic potential (D_50_ > 75) and yielded ∼41-fold fluorescence turn-on upon HaloTag7 labeling (Supplementary Table S3 and Fig. S14c, d), which makes it attractive for no-wash, live-cell applications. Although *in-vitro* and live-cell labeling kinetics showed that **GR618-HTL** labeled HaloTag protein slower than **MaP618-HTL** (Supplementary Fig. S15), GR-HTL probes display equivalent signal brightness and fluorescence life-times in live-cell labeling experiments compared to their established analogues without the COT moiety (Supplementary Fig. S16, Table S3). The DPBF assay measuring singlet oxygen generation demonstrated that both **GR555-HTL** and **GR618-HTL** featured lower singlet oxygen generation (by 10-/4-fold respectively) than their MaP counterparts (Supplementary Fig. S17 and Table S1).

We then designed an *in-vitro* assay to assess the ROS damage to a functional protein that is fused to HaloTag7 (Fig. 4b). Firefly luciferase (FLuc) HaloTag7-fusion protein was labeled with **GR555-HTL** or **MaP555-HTL** (Supplementary Fig. S18a-c). After green light excitation (LED 520-530 nm, 50 mW/cm^2^) for different periods (0 – 40 min), D-Luciferin was added and the luciferase activity was measured using a bioluminescence assay. FLuc-HaloTag7 labeled with **MaP555-HTL** exhibited a severe drop (> 85%) of enzymatic activity during the 40-min illumination, indicating that the ROS generated from **MaP555-HTL** is profoundly damaging the FLuc. In contrast, FLuc-HaloTag7 labeled with **GR555-HTL** showed only a drop of < 20% after 40 min of illumination. Of note, the remaining fluorescence signal after 40 min illumination of protein samples was > 60%, in which **GR555-HaloTag** was slightly more photostable than **MaP555-HaloTag** (Fig. 4c and Supplementary Fig. S18d). Therefore, we conclude that the lower phototoxicity attributed to **GR555-HTL** indeed prevents damage to closeby proteins as compared to regular **MaP555-HTL** labeling.

To further validate the reduced phototoxicity of **GR555-HTL** at the cellular level, we fused HaloTag7 to the nuclear histone 2B (H2B). H2B is the major protein component of chromatin with a high abundance inside the nucleus. Moreover, H2B acting as the spool of DNA winding featured an extremely close spatial distance with DNA, which allowed us to assess the cellular phototoxicity of **GR555-HTL** using the XRCC1 assay (Fig. 3c). We first stained H2B-HaloTag7-expressing U-2 OS cells **GR555-HTL** or **MaP555-HTL** (500 nM each), resulting in fluorescent histone with comparable brightness (Supplementary Fig. S19). To compare the phototoxicity, we next transfected the same cells with hXRCC1-GFP and imaged its accumulation during continuous exposure to a 560 nm laser. For **MaP555-HTL**-treated samples, a uniform distribution of hXRCC1-GFP with few puncta switched to a dense scattered distribution, with a maximum puncta number of about 300-400, within 10 min of intense laser exposure. In contrast, the cells labeled with **GR555-HTL** experienced a slow increase in hXRCC1 puncta with a maximum puncta number of ∼100 (Fig. 4d, e), supporting that COT conjugation significantly reduces cellular photodamage (by 3-to 4-fold) on H2B-HaloTag7 under long-term illumination. Similar results were obtained for **GR618-HTL** (Supplementary Fig. S20). Next, we targeted HaloTag7 to the outer plasma membrane by C-terminal fusion to the transmembrane domain of the platelet-derived growth factor (PDGFR^tmb^, pDisplay sequence) and stained mEGFP-HaloTag7-PDGFR^tmb^ expressing HeLa cells with 500 nM **MaP555-HTL** or **GR555-HTL**, reaching a similar brightness on the plasma membrane (Fig. 4f, g). During time-lapse confocal recordings, severe rupture of plasma membrane structures was recorded in cells labeled with **MaP555-HTL** after 85 min (800 frames), as characterized by the formation of blebs. In contrast, **GR555-HTL**-labeled cells did not undergo significant apoptosis during up to 214 min (2000 frames) of time-lapse imaging (Fig. 4f, g and Supplementary Movie S2). Also, more dynamic behaviors of the plasma membrane were observed and less apoptosis arose when employing Gentle Rhodamine HaloTag7 probes than traditional probes for long-term imaging (Fig. 4f and Supplementary Fig. S21).

The red dye **GR618-HTL** opens new possibilities for spectral multiplexing and super-resolution imaging with low phototoxicity and excellent photostability. First, it can be combined with orange probes like **GR555-DNA** and near-infra-red probes like **GR650M** for multi-color imaging. We combined those probes to stain primary rat hippocampal neurons expressing CalR-HaloTag7-KDEL and recorded a 4 h (150 frames) confocal time-lapsed video (Supplementary Fig. S22 and Movie S3) with few signs of phototoxicity. Under similar conditions, the established probes **MaP555-DNA, MaP618-HTL**, and **SiRM** showed a loss of mitochondrial integrity (reduction of SiRM signal) after 2 h (70 frames). Second, it can be used on commercial STED systems having a 775 nm depletion laser. To highlight the capability of this approach, we combined **GR618-HTL** and **GR650M** and recorded dual-color STED images of mitochondria and the endoplasmic reticulum (CalR-HaloTag7-KDEL) in live neurons (Fig. 5a). To further boost the photostability, we synergistically combined the GR strategy with exchangeable HaloTag7 ligands (xHTLs)^49^. **GR618** was conjugated to an xHTL linker, giving rise to the non-covalent probes **GR618-S5** (**22**) (Supplementary Fig. S23a). The photobleaching behaviors of covalent (**MaP/GR618-HTL**) and the exchangeable HaloTag substrates (**MaP/GR618-S5**) were profiled using time-lapsed STED nanoscopy on U-2 OS cells expressing TOM20-HaloTag7 (Fig. 5b). Here, **GR618-S5** exhibits a slower photobleaching rate compared to **GR618-HTL**, suggesting the compatibility of **GR618** with xHTLs which gives approximately 5-fold enhancement in photostability (Fig. 5c). This finding was confirmed with a second xHTL (Hy5, Supplementary Fig. S23). Notably, the photobleaching profiles of **MaP618** and **GR618** are largely the same. This data, along with the similar photobleaching profiles of **GR555 and MaP555** (Fig. 4g), suggests that the COT conjugation alone is not able to the photostability of HaloTag-labeled rhodamines.

**Figure 5.**
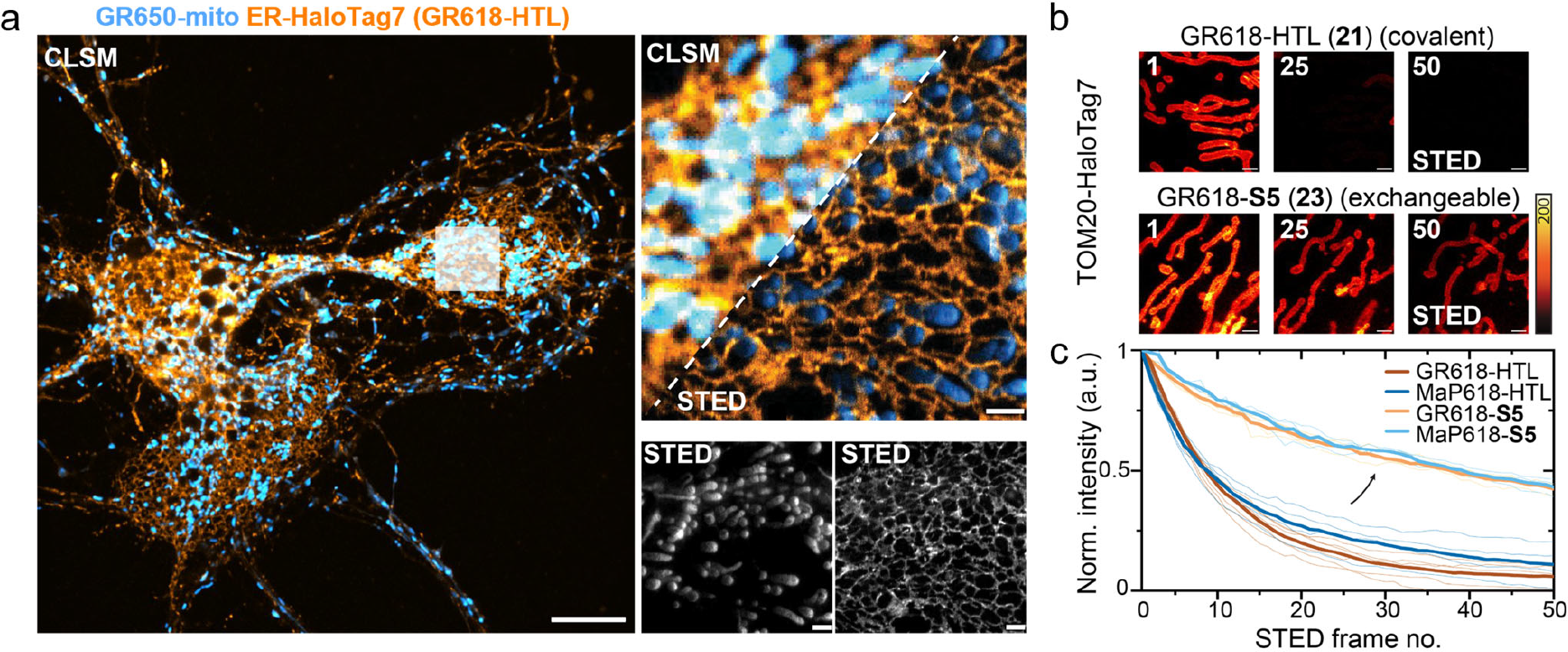
Gentle rhodamines are privileged dyes for long-term multi-color STED nanoscopy recordings. **a**.Dual-color confocal laser-scanning microscopy (CLSM) and STED imaging of the ER and mitochondria using gentle rhodamine probes in live cultured rat hippocampal neurons. Neurons (10 DIV) expressing CalR-HaloTag7 (rAAV transduction) were stained with **GR618-HTL** (cyan, 500 nM) and **GR650M** (red, 50 nM) for 30 min at 37 °C. **GR618-HTL** was excited with a 561 nm laser and **GR650M** with a 640 nm laser. Both dyes were depleted with a 775 nm depletion laser (STED). White rectangle in the CLSM overview (right) shows magnified FOV for STED imaging. Scale bars = 10 μm (overview), 2 μm (magnification).**b**. Time-lapse STED imaging showcasing different photobleaching behavior of **MaP/GR618** covalently conjugated to HaloTag (HTL) or its exchangeable counterpart (S5). Multi-frame STED imaging of U-2 OS mitochondria outer membrane (TOM20-HaloTag7) labeled with **GR618-(x) HTLs** over 50 consecutive frames in a 10 × 10 μm ROI using **MaP618/GR618-HTL, -S5**. Frame numbers are indicated in the top left corner. Scale bars: 1 μm. **c**. Bleaching curves (thick lines: mean value, thin lines: individual experiments) plotted for at least 4 image series (n ≥ 4) as shown in **b**.

Finally, we showcase long-time-lapse functional imaging on primary cells by combining **GR-HTL** probes with a chemigenetic voltage indicator. Voltron consists of genetically encoded Ace2 rhodopsin fused to HaloTag7 and a fluorescent HaloTag ligand.^50^ Compared to FRET-based indicators employing fluorescent proteins, Voltron offers a brighter signal and a larger dynamic range^51^. However, use of Voltron can result in phototoxicity, hampering voltage recordings (Fig. 6a). We labeled neonatal rat cardiomyocytes (NRCMs) expressing Voltron with **GR555-HTL** or **MaP555-HTL** respectively (Fig. 6b) and monitored their activity by recording the changes in fluorescence (△F/F_0_) to trace their spontaneous electrical signals. However, after 156 ± 25 seconds of continuous imaging at 100 Hz, the cardiomyocytes labeled with **MaP555-HTL** stopped beating and firing due to accumulated phototoxicity (Fig. 6c, d). In contrast, cardiomyocytes labeled with **GR555-HTL** provided a continuous voltage signal for up to 573 ± 64 seconds under identical imaging conditions (561 nm laser illumination at 2.16 W/cm^2^) before the firing stopped (Fig. 6c, e). Therefore, gentle rhodamines combined with self-labeling protein tags offer superior tools for the physiological time-lapsed studies of primary cells.

**Figure 6.**
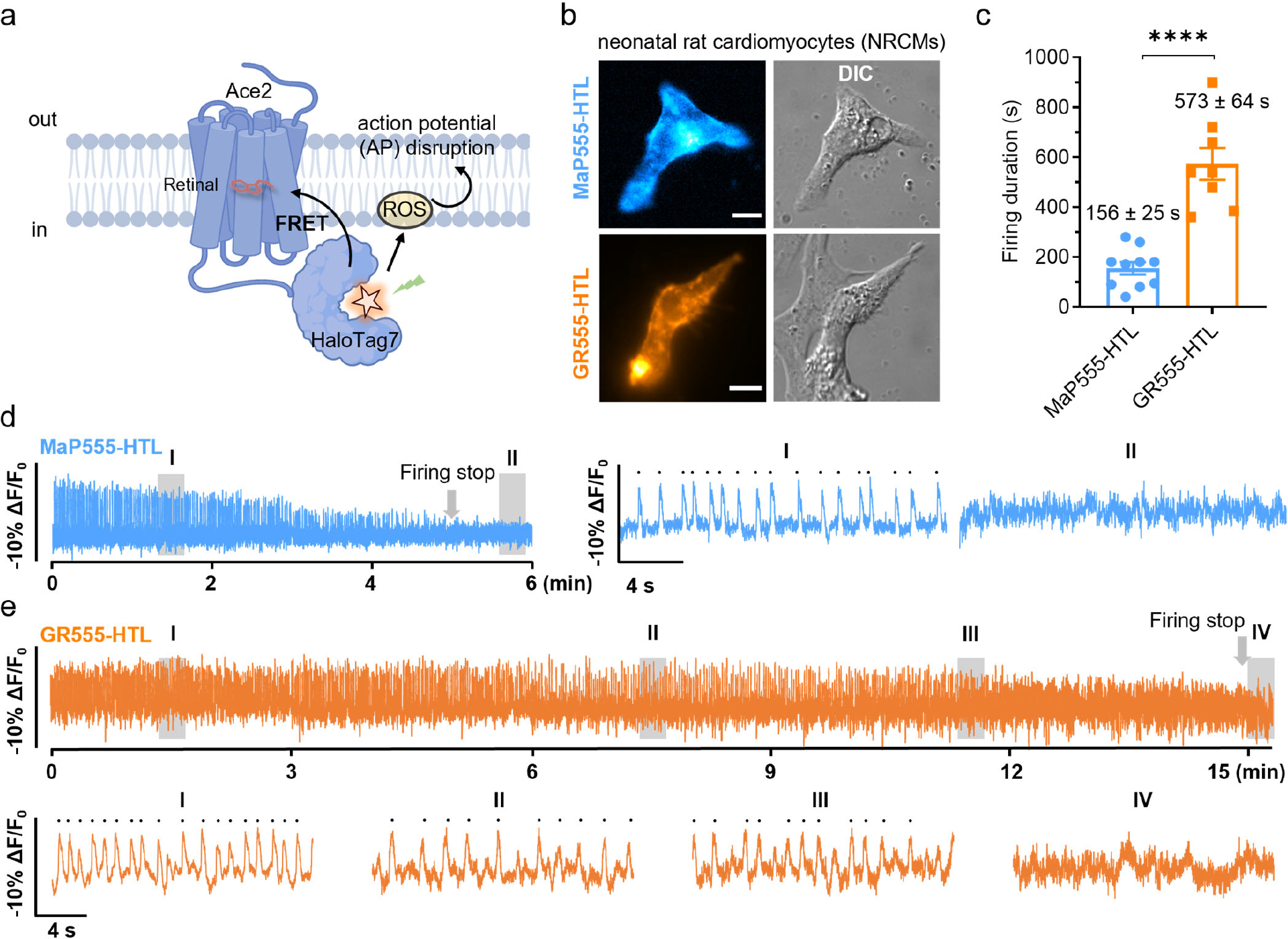
Gentle rhodamines are privileged dyes for voltage imaging of primary cells. **a**. Schematic representation of phototoxicity during long-term voltage imaging based on chemigenetic voltage-indicator Voltron. **b**. Wide-field microscopy of the neonatal rat cardiomyocytes (NRCMs) expressing Voltron (Ace2-HaloTag7) and labeled with **MaP555-HTL** or **GR555-HTL** (both 100 nM) for 25 min at 37 °C. Scale bar = 10 μm. **c**. Firing duration of NRCMs expressing Voltron (Ace2-HaloTag7) labeled with **MaP555-HTL** or **GR555-HTL**. The illumination intensities of 561 nm lasers were 2.16 W·cm^-2^. Bars indicate the mean of seven cells. Error bars indicate the standard error of the mean. Significance was determined using a two-tailed unpaired t-test followed by Sidak’s multiple comparisons test. *P=* **** < 1.0 ×10^−4^. **d. & e**. A representative fluorescence trace of NRCMs expressing Voltron and labeled with **MaP555-HTL** or **GR555-HTL**. Each peak on the traces showed spontaneous spikes of each NRCM and signals were corrected for photobleaching. The illumination intensity of the 561 nm laser was 2.16 W/cm^2^. **f**. 6 min (36,000 frames) recordings at 100 frames/sec were performed. Two zoomed-in signals (i-ii) from two shaded regions (I-II) were presented at the right. Each black dot represents one spontaneous spike. **g**. 15 min (90,000 frames) recordings at 100 frames/sec were performed. Four zoomed-in signals (i-iv) from four shaded regions (I-IV) were presented at the bottom. Each black dot represents one spontaneous spike.

## Discussion and Conclusion

In this work, we demonstrate that rhodamine dyes, the privileged toolkit for live cell imaging, can be rendered less phototoxic upon the conjugation of COT at the proximal 3-carboxyl group. Contrary to our expectations, the rhodamine-COT conjugates are generally less phototoxic than parental rhodamines but exhibit unpredictable behaviors in photobleaching (Fig. 1c, 4g and Supplementary Fig. S1f, S3, S5c-d, S6c-d, S13b, and S18d). These results prompt us to reconsider the intriguing relationship between phototoxicity and photostability. Unlike Cy5 chromophore whose photobleaching is mainly attributed to cycloaddition with the triplet-state-sensitized singlet oxygen^52^, we speculate that the main photobleaching pathways of xanthenes may not stem from the excited triplet state^32^. In addition, while rhodamines emitting orange to far-red synergize well with COT conjugation, the yellow emitting **GR510M** bears a decreased fluorescence quantum yield (Table S2), corroborating a recent theoretical study on the impact of COT to the singlet states of blue dyes^53^. Collectively, this work offers insight into the elusive photophysics of rhodamine dyes, and establishes COT-conjugation as a general strategy for alleviating phototoxicity, if not as general a method for enhancing photostability.

With the increasing demand for spatial and temporal resolutions in live cell imaging, we argue that phototoxicity in live-cell imaging is a fundamental challenge of growing importance^5,7^. In this work, photodamage was thoroughly assessed through assays ranging from *in-vitro* ROS generation, proximity protein damage, cell death, and morphological and physiological alterations. We have now established TSQ conjugation as a primary approach to systematically alleviate phototoxicity^11,22^ while minimizing the non-specific binding of dyes is another viable direction in parallel^54^. In the future, we will further assess and optimize the tissue permeability of gentle rhodamines towards *in vivo* applications. As rhodamines are modular building blocks that can be readily combined with state-of-the-art labeling technologies, the gentle rhodamines reported here thus represent chemical solutions to phototoxicity issues in live-cell imaging. These chemical approaches would eventually synergize with mathematical, optical, and spectroscopical approaches^1,55^ to enable time-lapse dynamic imaging, offering long-lasting fluorescence signals that transfer into multiplexed spatial and temporal information with uncompromised physiological relevance.

## Acknowledgments

This project was supported by funds from the Beijing Municipal Science & Technology Commission (Project: Z221100003422013 to Z.C.). We thank Prof. Yulong Li for the gift of the plasmid CMV-HaloTag7-pDisplay, Prof. Wulan Deng, and Ying Bi for the U-2 OS H2B-HaloTag7 cells, Drs. Jie Su and Yuanhe Li for single crystal data analysis, Dr. Shuzhang Liu for support on voltage imaging and data analysis, A. Bergner, D. Ginkel, A. Herold (MPIMR) for providing materials and reagents. We thank the analytical instrumentation center of Peking University, the NMR facility and optical imaging facility of the National Center for Protein Sciences at Peking University, and the MS facility of MPIMR for assistance with data acquisition.

## Data availability

All data reported in this paper will be shared by the corresponding author upon reasonable request.

## Author contributions

L.W., K. J., and Y. Z., and Z. C., independently conceived the concept and the three teams combined their efforts. T. L., J. K., and J. L. designed, performed, and analyzed the biological assays. J. C., and Z. Y., contributed to the phototoxicity and imaging experiments. T. L., J. K., N. L., Y. Z., L. R., P. C., M. T., Z. Y., H. Z., Y. L., and S.P. performed the chemical synthesis and characterizations. P. Z. supervised the voltage imaging assay. K. J. and Z.C. supervised the project. T.L., J.K., K.J., and Z.C. wrote the paper.

## Reference

1 Huang, X. et al. Fast, long-term, super-resolution imaging with Hessian structured illumination microscopy. Nat. Biotechnol. 36, 451–459 (2018).

2 Gwosch, K. C. et al. MINFLUX nanoscopy delivers 3D multicolor nanometer resolution in cells. Nat. Methods 17, 217–224 (2020).

3 Westphal, V. et al. Video-Rate Far-Field Optical Nanoscopy Dissects Synaptic Vesicle Movement. Science 320, 246–249 (2008).

4 Tosheva, K. L., Yuan, Y., Matos Pereira, P., Culley, S. & Henriques, R. Between life and death: strategies to reduce phototoxicity in super-resolution microscopy. J. Phys. D 53, 163001(2020).

5 Kilian, N. et al. Assessing photodamage in live-cell STED microscopy. Nat. Methods 15, 755–756 (2018).

6 Daddysman, M. K., Tycon, M. A. & Fecko, C. J. in Photoswitching Proteins: Methods and Protocols (ed Sidney Cambridge) 1–17 (Springer New York, 2014).

7 Laissue, P. P., Alghamdi, R. A., Tomancak, P., Reynaud, E. G. & Shroff, H. Assessing phototoxicity in live fluorescence imaging. Nat. Methods 14, 657–661 (2017).

8 Dixit, R. & Cyr, R. Cell damage and reactive oxygen species production induced by fluorescence microscopy: effect on mitosis and guidelines for non-invasive fluorescence microscopy. Plant J. 36, 280–290 (2003).

9 Dahl, T. A., Robert Midden, W. & Hartman, P. E. Pure exogenous singlet oxygen: Nonmutagenicity in bacteria. Mutat. Res.-Fund. Mol. M. 201, 127–136 (1988).

10 Cadenas, E. biochemistry of oxygen toxicity. Annu. Rev. Biochem. 58, 79–110 (1989).

11 Yang, Z. et al. Cyclooctatetraene-conjugated cyanine mitochondrial probes minimize phototoxicity in fluorescence and nanoscopic imaging. Chem. Sci. 11, 8506–8516 (2020).

12 Redmond, R. W. & Kochevar, I. E. Spatially resolved cellular responses to singlet oxygen. Photochem. Photobiol. 82, 1178–1186 (2006).

13 Agostinis, P. et al. Photodynamic therapy of cancer: An update. CA Cancer J. Clin. 61, 250–281 (2011).

14 Wojtovich, A. P. & Foster, T. H. Optogenetic control of ROS production. Redox. Biol. 2, 368–376 (2014).

15 Srinivas, U. S., Tan, B. W. Q., Vellayappan, B. A. & Jeyasekharan, A. D. ROS and the DNA damage response in cancer. Redox. Biol. 25, 101084 (2019).

16 Wäldchen, S., Lehmann, J., Klein, T., van de Linde, S. & Sauer, M. Light-induced cell damage in live-cell super-resolution microscopy. Sci. Rep. 5, 15348 (2015).

17 Magidson, V. & Khodjakov, A. Circumventing photodamage in live-cell microscopy. Methods Cell Biol. 114, 545–560 (2013).

18 Rajendraprasad, G., Rodriguez-Calado, S. & Barisic, M. SiR-DNA/SiR–Hoechst-induced chromosome entanglement generates severe anaphase bridges and DNA damage. Life Science Alliance 6, e202302260 (2023).

19 Altman, R. B. et al. Cyanine fluorophore derivatives with enhanced photostability. Nat. Methods 9, 68–71 (2012).

20 van der Velde, J. H. M. et al. A simple and versatile design concept for fluorophore derivatives with intramolecular photostabilization. Nat. Commun. 7, 10144 (2016).

21 Grenier, V. et al. Molecular Prosthetics for Long-Term Functional Imaging with Fluorescent Reporters. ACS Cent. Sci. 8, 118–121 (2022).

22 Liu, T. et al. Multi-color live-cell STED nanoscopy of mitochondria with a gentle inner membrane stain. Proc. Natl. Acad. Sci. U.S.A. 119, e2215799119 (2022).

23 Liu, S. et al. Orange/far-red hybrid voltage indicators with reduced phototoxicity enable reliable long-term imaging in neurons and cardiomyocytes. Proc. Natl. Acad. Sci. U.S.A. 120, e2306950120 (2023).

24 Lukinavičius, G. et al. A near-infrared fluorophore for live-cell super-resolution microscopy of cellular proteins. Nat. Chem. 5, 132–139 (2013).

25 Lukinavičius, G. et al. Fluorogenic probes for live-cell imaging of the cytoskeleton. Nat. Methods 11, 731–733 (2014).

26 Lukinavičius, G. et al. SiR–Hoechst is a far-red DNA stain for live-cell nanoscopy. Nat. Commun. 6, 8497 (2015).

27 Wang, L., Frei, M. S., Salim, A. & Johnsson, K. Small-Molecule Fluorescent Probes for Live-Cell Super-Resolution Microscopy. J. Am. Chem. Soc. 141, 2770–2781 (2019).

28 Wang, L. et al. A general strategy to develop cell permeable and fluorogenic probes for multicolour nanoscopy. Nat. Chem. 12, 165–172 (2020).

29 Grimm, J. B. et al. A general method to improve fluorophores for live-cell and single-molecule microscopy. Nat. Methods 12, 244–250 (2015).

30 Grimm, J. B. et al. A general method to fine-tune fluorophores for live-cell and in vivo imaging. Nat. Methods 14, 987–994 (2017).

31 Kolmakov, K. et al. A Versatile Route to Red-Emitting Carbopyronine Dyes for Optical Microscopy and Nanoscopy. Eur. J. Org. Chem. 2010, 3593–3610 (2010).

32 Butkevich, A. N., Bossi, M. L., Lukinavičius, G. & Hell, S. W. Triarylmethane Fluorophores Resistant to Oxidative Photobluing. J. Am. Chem. Soc. 141, 981–989 (2019).

33 Grimm, J. B. et al. A General Method to Improve Fluorophores Using Deuterated Auxochromes. JACS Au 1, 690–696 (2021).

34 Roßmann, K. et al. N-Methyl deuterated rhodamines for protein labelling in sensitive fluorescence microscopy. Chem. Sci. 13, 8605–8617 (2022).

35 Uno, S.-n. et al. A spontaneously blinking fluorophore based on intramolecular spirocyclization for live-cell super-resolution imaging. Nat. Chem. 6, 681–689 (2014).

36 Lardon, N. et al. Systematic Tuning of Rhodamine Spirocyclization for Super-resolution Microscopy. J. Am. Chem. Soc. 143, 14592–14600 (2021).

37 Los, G. V. et al. HaloTag: A Novel Protein Labeling Technology for Cell Imaging and Protein Analysis. ACS Chem. Biol. 3, 373–382 (2008).

38 Keppler, A. et al. A general method for the covalent labeling of fusion proteins with small molecules in vivo. Nat. Biotechnol. 21, 86–89 (2003).

39 Hein, B. et al. Stimulated Emission Depletion Nanoscopy of Living Cells Using SNAP-Tag Fusion Proteins. Biophys. J. 98, 158–163 (2010).

40 Mo, J. et al. Third-Generation Covalent TMP-Tag for Fast Labeling and Multiplexed Imaging of Cellular Proteins. Angew. Chem. Int. Ed. 61, e202207905 (2022).

41 Werther, P. et al. Bio-orthogonal Red and Far-Red Fluorogenic Probes for Wash-Free Live-Cell and Super-resolution Microscopy. ACS Cent. Sci. 7, 1561–1571 (2021).

42 Butkevich, A. N., Bossi, M. L., Lukinavičius, G. & Hell, S. W. Triarylmethane Fluorophores Resistant to Oxidative Photobluing. J. Am. Chem. Soc 141, 981–989 (2019).

43 Zheng, Q. et al. Electronic tuning of self-healing fluorophores for live-cell and single-molecule imaging. Chem. Sci. 8, 755–762 (2017).

44 Entradas, T., Waldron, S. & Volk, M. The detection sensitivity of commonly used singlet oxygen probes in aqueous environments. J. Photochem. Photobiol. B, Biol. 204, 111787 (2020).

45 Monsigny, M., Roche, A.-C., Sene, C., Maget-Dana, R. & Delmotte, F. Sugar-Lectin Interactions: How Does Wheat-Germ Agglutinin Bind Sialoglycoconjugates? Eur. J. Biochem. 104, 147–153 (1980).

46 Caldecott, K. W. XRCC1 protein; Form and function. DNA Repair 81, 102664 (2019).

47 Ryumina, A. P. et al. Flavoprotein miniSOG as a genetically encoded photosensitizer for cancer cells. Biochim. Biophys. Acta. Gen. Subj. 1830, 5059–5067 (2013).

48 Surrey, T. et al. Chromophore-assisted light inactivation and self-organization of microtubules and motors. Proc. Natl. Acad. Sci. U.S.A. 95, 4293–4298 (1998).

49 Kompa, J. et al. Exchangeable HaloTag Ligands for Super-Resolution Fluorescence Microscopy. J. Am. Chem. Soc. 145, 3075–3083 (2023).

50 Abdelfattah, A. S. et al. Bright and photostable chemigenetic indicators for extended in vivo voltage imaging. Science 365, 699–704 (2019).

51 Vogt, N. A bright future for voltage imaging. Nat. Methods 16, 1076–1076 (2019).

52 Gorka, A. P., Nani, R. R. & Schnermann, M. J. Cyanine polyene reactivity: scope and biomedical applications. Org. Biomol. Chem. 13, 7584–7598 (2015).

53 Chanmungkalakul, S., Abedi, S. A. A., Hernández, F. J., Xu, J. & Liu, X. The dark side of cyclooctatetraene (COT): Photophysics in the singlet states of “self-healing” dyes. Chin. Chem. Lett., 109227 (2023).

54 Zhang, J. et al. Red- and Far-Red-Emitting Zinc Probes with Minimal Phototoxicity for Multiplexed Recording of Orchestrated Insulin Secretion. Angew. Chem. Int. Ed. 60, 25846–25855 (2021).

55 Ludvikova, L. et al. Near-infrared co-illumination of fluorescent proteins reduces photobleaching and phototoxicity. Nature Biotechnology, doi:10.1038/s41587-023-01893-7 (2023).

