## Supplementary Information for "Gentle rhodamines for live-cell fluorescence microscopy"

1. College of Future Technology, Institute of Molecular Medicine, National Biomedical Imaging Center, Beijing Key Laboratory of Cardiometabolic Molecular Medicine, Peking University, Beijing 100871, China
2. Peking-Tsinghua Center for Life Science, Academy for Advanced Interdisciplinary Studies, Peking University, Beijing 100871, China
3. Department of Chemical Biology, Max Planck Institute for Medical Research, Heidelberg 69120, Germany;
4. Biomolecular Screening Facility, École Polytechnique Fédérale de Lausanne (EPFL), Lausanne 1015, Switzerland
5. PKU-Nanjing Institute of Translational Medicine, Nanjing 211800, China
6. GenVivo Tech, Nanjing 211800, China
7. Spirochrome AG, Chalberwiedstrasse 4, CH-8260 Stein am Rhein, Switzerland.
8. College of Chemistry and Molecular Engineering, Synthetic and Functional Biomolecules Center, Beijing National Laboratory for Molecular Sciences, Key Laboratory of Bioorganic Chemistry and Molecular Engineering of the Ministry of Education, PKU-IDG/McGovern Institute for Brain Research, Peking University, Beijing 100871, China
9. Key Laboratory of Smart Drug Delivery, Ministry of Education, School of Pharmacy, Fudan University, 201203 Shanghai, China;
10. Equal contribution

**This PDF file includes:**

Supplementary information text
- Legends for Movies S1 to S3
- Material and Methods
- Figures S1 to S23
- Table S1-S12

Chemical synthesis and characterization of new compounds

SI References

**Other supplementary materials for this manuscript include the following:**

Movies S1 to S3

**Legends for Movies S1 to S3**

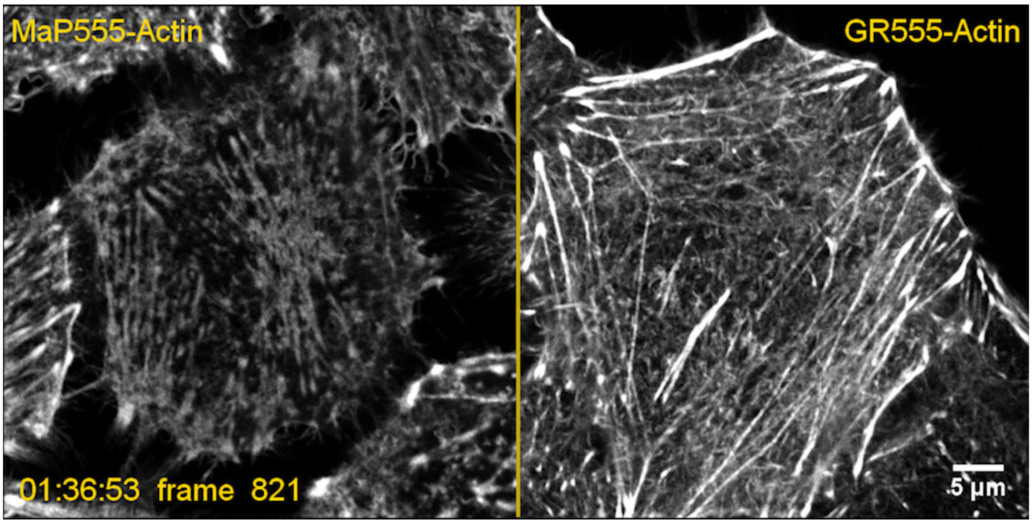


**Movie S1.** Long-term time-lapse confocal recordings of HeLa cells labeled with **MaP555-Actin** or **GR555-Actin**. Scale bar = 5 µm.

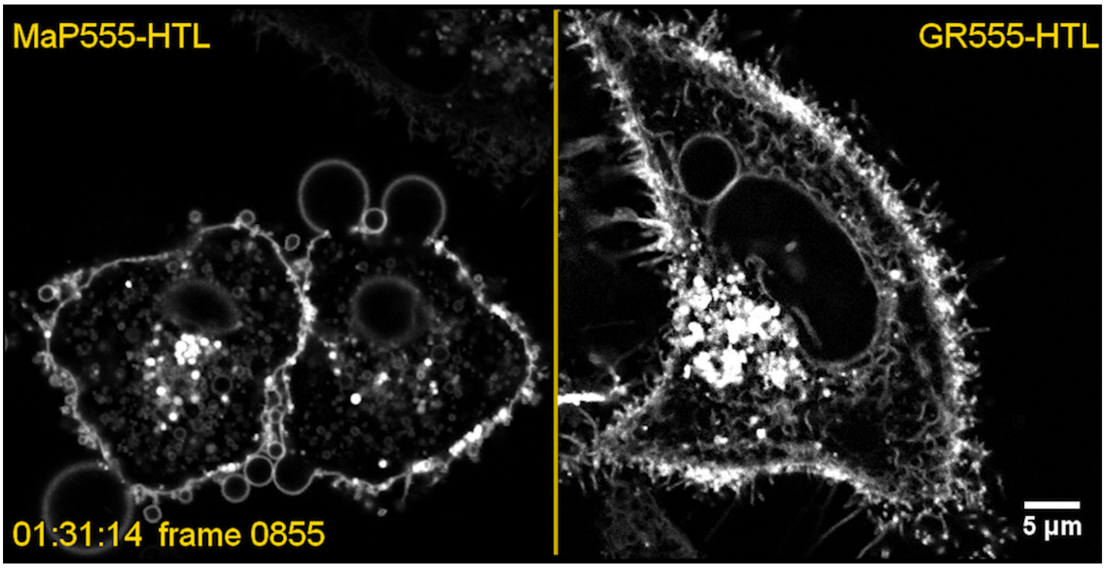


**Movie S2.** Long-term time-lapse confocal recordings of HeLa HaloTag7-PDGFR$^{tmb}$ cells labeled with **MaP555-HTL** or **GR555-HTL**. Scale bar = 5 µm.

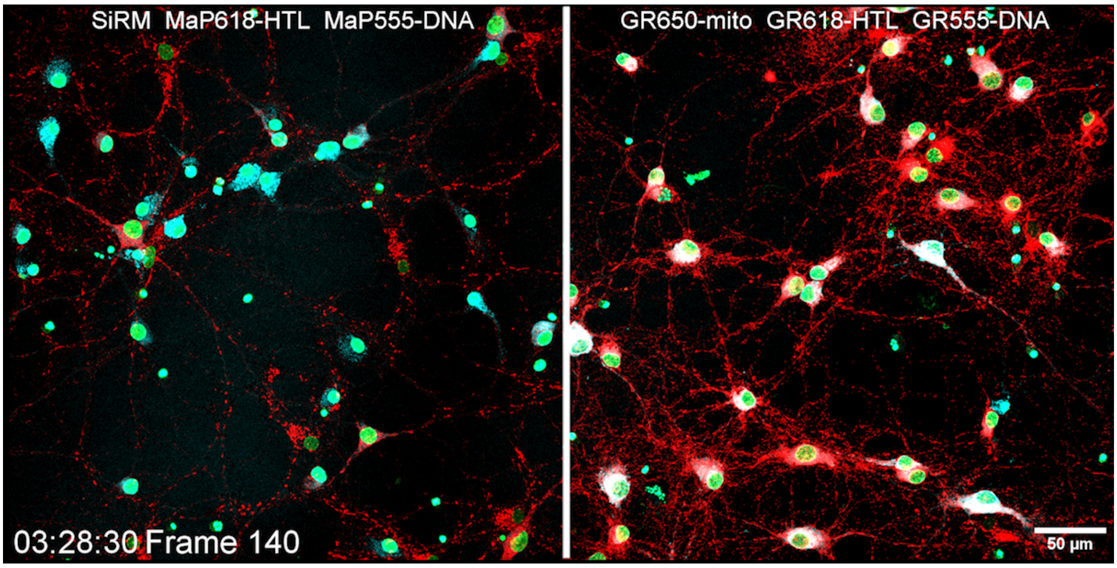


**Movie S3.** Three-color time-lapse confocal recordings of mitochondria, endoplasmic reticulum, and DNA on rat hippocampal neurons expressing CalR-HaloTag7-KDEL. Confocal recordings of 12 z-stacks per channel at a frame rate of 90 sec/frame were performed for 4 h. Scale bar = 50 µm.

### Supplementary Information Text

#### Material and Methods

**Bulk bleaching measurement**

On polymer films: Compounds **1-9** were immobilized within polyvinyl alcohol (PVA) and polymethylmethacrylate (PMMA) films by using a SW-4A spin coater (Setcas, Beijing, China) and round glass coverslips (diameter = 25 mm). For PVA films, 100 µL 5% PVA (S27770-500g; Yuanye, Shanghai, China) in Mili-Q water containing 2 µM dyes were applied to the coverslip followed by spin coating (80,000 rpm, 10 sec; 300,000 rpm, 45 sec); for PMMA films, 100 µL 5% PMMA in chloroform (112048; Tongguang, Beijing, China) containing 2 µM dyes were applied to coverslip followed by spin coating (80,000 rpm 10 sec; 800,000 rpm 45 sec). These polymer films were then irradiated using a STEDYCON confocal microscope (Abberior Instruments GmbH, Göttingen, Germany) equipped with a 20x Air CFI Super Fluor/0.75 NA objective (Nikon, Tokyo, Japan). Three groups of time-lapse images (500 frames, 50 µm x 150 µm, 1 fps, 100% laser power) were acquired for each dye, and quantification of the fluorescence intensity was achieved with the Fiji software[1] from three parallel experiments.

In phosphate buffers: Compound **1-5** were diluted with phosphate buffer (PBS, pH=7.4) to 10 µM and followed by irradiation with LED lamp (520-530 nm, 50 mW/cm$^2$). UV-Vis absorption spectra were measured on a microplate reader (TECAN Infinite M Nano+, Switzerland) in 96-well plates.

**Measurement of singlet oxygen quantum yield**

Singlet oxygen quantum yields of compounds **1-13** and methylene blue (MB) were determined using 1,3-Diphenylisobenzofuran (DPBF, Q105708; Dibai, Shanghai, China) as a chemical indicator. 1-5 µM dyes and 8 x 10$^{-5}$ M DPBF were dissolved in air-saturated acetonitrile (ACN) or methanol (MeOH), followed by irradiation with an LED lamp (520-530 nm, 50 mW/cm$^2$ for compound **1-7** and**10-13**; 620-630 nm, 0.5 mW/cm$^2$ for compound **8-9** and MB). Singlet oxygen quantum yields of compound **14-21** were determined using DPBF decay assay in air-saturated ACN containing 0.1% TFA followed by irradiation with an LED lamp (520-530 nm, 50 mW/cm$^2$ for compound **14-19**; 620-630 nm, 50 mW/cm$^2$ for compound **20-21**). The absorbance values at 525 nm or 625 nm were corrected for the dye concentration. DPBF absorption was monitored at 415 nm. The linear decay slope of 415 nm is positively correlated with singlet oxygen quantum yield. **TMRE** ($\varphi = 0.023$) in acetonitrile , **Rho123** (**6**) ($\varphi = 0.03$) in methanol, and **MB** ($\varphi = 0.49$) in ethanol were used as references for rhodamines in different colors[2].

**Protein production and purification**

E. coli BL21(DE3) (ZC121; Zoman Biotechnology, Beijing, China) or BL21(DE3)-pLysS (Novagen, Sigma Aldrich) were transformed with plasmid DNA. Single colonies were cultured in lysogeny broth (LB) medium containing 1% ampicillin at 37 °C until the $OD_{600}$ reached 0.6-1.0. Isopropyl β-D-Thiogalactoside (IPTG) was then added for inducing protein expression at a final concentration of 1 mM.

Firefly luciferase (FLuc)-HaloTag7 (T7-6×His vector) was produced for 5 h at 37 °C, bacteria were collected by centrifugation at 5,000g (15 min, 4 °C), the pellet was resuspended in PBS (pH 7.4) solution added with 1 mM phenylmethylsulfonyl fluoride (PMSF) and then lysed by sonification (SCINENTZ JY192-IIN, Ningbo, China; 30% power, "3s-on" / "6s-off", 10 circles). The total protein was harvested from the supernatant collected after centrifugation (12,000g, 30 min) and performed following labeling assay and luciferase activity assay.

HaloTag7 (pET-10xHis-TEV vector) was produced for 20 h at 16 °C, bacteria were collected by centrifugation at 4,000 g (15 min, 4 °C), the pellet was resuspended in 50 mM $KH_2PO_4$ buffer with 300 mM NaCl, 5 mM imidazole (pH 8.0) supplemented with 1 mM phenylmethylsulfonyl fluoride (PMSF) and 0.25 mg/mL lysozyme. The cells were lysed by sonification (SONOPLUS, 7 min, 50% on/off cycles, 70% amplitude) and the debris was removed by centrifugation (20 min, 50,000 g, 4 °C). The supernatant was charged on a HisTrap FF crude column (Cytiva, Marlborough, MA) and purified by immobilized metal ion affinity chromatography (IMAC) using a ÄktaPure FPLC instrument (Cytiva). Afterwards the protein was concentrated using Ultra 15 mL Centrifugal Filters (Amicon®, MWKO: 10 kDa, Merck KGaA, Darmstadt, Germany) and the buffer was switched to 50 mM HEPES buffer with 50 mM NaCl (pH 7.3) by washing trice.

**In-vitro protein damage assay**

Firefly Luciferase (Fluc)-HaloTag7 (crude total protein concentration: 0.85 mM, based on average molecular mass of protein in E.coli: 40 KDa[3]) was incubated with excessive **MaP555-HTL**, **GR555-HTL** or **SiR-HTL** (24 μM in PBS) at r.t. in the dark and shaken for 2 h. Fluc-HaloTag7 was first labeled with **MaP555-HTL** or **GR555-HTL** respectively in-vitro, and then labeled with **SiR-HTL** again, as a pulse-chase experiment, for the measurement of labeling efficiency of **MaP555-HTL** or **GR555-HTL**. After labeling, the reaction mix was passed through Zeba Spin Desalting Columns (7K MWCO, 89882; Thermo Fisher Scientific, Waltham, MA, USA) to remove excess dyes. The desalted proteins were analyzed by SDS-PAGE to obtain the labeling efficiency of **MaP555-HTL** or **GR555-HTL**. The labeling efficiency (%) was calculated according to the following equation: cell viability % = (A-B) / A *100%, where A = the fluorescence intensity of band in Cy3 channel, and B = the fluorescence intensity of band in Cy5 channel. The labeling efficiency of **SiR-HTL** was assumed to be 100%. For protein damage assay, the labeled protein was transferred to a transparent 384-well plate (3656; Corning, NY, USA) and illuminated with a green LED lamp (520-530 nm, 50 mW/cm$^2$) for different periods. After illumination, the samples were transferred to a white bottom 384-well plate (3572; Corning), the FLuc substrate (100x D-Luciferin from Dual Luciferase Reporter Gene Assay Kit, KGAF0400; KeyGEN BioTECH, Nanjing, China) was added and the FLuc activity was measured by luminescence using a fluorophotometer RF-5301 (Shimadzu, Tokyo, Japan) at 1s integration time.

**Measurements of UV-Vis, fluorescence spectroscopy and quantum yields**

Dyes for spectroscopic characterizations were prepared as stock solutions (1 – 10 mM) in DMSO and diluted such that the DMSO concentration did not exceed 0.5% (v/v). The concentration of compound **S20** (**GR555-COOH**) and compound **S22** (**GR618-COOH**) was determined by qNMR using a dioxane standard (5 mM). Spectral characterization experiments were carried out in 50 mM NaCl, 50 mM HEPES buffer (pH 7.4) containing 1% (w/v) BSA. HaloTag7 labeling was performed for 2 h at room-temperature using the protein in excess.

UV-Vis and fluorescence spectroscopy

Compound **1-9** was diluted with acetonitrile (ACN) or phosphate buffer saline (PBS) to 1 μM. UV-vis absorption spectra were measured on a microplate reader (TECAN Infinite M Nano+, Switzerland) in 96-well plates. Fluorescence emission spectra were measured using a Shimadzu RF-5301PC spectrofluorometer (Shimadzu) in a 1 cm square quartz cuvette.

Fluorescence increase upon protein labeling assay

**GR-HTLs** (50 nM) were prepared in buffer in presence and absence of HaloTag7 (15 μM). GR-DNA probes (50 nM) were prepared in presence and absence of hpDNA (50 μM) following the protocol of G. Lukinavičius *et al*[5]. Fluorescence emission scans were recorded in a black flat bottom 384 well plate (Greiner) by exciting the fluorophore and measuring the fluorescence emission intensity on a microplate reader (Spark20M, Tecan). The fluorescence intensity at the emission maxima (**Table S3**) was averaged over three technical replicates and the fluorescence increase was obtained as the ratio of fluorescence intensity in presence and absence of the HaloTag7 protein (ΔF).

Extinction coefficients measurement

A dilution series (0.5 – 6 μM) of **GR555-/618-HTL** in buffer, 0.1% SDS or HaloTag7 (15 μM) was prepared. Absorbance spectra were recorded in clear bottom non-binding 96 well plates (Greiner) on a plate reader (Spark20M, Tecan, 400 to 700 nm, 2 nm step size). Technical triplicates were averaged and the data was baseline corrected. The maximum absorbance (**Table S3**) was plotted against the concentration and the data fitted to a linear function. The path length (200 μL volume) was obtained by performing control experiments using **MaP555-** and **MaP618-HTL** in 0.1% SDS for which the extinction coefficient was previously reported[4]. The extinction coefficients were calculated using the Lambert-Beer's law.

Quantum yield measurement

**GR555-/618-HTL** (1 μM) were prepared in absence and presence of HaloTag7 (15 μM). **MaP555-/GR555-DNA** (1 μM) was incubated in presence of hpDNA as previously reported.[5] Quantum yields were measured in Glass Crimp Neck Vial (8 mm, Fisherbrand™, Fisher Scientific) from three individual replicates (n = 3) by exciting the fluorophore at its maximum absorbance (**Table S3**) in an absolute quantum yield spectrometer (Quantaurs-QY, model C11347, Hamatsu, Hamamatsu City, Japan).

**Water-Dioxane Titration**

**GR555-/618-HTL** (5 µM) were diluted in 10/90 to 80/20 (v/v) water-1,4-dioxane mixtures (dry, Acros). Absorbance spectra were recorded in transparent flat-bottom polypropylene 96-well plates (Greiner, Chimney Well) on a plate reader (Spark20M, Tecan, 400 to 700 nm, 2 nm step size). Technical triplicates were averaged and the data was baseline corrected. The maximum absorbance (**Table S3**) was normalized to the max. and min. absorbance and plotted against the dielectric constants $\varepsilon_R$ corresponding to the water/dioxane ratio[6]. $D_{50}$-values were determined as the dielectric constant at half-maximal absorbance by fitting the data as previously reported[7].

**In-vitro HaloTag7 labeling kinetics measurement**

**MaP618/-GR618-HTL** (1 µM) was prepared in buffer (50 mM HEPES, 50 mM NaCl, 0.5% BSA, pH 7.3) in a black flat bottom 384 well plate (Greiner). The fluorescence intensity was measured by exciting the fluorophore (600±10 nm) and measuring the fluorescence emission intensity (626±10 nm) on a microplate reader (Spark20M, Tecan) containing a humidity cassette at 37 °C. 100 µM HaloTag7 were added and the fluorescence emission increase was measured every 15 s in technical triplicates, averaged and normalized to the initial fluorescence signal. The dead-time was 12s and the fluorescence intensity in absence of HaloTag7 was tracked as well.

**Plasmids**

Molecular cloning was performed by Gibson assembly as described previously[9] to combine the insert fragments and vectors after PCR. Detailed protocols on molecular cloning were referenced to previous work[10]. All plasmids used in this study are listed in **Table S4**. To generate pRetroQ-hXRCC1-AcGFP, the plasmid pRetroQ-lifeact-AcGFP (a gift from Prof. Jincai Luo) and the custom-synthesized hXRCC1 gene (Genewiz, Suzhou, China) were amplified using primer (a) - (d). T7-6×His-firefly luciferase-HaloTag7 was generated from T7-6×His-firefly luciferase (a gift from Prof. Lei Chen) and HaloTag7 amplified from the plasmid pCMV-TOM20-HaloTag7[11] using primer (e) - (h). The plasmid (for stable cell lines) PB-pCMV-mEGFP-HaloTag7-PDGFR$^{tmb}$ was generated from pCMV-mEGFP-HaloTag7-PDGFR$^{tmb}$ (a gift from Prof. Yulong Li) and PiggyBac (PB) backbone PB-PB-PGK-Puro-PL30 (a gift from Prof. Wulan Deng) using primer (i)-(l). To generate pCMV-Voltron (Ace2-HaloTag7), the fragment Voltron was first constructed by point-mutation PCR using primer (m) - (r). The plasmid pCMV-CheRiff-eGFP (a gift from Prof. Peng Zou) and Voltron gene were amplified using primer (s) - (v).

(a) 5'-CGAATTCTGCAGTCGACGGT-3'
(b) 5'-CATGGTCTCGAGATCTGAGTCCG-3'
(c) 5'-ACTCAGATCTCGAGACCATGCCTGAAATTAGACTGAGACACGTGGT-3'
(d) 5'-ACCGTCGACTGCAGAATTCGGGCCTGGGGCACCAC-3'
(e) 5'-GATTAGCGGCTAACAAAGCCCGAAAGGAAGCTG-3'
(f) 5'-ATAGAGCCGCCGCCGCCCACGGCGATCTTGCCGC--3'
(g) 5'-GGCGGCGGCTCTATGGGCAGCGAAATTGGCA-3'
(h) 5'-CGGGCTTTGTTAGCCGCTAATCTCCAGTGTACTCAG-3'
(i) 5'-ACATTGATTATTGACTAGTTATTAATAGTAATCAATTACGGG-3'
(j) 5'-CGACCCAGCGGCCCTTAACGTGGCTTCTTCTGCCA-3'
(k) 5'-TAAGGGCCGCTGGGTCG-3'
(l) 5'-TCAATAATCAATGTTATATCTGGCCCGTACATATCGCG-3'
(m) 5'-TACGCAAGATATATTAACTGGGTGCTGACC-3'
(n) 5'-TGTGGTCAGCACCCAGTTAATATATCTTGC-3'
(o) 5'-CCACTGCTCCTGCTCGATCTCATCGTCATG-3'
(p) 5'-GGTCATGACGATGAGATCGAGCAGGAGCAG-3'
(q) 5'-CTGTCCGTCGACAATGAGGCCATTCTCATG-3'
(r) 5'-TCCCATGAGAATGGCCTCATTGTCGACGGA-3'
(s) 5'-TAAGGCGGCCGCGACTC-3'

(t) 5’-GGTGGCGGATCCCGGTG-3’
(u) 5’-CACCGGGATCCGCCACCATGGCTGACGTGGAAACCGAG-3’
(v) 5’-GAGTCGCGGCCGCCTTAGAATCGAGTAGCCTCGGGGAAATC-3’

**Cell culture**
HeLa (MB5422-S; HARVEYBIO, Beijing, China), U-2 OS EF1α H2B-HaloTag7-expressing cells (gift from Prof. Wulan Deng) and stable U-2 OS Flp-IN T-REx™ (ThermoFischer Scientifics) cell-lines were cultured in a high-glucose Dulbecco´s modified Eagle´s medium (DMEM) (Gibco, 11965092; Thermo Fisher Scientific) containing 10% (v/v) heat-inactivated fetal bovine serum (SE100-011; Vistech, Sydney, Australia). In some cases, 1% (v/v) penicillin sulfate and streptomycin (CC004; Macgene, Beijing, China) were supplemented.

**Preparation of neuron cultures**
Primary rat neurons were isolated from new born pups (0-1 days, WISTAR rats). Procedures were performed in accordance with the Animal Welfare Act of the Federal Republic of Germany (Tierschutzgesetz der Bundesrepublik Deutschland, TierSchG) and the Animal Welfare Laboratory Animal Regulations (Tierschutzversuchsverordnung). According to the TierSchG and the Tierschutzversuchsverordnung, no ethical approval from the ethics committee is required for the procedure of euthanizing rodents for subsequent extraction of tissues. The procedure for euthanizing rats performed in this study was supervised by animal welfare officers of the Max Planck Institute for Medical Research (MPImF) and conducted and documented according to the guidelines of the TierSchG (permit number assigned by the MPImF: MPI/T-35/18).
After sacrificing, the hippocampi were isolated, digest with trypsin and mechanically dissected using a pipette to obtain a homogenous solution. The solution was filtered through a cell strainer (40 µm pore size) and transferred to pre-coated (poly-L-ornithine 100 µg/mL, laminin 1 µg/mL in 1x HBSS) µ-Slide 8-well glass-bottom plates (80826; ibidi, Gräfelfing, Germany). The medium was replaced 2 h after seeding with fresh phenol-red free 27 neurobasal medium (NB) supplemented with Penicilin/Streptomycin (Life Technologies), GlutaMAX and B27 was added. Hippocampal neurons were maintained in a humidified incubator at 37°C and 5% $CO_2$ atmosphere.

**Preparation of cardiomyocyte**
Neonatal rat cardiomyocytes were isolated from Sprague-Dawley rats born within 24 h. The whole hearts were isolated, minced and rinsed in Tyrode's buffer. And then, six cycles of digestion using collagenase type II (0.08% w/v, Worthington Biochemical, New Jersey, USA) and pancreatin (0.1% w/v, Sigma Aldrich, Munich, Germany) were performed for each cycle about 6 min at 37 °C. At the end of each cycle, the suspension was centrifuged for 5 min at 100 g and the supernatant was collected, pooled, and resuspended in cardiomyocytes culture medium (DMEM containing 10% v/v FBS, 100 M BrdU and penicillin-streptomycin solution). Isolated cells were pre-plated for 2 h on culture flasks in a humidified incubator at 37 °C with 5% $CO_2$ to separate cardiomyocytes and fibroblasts. Isolated cardiomyocytes were resuspended in cardiomyocytes culture medium. The density of cardiomyocytes was adjusted to $1.5 \times 10^5$ cells/mL. After neonatal rat cardiomyocytes had been cultured for 36-48 h, the medium was rinsed for further experiments.

**Transfection of cells**
For expression of fusion proteins, HeLa and U-2 OS cells were transfected 24-48 h prior to imaging using lipofectamine™ 3000 (L3000015; Thermo Fisher Scientific). Transfection was carried out according to the manufacturer’s protocols using 2.5 µg plasmid DNA. Cardiomyocytes were transfected on DIV1-2, and cells were switched from the culture medium into Opti-MEM™ (11058021; Thermo Fisher Scientific) for more than 30 min in the incubator. Thereafter, cells were transfected with 1.5 µg plasmids using 1.5 µL PLUS™ reagent (15338100; Thermo Fisher Scientific) and 1.5 µL Lipofectamine™ LTX reagent (15338100; Thermo Fisher Scientific) mixed in Opti-MEM™ Medium. After 12 h, the transfection medium was changed to the culture medium, and the transfected cardiomyocytes were incubated for 2-4 days before imaging experiments.

**Engineering of stable cell lines**

HeLa cells stably expressing pCMV-mEGFP-HaloTag7-PDGFR$^{tmb}$ were produced by PiggyBac (PB) Transposon System as described previously[12]. Stable cell lines were generated by co-transfecting the PB-pCMV-mEGFP-HaloTag7- PDGFR$^{tmb}$ and helper plasmid overexpressing Super PB Transposase (a gift from Prof. Wulan Deng) (System Biosciences, 1:1 molar ratio). Cells were transfected for 24 h using lipofectamine™ 3000 and then subjected to selection under puromycin (2-5 µg/mL) for 14 d. Cells were stained with **MaP555-HTL** (250 nM in DMEM) for 15 min at 37 °C with 5% $CO_2$ and washed twice with fresh DMEM for 30 min for subsequent fluorescence-activated cell sorting (FCAS). Cells with double positive fluorescent signals of GFP (Ex=488 nm, Em= 530 nm) and MaP555-HTL (Ex=561 nm, Em= 582 nm) were collected by BD FACS Aria™ III Cell Sorter (BD Biosciences, New Jersey, USA).

**rAAV production and neuron transduction**

Recombinant AAVs (rAAVs) were generated as described earlier.[8] pGP-AAV-hSyn1-CalR-HaloTag7-KDEL-WPRE-SV40 plasmid was transfected via Polyethylenimine 25000 (408727, Sigma-Aldrich) into HEK293 cells. The cells were harvested 5 days post transfection and lysed using TNT extraction buffer (20 mM Tris pH7.5, 150 mM NaCl, 1% TX-100, 10 mM $MgCl_2$). The cell debris was pelleted and the cell supernatant treated with Benzonase (Turbo Nuclease, EN-180S, Jena Bioscience). The rAAVs were purified from the medium and cell supernatant via FPLC using AVB Sepharose columns, which were subsequently concentrated using centrifugal filter devices (Amicon®, Merck KGaA, Darmstadt, Germany) with a MWCO of 100 kDa and buffer exchanged to PBS pH 7.3.

***In- vitro* wheat germ agglutinin (WGA) conjugation**

Compound **2**-succinimidyl ester (Compound **S23**) conjugate was synthesized from 6-carboxy tetramethyl rhodamine (CAS: 91809-67-5, E080753; energy chemical, Shanghai, China), followed by in-vitro labeling to WGA. **S23** was dissolved in dry DMSO (D12345; Thermo Fisher Scientific) to prepare a 1 mM stock solution. 90 µL 0.01M PBS, 100 µL 50 µM WGA (45064; Yuanye), and 10 µL **S23** stock solution was mixed in sequence. The reaction system was left shaking for 1 h at r.t. Afterwards, the reaction mixture was passed through Zeba Spin Desalting Columns (7K MWCO, 89882; Thermo Fisher Scientific) to remove unlabeled dyes to obtain the purified **GR555PM** for subsequent imaging experiments. Validation of the conjugation was performed by SDS-PAGE on 15% Tris-Gly gels for 10 min at 80 V and then 120 V for 1.5 h.

**In-cellulo fluorescence intensity, labeling kinetics and life-time measurement**

Confocal and fluorescence life-time imaging (FLIM) was performed at a Leica DMi8 microscope equipped with a Leica TCS SP8 X scanhead, SuperK white light laser, HC PL APO 40x/1.10 W motCORR CS2 water objective and hybrid detectors (Leica Microsystems) using the FALCON system with a pulsed laser at a frequency of 80 MHz.

U-2 OS Flp-IN T-REx$^{TM}$ cells stabely expressing H2B-HaloTag7-T2A-mEGFP[7] were grown in µ-Slide 8 Well Glass Bottom slides (80826; ibidi, Gräfelfing, Germany). 2 days prior to imaging 0.1 µg/mL doxycycline was added to the medium to induce the transgene expression. The cells were labeled using **MaP555-**, **GR555-**, **MaP618-** or **GR618-HTL** (500 nM) in phenol-red free medium (DMEM GlutaMAX™, 10% FCS, Gibco) for 2 h at 37 °C for end-point labeling intensity measurements. The fluorescence signal intensity was measured from (n≥80 cells) from three fields of view (FOVs) and normalized to the mEGFP signal for expression.

Live-cell labeling kinetic experiments were performed at 50 nM dye concentration and by recording z-stacks every 30s. The nuclear fluorescence intensity was quantified from >150 cells overtime from 3 individual experiments, averaged and normalized to the initial and maximal fluorescence signal.

For FLIM, experiments, the fluorescent nuclei were imaged, collecting 1,000 photons per pixel from which the fluorescence lifetimes (τ) were derived (n≥3 images, mean value and standard deviation). Background signal from empty coverslip space was removed (threshold). Mean fluorescence lifetimes were calculated in the LAS X software (Leica Microsystems) by fitting a mono-exponential decay model to the decay ($\chi^2 < 1.2$).

**Viability-based phototoxicity assay of mitochondrial dyes**

HeLa cells were grown to 80-90 % confluency in a 96-well plate (costar 3599; Corning) before treatment with 250 nM TMRM and compounds **1-4** (for fig. 1f), or 250 nM compound **6** (SiRM) or **7** (GR650M) (for fig.2e), respectively in DMEM for 15 min at 37°C with 5% $CO_2$. Then, the cells were washed with PBS twice and maintained in fresh medium for subsequent imaging analysis. 96-well plates were analyzed using a high-content imaging system Opera Phenix (PerkinElmer, Waltham, MA, USA) equipped with a 20 ×/0.4 NA air objective and live-cell imaging device. The average light intensity on cells was measured to be 1.4 W/cm$^2$ (561 nm) or 1.2 W/cm$^2$ (640 nm). The cells in each well were irradiated with the maximum light intensity at different time points. Time points were selected as 0, 0.5, 1, 2, 5, and 10 min (for fig. 1h); 0, 1, 2, 5, 10, 15, and 20 min (for fig. 2e). Each time point with three parallel repeats. After illumination, the cells were directly used for subsequent imaging analysis. The cells were washed with PBS once and a 100 µL cell viability assay solution containing 5 µM propidium iodide (PI, M155695; MREDA, Beijing, China) was added to each sample. Cells without dye incubation were included as controls. Images of two channels (PI: Cy3 channel, 568 nm; Digital phase contrast (DPC) mode) were recorded on Opera Phenix. The cell viability (%) was calculated according to the following equation: cell viability % = (A-B) / A *100%, where A = the number of total cells before illumination using DPC mode, and B = the number of PI-positive cells after illumination. More than 400 cells were counted at each time point.

**Viability-based phototoxicity assay of WGA-TMRM or GR555PM**

HeLa cells were grown to 80-90 % confluency in a 96-well plate (costar 3599; Corning) before 30 µg/mL **WGA-TMRM** or 50 µg/mL **GR555PM** in HBSS (Gibco 14025092; Thermo Fisher Scientific) were added, respectively for 5 min at 37 °C with 5% $CO_2$. Then, the cells were washed with HBSS twice and maintained in fresh medium for subsequent imaging analysis. 96-well plates were analyzed using a high-content imaging system ImageXpress Micro XLS (Molecular Devices, San Jose, CA, USA) equipped with a 20 ×/0.4 NA air objective and live-cell imaging device. The average light intensity on cells was measured to be 2.6 W/cm$^2$ (532 nm). The cells in each well were irradiated with the maximum light intensity at different time points. Time points were selected as 0, 2, 4, 6, 8, 10, and 12 min. Each time point with three parallel repeats. After illumination, the cells were directly used for subsequent imaging analysis. The cells were washed with PBS once and a 100 µL cell viability assay solution containing 1 µM Calcein AM (C2012-0.1ml; Beyotime, Shanghai, China) was added to each sample. Cells without dye incubation were included as controls. Images of two channels (plasma membrane: Cy3 channel, 568 nm; Calcein AM: FITC channel, 488 nm) were recorded on ImageXpress Micro XLS. The cell viability (%) was calculated according to the following equation: cell viability % = B / A *100%, where A = the number of total cells before illumination, and B = the number of Calcein AM-positive cells after illumination. More than 500 cells were counted at each time point.

**Measurement of phototoxicity of Rho123 and GR510M using analysis of mitochondrial membrane potential (MMP)**

HeLa cells were grown to 80-90 % confluency in a 96-well plate (costar 3599; Corning) before treatment with 250 nM **Rho123** or **GR510M** for 15 min at 37 °C with 5% $CO_2$. Then, the cells were washed with PBS twice and maintained in fresh medium for subsequent imaging analysis. Glass slides coated with polymer films containing 1 µM **Rho123** or **GR510M** were used as an in-vitro control for bleaching correction and were placed on the same 96-well plate. 96-well plates were analyzed using a high-content imaging system Opera Phenix (PerkinElmer, Waltham, MA, USA) equipped with a 20 ×/0.4 NA air objective and live-cell imaging device. The average light intensity on cells was measured to be 1.7 W/cm$^2$ (488 nm). The samples in each well were irradiated with the maximum light intensity at different time points. Time points were selected as 0.5, 1, 2, and 5 min. Each time point with three parallel repeats. The drop in MMP for each sample was measured by the decrease in a fluorescent signal of **Rho123** or **GR510M**. The mean grey values for each period were evaluated in Fiji after bleach-correcting the experimental groups using control samples. More than 400 cells were analyzed at each time point of experimental samples.

**Measurement of cytotoxicity using Cell Counting Kit-8 (CCK8) reagent**

HeLa cells were grown to 80-90 % confluency in a 96-well plate (costar 3599, Corning) the day before treatment. Cells were either stained with **TMRM**, **GR555M**, **MaP555-DNA**, and **GR555-DNA** with different concentrations (0.25, 0.5, 1, 2, 5 µM in DMEM) 2 hours at 37 °C with 5% $CO_2$. Cells were washed with PBS once after the staining procedure and incubated with cultured medium containing 10% Cell Counting Kit-8 reagent (CCK8; C0038; Beyotime) for 2 hours at 37 °C with 5% CO2. The absorbance (450 nm) of each well was recorded using a TECAN Infinite M Nano+ microplate reader (TECAN, Männedorf, Switzerland).

**Measurement of DNA damage using hXRCC1-GFP assay**
HeLa cells or U-2 OS EF1α H2B-HaloTag7 cells transiently expressing hXRCC1-GFP were seeded in glass-bottom dishes (STGBD-035-1; Standard Imaging, Beijing, China) one day prior to imaging at 37 °C with 5% $CO_2$. Cells were labeled with **MaP555-DNA** (200 nM in DMEM, 60 min), **GR555-DNA** (2000 nM in DMEM, 60 min), or **MaP555-HTL** or **GR555-HTL** (500 nM in DMEM, 30 min) respectively. After removing the staining solution, the cells were washed with DMEM once and maintained in fresh DMEM for following confocal imaging at r.t. The labeled live cells were recorded using a Facility Line STED microscope (Abberior Instruments GmbH, Göttingen, Germany) equipped with an Olympus UPlanXAPO 60x oil, NA1.42 objective (Olympus, Tokyo, Japan). The dual-color time-lapse imaging of hXRCC1-GFP and DNA or H2B-HaloTag7 was recorded with a time interval of 2 minutes in confocal mode. During the time interval, the cells in the field of view were continuously irradiated by a 561-nm pulsed laser with maximum power. The live-cell dual-color confocal recordings were recorded with a fixed frame size of 80 × 80 µm and a pixel size of 100 nm. hXRCC1-GFP was excited at $\lambda_{ex}$ = 485 nm, and the dyes for DNA/H2B-HaloTag7 labeling were excited at $\lambda_{ex}$ = 561 nm. The pixel dwell time was set to 6.5 µs and 2-line accumulations were recorded for each channel.

**Live-cell long-term confocal microscopy**
Wild-type HeLa and HeLa IgKO-HaloTag7-mEGFP-pDisplay cells were seeded in glass-bottom dishes (STGBD-035-1; Standard Imaging) one day prior to imaging at 37 °C with 5% $CO_2$. Then, wild-type HeLa cells were stained with **MaP555-Actin** or **GR555-Actin** (100 nM in DMEM containing 10 µM verapamil, 3 h) and HeLa pDisplay cells were stained with **MaP555-/GR555-HTL** (500 nM in DMEM, 30 min) at 37 °C with 5% $CO_2$. After removing the staining solution, the cells were then washed with DMEM once and maintained in fresh DMEM (for actin labeling containing 10 µM verapamil) for following confocal imaging at r.t. Cells were recorded using a STEDYCON STED microscope equipped with a CFI Plan Apochromat Lambda D 100x oil, NA1.45 objective (Nikon, Tokyo, Japan), and a pulsed excitation laser at 561 nm wavelength. Long-term time-lapse imaging was recorded in the confocal mode without additional time interval, with a frame size of ~50 × 50 µm and a pixel size of 100 nm. The pixel dwell time was set to 5 µs and 2-line accumulations were recorded.

Hippocampal rat neurons were transduced with pGP-AAV-hSyn1-CalR-HaloTag7-KDEL-WPRE-SV40 rAVVs after 5 days in culture (DIV). After 10 DIV, the neurons were stained with **MaP555-/GR555-DNA** (500 nM /2 µM), **MaP618-/GR618-HTL** (500 nM) and **SiRM**/**GR650M** (50 nM) at 37 °C with 5% $CO_2$ for 1h. Long-term time-lapse confocal imaging was recorded on a Stellaris 5 inverted microscope (Leica) equipped with a white line laser (WLL) and hybrid photodetectors. A 40x/1.10 water immersion objective was used at to image a 12 z-planes at 1024x1024 pixel resolution (0.75x zoom, 600 Hz scan speed, with a frame size of 387.5 x 387.5 µm). The pixel dwell time was 862 ns and the three following channels were selected: Ext.: 540 nm (8%), Em.: 550 – 580 nm. Ext.: 600 nm (20%), Em.: 610 – 630 nm. Ext.: 660 nm (6%), Em.: 670 – 770 nm.

**STED microscopy**
For single-color STED microscopy, COS-7 cells were stained with **GR650M** (250 nM in DMEM) at 37 °C with 5% $CO_2$ for 1h. After removing the staining solution, the cells were then washed with DMEM once and maintained in fresh DMEM. Live cells STED nanoscopy was performed Facility Line STED microscopes (Abberior Instruments GmbH, Göttingen, Germany) equipped with an Olympus UPlanXAPO 60x oil, NA1.42 objective (Olympus, Tokyo, Japan). Pixel sizes of 30 nm were used for STED nanoscopy. **GR650M** was excited at 640 nm wavelength and STED was performed using a pulsed depletion laser at 775 nm wavelength with gating of 1-7 ns and dwell times of 10 µs.

For dual-color STED microscopy, hippocampal rat neurons expressing CalR-HaloTag7-KDEL were stained with **MaP618-/GR618-HTL** (500 nM) and compound **5/GR650M** (50 nM) at 37 °C with 5% $CO_2$ for 1h. Live cells STED nanoscopy was performed using an Abberior STED Expert Line 595/775/RESOLFT QUAD scanning microscope (excitation lines: 561 nm, 640 nm; STED lines: 775 nm) equipped with a UPlanSApo 100x/1.4 oil immersion objective lens (Abberior Instruments). Detection was performed with avalanche photodiodes (APD) and spectral detection. Images were recorded at 10 x 10 and 6 x 18 µm frame size, 60 nm (confocal) or 30 nm (STED) pixel sizes, 15 µm pixel dwell time and 4 times line accumulation. The two following channels were selected: Ext.: 561 nm (30%), Em.: 587 – 630 nm. Ext.: 640 nm (0.2%), Em.: 670 – 746 nm. STED 775 nm (20%). 'Hot' LUT were applied for data representation.

For time-lapsed STED microscopy, U-2 OS FlpIn TRex™ TOM20-HT7 cells[13] were stained with **MaP618-/GR618-HTL, -S5** or **-Hy5** (500 nM) at 37 °C with 5% $CO_2$ for 1h. Live cells STED nanoscopy was performed at the same set-up as described above. Images were recorded at 8 x 8 µm frame size, 30 nm (STED) pixel sizes, 15 µm pixel dwell time and 2 times line accumulation over 50 consecutive frames Ext.: 640 nm (2%), Em.: 670 – 746 nm. STED 775 nm (20%). 'Hot' LUT were applied for data representation. The mean pixel values (borders were excluded) over the image series from at least 4 image series ($n \geq 4$) was extracted using *ImageJ* image. To generate bleaching curves, the data was background corrected and normalized to the first frame intensity. Average bleaching curve and single traces are shown.

**Voltage imaging**

Neonatal rat cardiomyocytes expressing Voltron (Ace2-HaloTag7) were labeled with **MaP555-HTL** or **GR555-HTL** (100 nM in DMEM) at 37 °C with 5% $CO_2$ for 25 min. After removing the staining solution, the cells were then washed with HBSS once and maintained in fresh culture medium. Voltron imaging were performed with an inverted fluorescence microscope (Nikon-TiE) equipped with a objective CFI Plan Fluor 40× oil, NA 1.3 (Nikon, Tokyo, Japan), one laser line (561 nm, Coherent OBIS), and one scientific CMOS camera (Hamamatsu ORCA-Flash 4.0 v2). The microscope, laser, and camera were controlled by LabVIEW software (National Instruments, 15.0 version). Images were captured using a camera on 2×2 pixel binning and a 100-ms exposure duration. Images were analyzed using ImageJ/Fiji (version 1.52d).

### Supplementary Figures

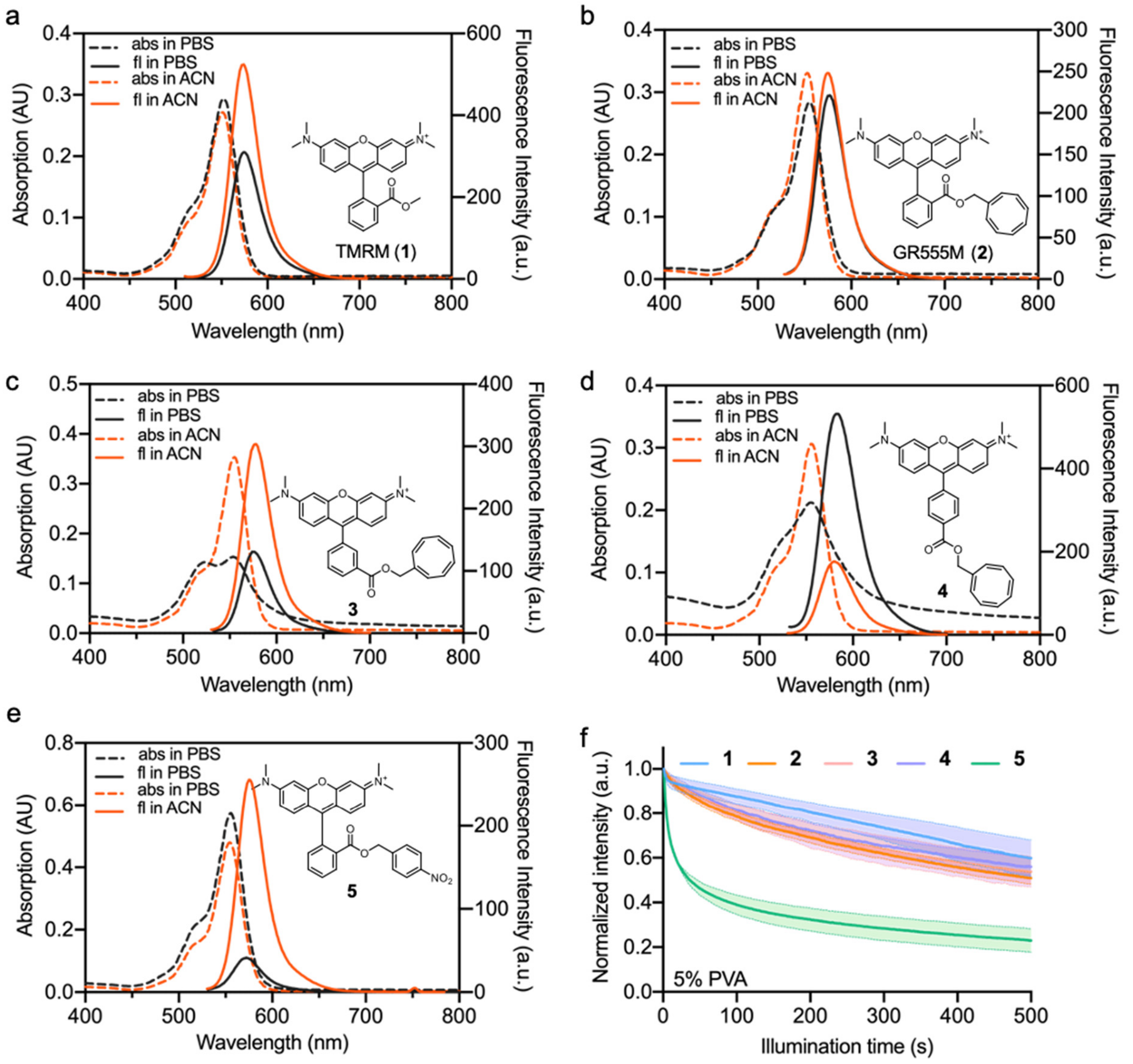


**Figure S1. Photophysical properties of TMRM and compounds 1-5.** (a-d) Absorption (abs) and fluorescence (fl) spectra of and compounds **1** (**TMRM**) (a), **2** (**GR555M**) (b), **3** (c), **4** (d), and **5** (e) in acetonitrile and phosphate buffer (PBS). The chemical structures of the corresponding compounds marked in the figures. (f) Photobleaching curves of compounds **1**-**5** in polyvinyl alcohol (PVA) film recorded under continuous 561 nm laser scanning of a confocal microscope. Data points represent averaged fluorescence bleaching curves of three independent replicates. Error bars, showing light-shaded areas, indicate standard deviation.

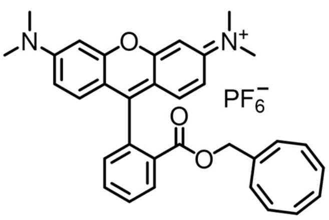


Compound **2** (GR555M)

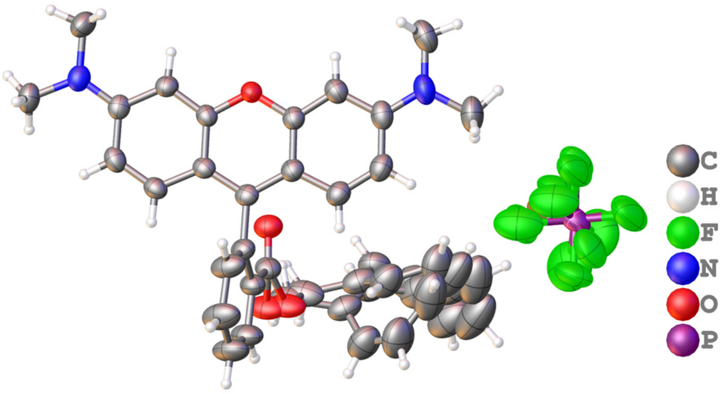


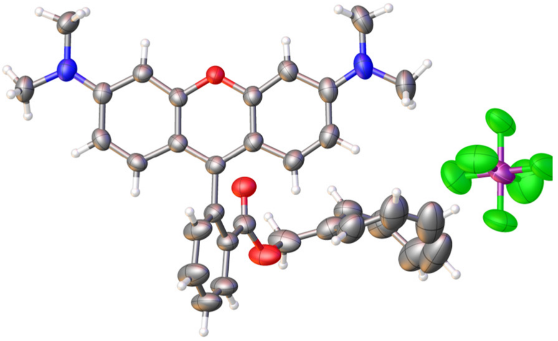

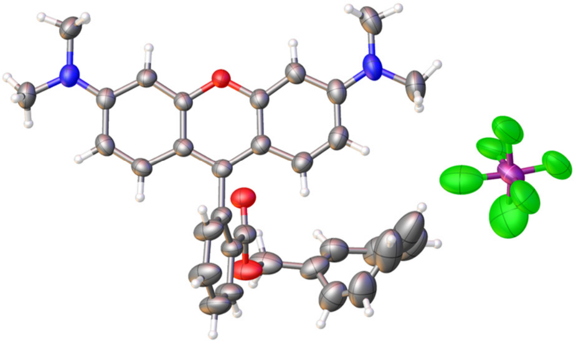

**Major component**

Cyclooctatetraene part: 78% occupancy;
$PF_{6^-}$ part: 90% occupancy.

**Minor component**

Cyclooctatetraene part: 22% occupancy;
$PF_{6^-}$ part: 10% occupancy.

**Figure S2. Single crystal structures of compound 2 (GR555M)**. Single crystals of compound **2** (**GR555M**) were obtained via evaporation at 24 °C from its acetonitrile solution. A suitable crystal was selected and measured on a XtaLAB Synergy R, DW system, HyPix diffractometer. The crystal was kept at 180.00 (10) K during data collection. Using Olex2[14], the structure was solved with the SHELXT[15] structure solution program using Intrinsic Phasing and refined with the SHELXL[15] refinement package using Least Squares minimization.

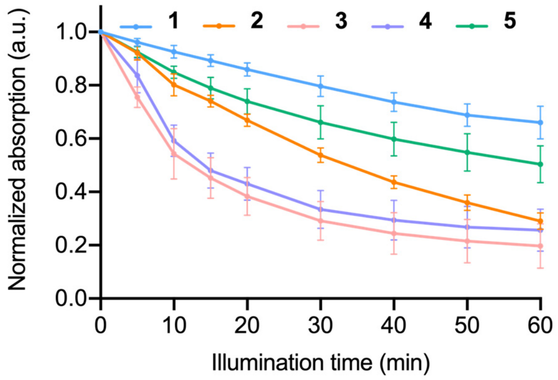


**Figure S3. Photobleaching of compounds 1-5 in phosphate buffer (PBS)**. Photobleaching curves of compound **1-5** (10 μM) in PBS by illuminated with green LED (520-530 nm, 50 mW/cm$^2$). Data points represent averaged fluorescence bleaching curves of three independent replicates. Error bars indicate standard deviation.

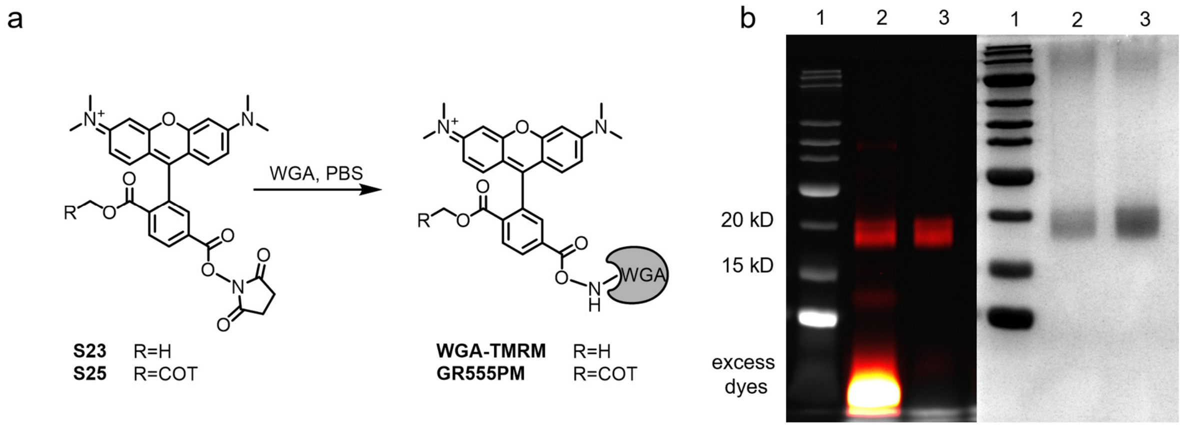


**Figure S4. Preparation of the live-cell plasma membrane probes.** (a) Conjugation reaction of **GR555PM** and **WGA-TMRM**. (b) In-gel fluorescence (left) and Coomassie brilliant blue staining (right) analysis of reaction mix (lane 2), and desalted conjugate (lane 3).

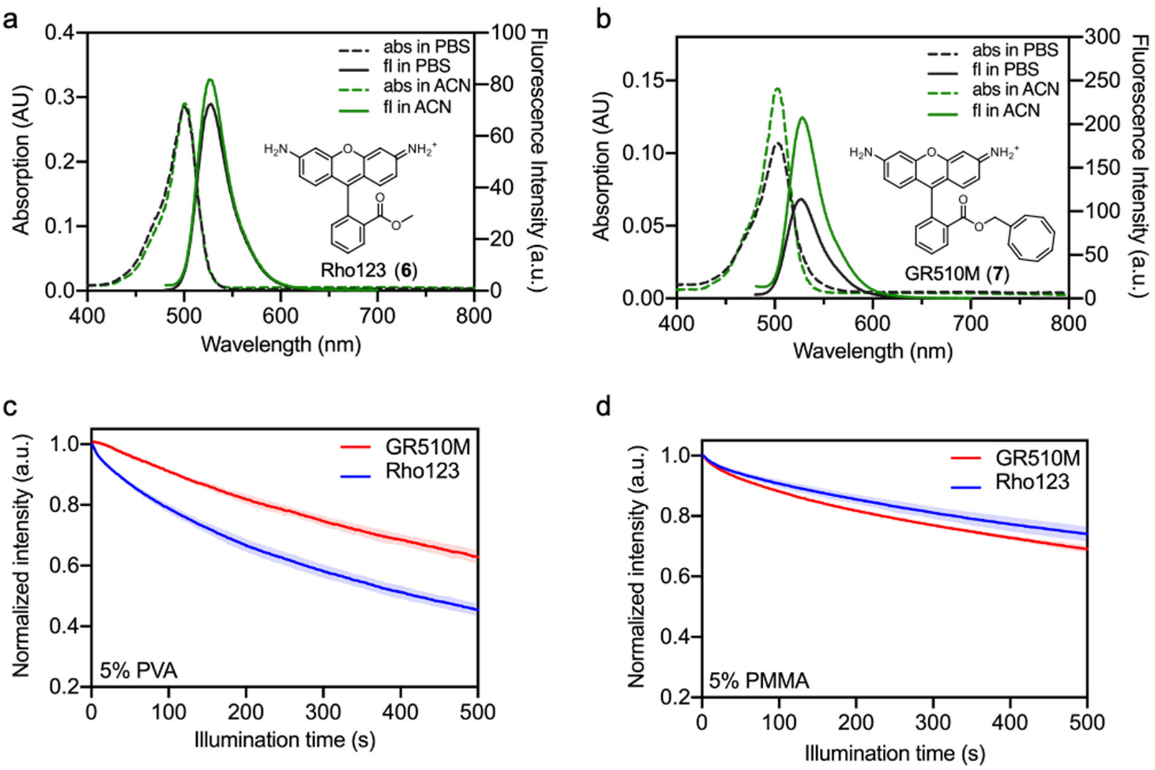


**Figure S5. Photophysical properties of compound 6 (Rho123) and compounds 7 (GR510M).** (a-b) Absorption (abs) and fluorescence (fl) spectra of **Rho123** (a) and **GR510M** (b) in acetonitrile and PBS. The chemical structures of the corresponding compounds are marked in the figures. (c-d) Photobleaching curves of **Rho123** and **GR510M** in polyvinyl alcohol (PVA) (c) and polymethyl methacrylate (PMMA) film (d) recorded under continuous 488 nm laser scanning of a confocal microscope. Data points represent averaged fluorescence bleaching curves of three independent replicates. Error bars, showing light-shaded areas, indicate standard deviation.

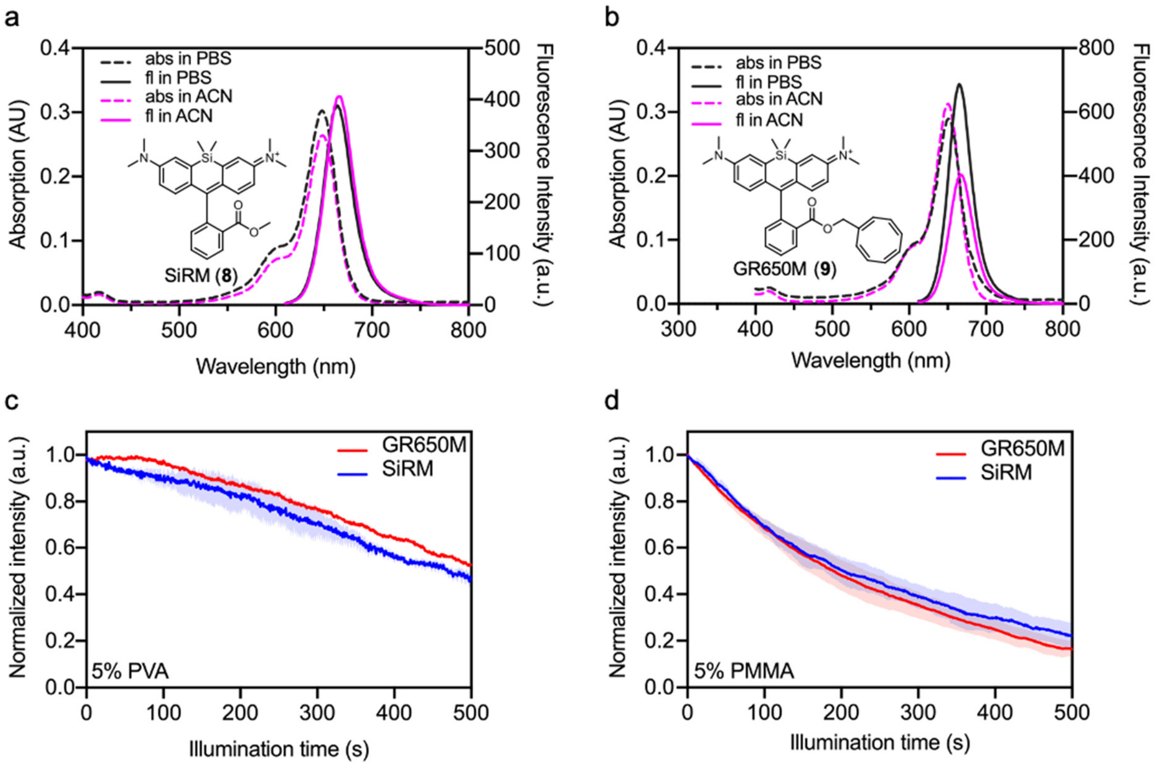


**Figure S6. Photophysical properties of compounds 8 (SiRM) and 9 (GR650M).** (a-b) Absorption (abs) and fluorescence (fl) spectra of **SiRM** (a) and **GR650M** (b) in acetonitrile and PBS. The chemical structures of the corresponding compounds are marked in the figures. (c-d) Photobleaching curve of compound **6** (**SiRM**) and **7** (**GR650M**) in polyvinyl alcohol (PVA) (c) and polymethyl methacrylate (PMMA) film (d) recorded under continuous 650 nm laser scanning of a confocal microscope. Data points represent averaged fluorescence bleaching curves of three independent replicates. Error bars, showing light-shaded areas, indicate standard deviation.

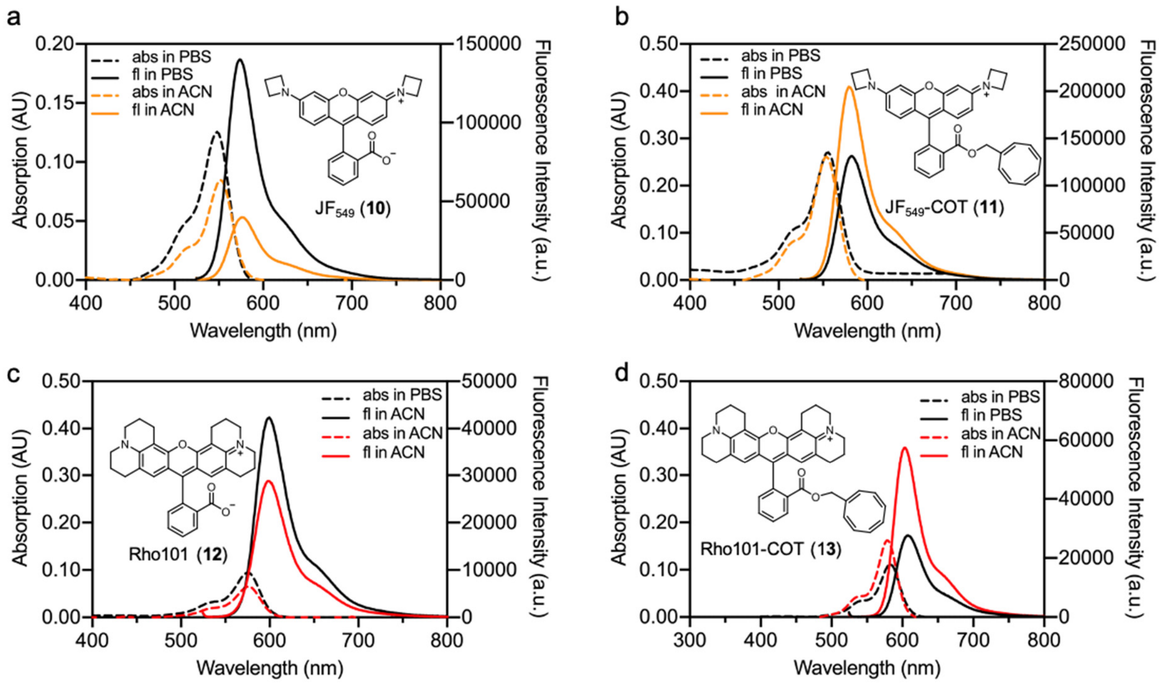


**Figure S7. Photophysical properties of compounds 10-13**. (a-d) Absorption (abs) and fluorescence (fl) spectra of and compounds **10:** $JF_{549}$ (a),**11**: $JF_{549}$-COT (b), **12**: Rho101 (c), and **13**: Rho101-COT (d) in acetonitrile and phosphate buffer (PBS). The chemical structures of the corresponding compounds marked in the figures.

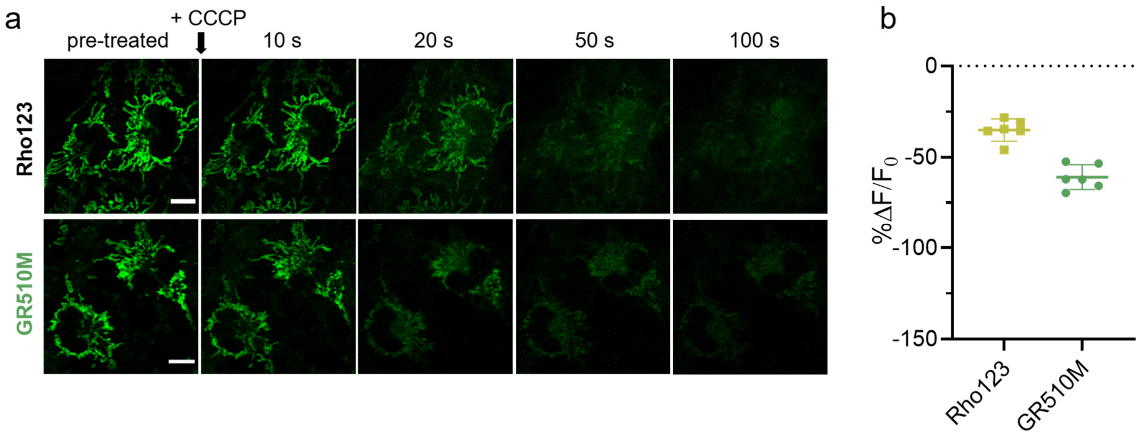


**Figure S8. Response of Rho123 and GR510M to oxidative phosphorylation uncoupler carbonyl cyanide 3-chlorophenylhydrazone (CCCP).** (a) live-cell time-lapse recording of COS-7 cells labeled with **Rho123** and **GR510M** (300 nM in DMEM, 60 min) before and after the addition of CCCP. Scale bars = 10 μm. (b) Fraction change in fluorescence (ΔF/F) at 100 s after the addition of CCCP. Data points represent average $\Delta F/F_0$ of six cells from two independent biological replicates. Error bars indicate the standard deviation.

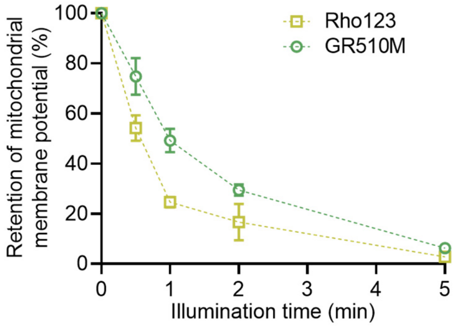


**Figure S9. Phototoxicity of GR510M or Rho123 on live HeLa cells.** Phototoxicity of **GR510M** or **Rho123** (200 nM, 30 min) in HeLa cells, as measured by the analysis of fluorescence retention in mitochondria after LED illumination at different times. Data points indicate the mean of at least 1,500 individual cells from three independent biological replicates. Error bars indicate standard deviation.

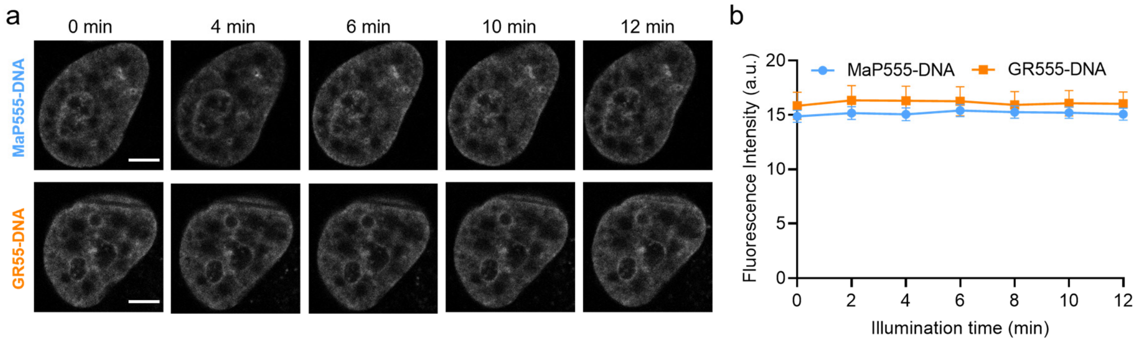


**Figure S10. Photostability of MaP555-DNA and GR555-DNA on live HeLa cells.** (a) Live-cell time-lapse recordings of HeLa hXRCC1-GFP transiently expressing cells labeled with **MaP555-DNA** (200 nM in DMEM, 60 min) or **GR555-DNA** (2 μM in DMEM, 60 min) respectively under continuous 561 nm laser scanning. The snapshots were acquired with 2 min intervals. Scale bars = 2 μm. (b) Photobleaching curves of **MaP555-DNA** and **GR555-DNA** on HeLa hXRCC1-GFP transiently expressing cells in time-lapse recordings. Data points represent average bleaching curves of eleven cells from four independent biological replicates. Error bars indicate the standard error of the mean (SEM). Frame number = 7; Frame rate = 2 min/frame; Duration time = 12 min.

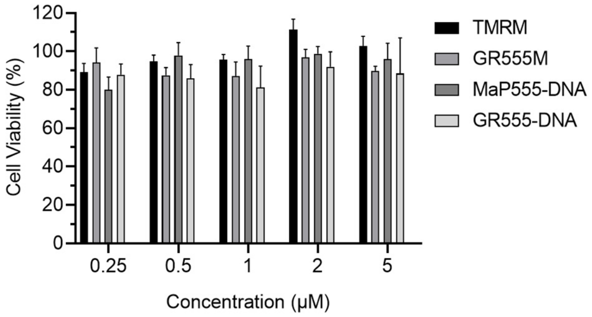


**Figure S11. Cytotoxicity assay of HeLa cells stained with mitochondrial probes (1, 2) and DNA dyes (14, 15).** Cytotoxicity of HeLa cells treated with **TMRM**, **GR555M**, **MaP555-DNA**, **GR555-DNA** in different staining concentrations (0.25, 0.5, 1, 2, 5 μM) for 2 hours. Bars indicate the mean of three individual repeats. Error bars indicate the standard error of the mean (SEM).

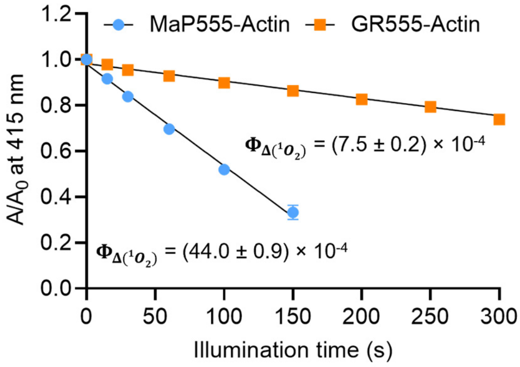


**Figure S12. In-vitro singlet oxygen generation of MaP555-Actin and GR555-Actin.** Singlet oxygen yield of **MaP555-Actin** or **GR555-Actin** in acetonitrile containing 0.1% trifluoroacetic acid based on the DPBF assay under continuous illumination with green LED (520-530 nm, 50 mW/cm$^2$). The absolute singlet oxygen yield is marked on the graph; Units: × $10^{-4}$. Data points represent averaged DPBF bleaching curves and standard deviation of three independent replicates.

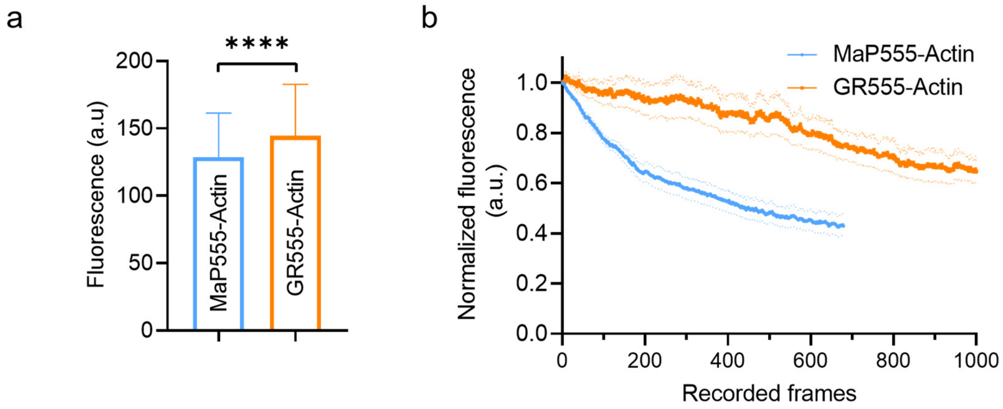


**Figure S13. Brightness and photobleaching of MaP555-Actin and GR555-Actin on live HeLa cells.** (a) Comparison of actin brightness on HeLa cells labeled with **MaP555-Actin** or **GR555-Actin** (100 nM in DMEM containing 10 μM verapamil, 3h). Bars indicate the mean of at least six individual cells from two independent biological repeats. Error bars indicate standard deviation. Significance was determined using two- tailed unpaired t-test. $P$ = **** < 1.0 $\times 10^{-4}$. (b) Photobleaching curves of HeLa cells labeled with **MaP555-Actin** or **GR555-Actin** (100 nM in DMEM containing 10 μM verapamil, 3h) during time-lapse confocal recordings. Curves show averaged bleaching curves of at least six individual cells from two independent biological replicates. Error bars show the standard error of the mean (SEM). Frame number = 679 or 1000 (for **MaP555-Actin** or **GR555-Actin**); Frame rate = 7.09 sec/frame; Duration time = 82 /121min (for **MaP555-Actin** or **GR555-Actin**).

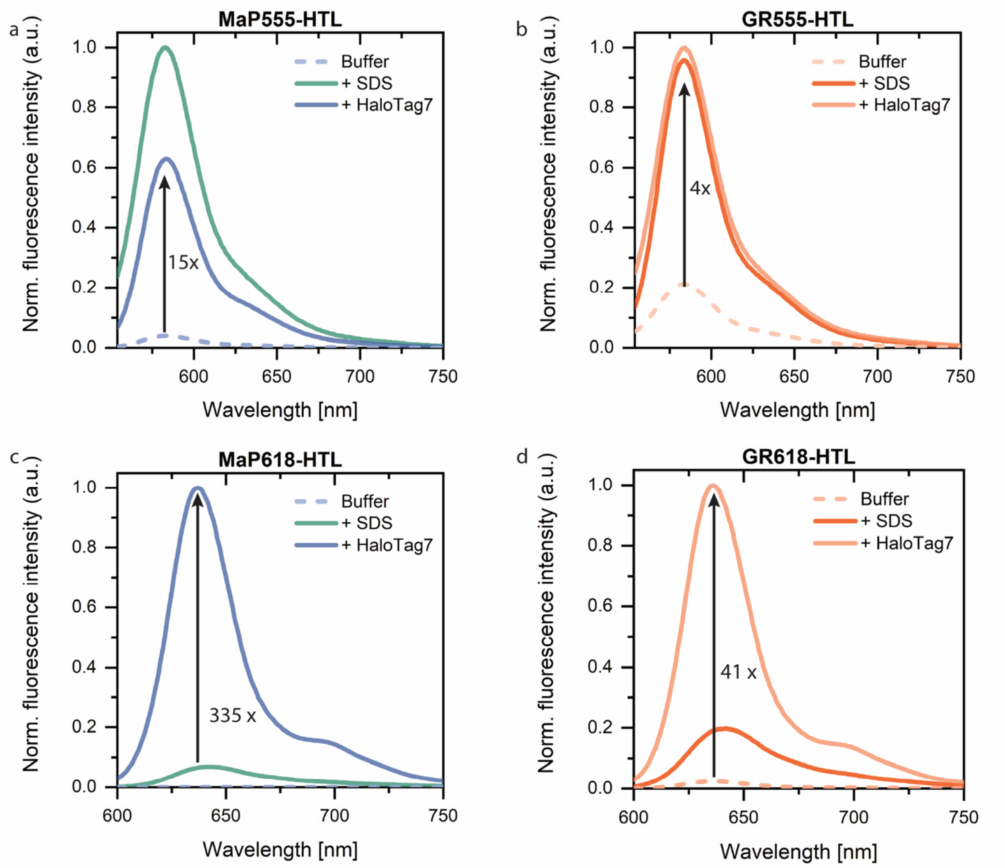


**Figure S14. Fluorescence emission response of regular (a, c) and gentle rhodamine HaloTag Ligands (b, d) to covalent HaloTag7 labeling.** Normalized fluorescence emission spectra of 50 nM probe measured in the absence (dashed line) and presence of HaloTag7 (15 µM) or SDS (0.1%, both straight lines) after 2 h incubation. Data from three technical replicates was averaged and the mean ratio of fluorescence intensities at the emission maxima in the presence and absence of HaloTag7 was calculated as indicated.

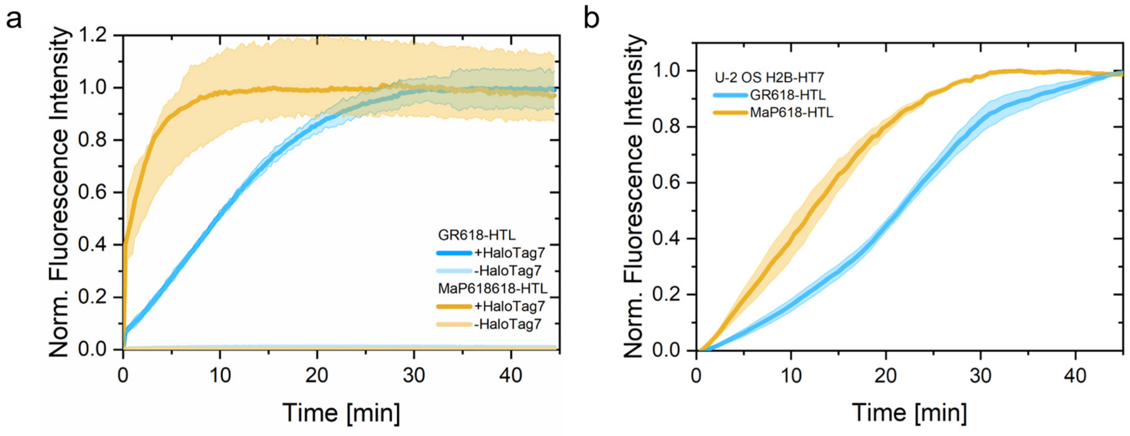


**Figure S15. Labeling kinetics of regular and gentle rhodamine HaloTag Ligands.** (a) In-vitro labeling kinetics of **GR618-/MaP618-HTL** (500 nM) with 50 µM HaloTag7 at 37 °C. The fluorescence increase was measured every 15 s in technical triplicates, averaged and normalized to the initial fluorescence signal, standard deviation given. The fluorescence intensity in absence of HaloTag7 was tracked as well. Full labeling (95% $I_{max}$) was reached after 7.2 min (**MaP618-HTL**) or 26.2 min (**GR618-HTL**). (b) Live-cell labeling kinetics of **GR618-/MaP618-HTL**. U-2 OS H2B-HT7 cells were incubated with 500 nM of **GR618-/MaP618-HTL** at 37 °C. The nuclear fluorescence intensity was quantified from >150 cells from 3 individual experiments, averaged and normalized to the initial and maximal fluorescence signal, error presented by the S.E.M.

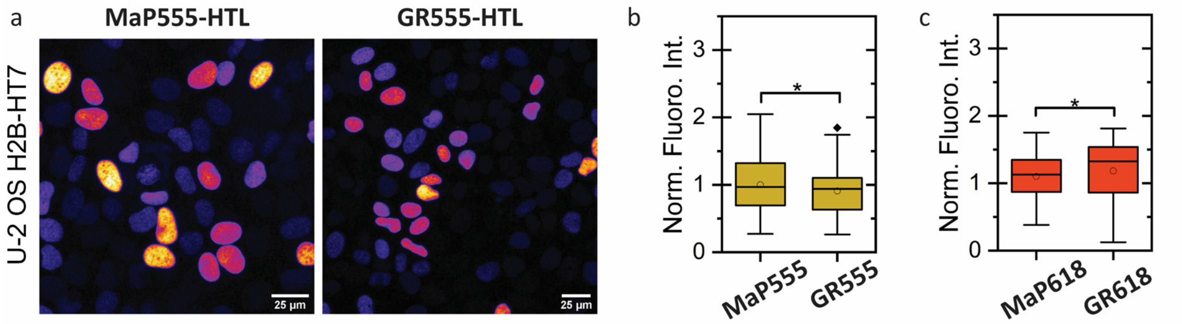


**Figure S16. Cellular brightness comparison of regular and gentle rhodamine HaloTag Ligands.** (a) Live-cell staining comparison of **MaP555-HTL** and **GR555-HTL** (500 nM). Confocal microscopy images taken of U-2 OS H2B-HaloTag7-T2A-mEGFP cells under no-wash conditions. Scales: 25 µm. (b, c) Quantification of the nuclear fluorescence intensity normalized to the protein expression of ≥50 cells. Box = 25%–75% percentile and whiskers = 5%–95% percentile, circle = mean, line = median, rhombus = outlier. Significance was calculated using two-sided t-tests including the Welch correction: *P* = n.s. (0.10, 0.14); * $1.5\times10^{-2}$

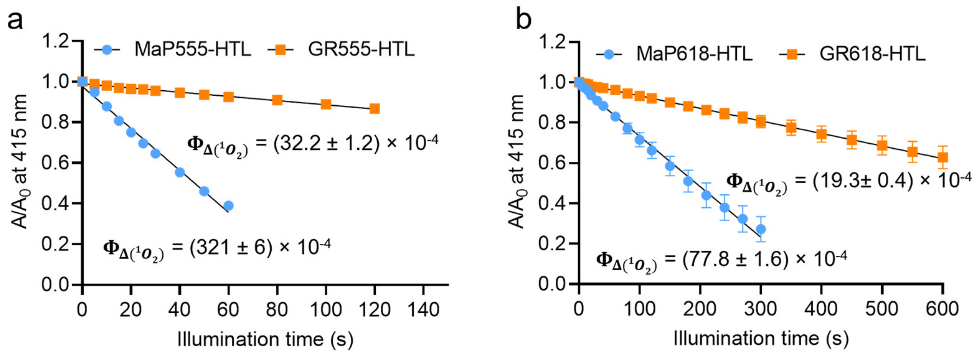


**Figure S17.** ***In-vitro* singlet oxygen generation of MaP555-/MaP618-HTL and GR555-/GR618-HTL.** (a) singlet oxygen yield of **MaP555-HTL** and **GR555-HTL** in acetonitrile containing 0.1% trifluoroacetic acid based on the DPBF assay under continuous illumination with green LED (520-530 nm, 50 mW/cm$^2$). The absolute singlet oxygen yield is marked on the graph; Units: × $10^{-4}$. Data points represent averaged DPBF bleaching curves and standard deviation of three independent repeats. (b) singlet oxygen yield of **MaP618-HTL** and **GR618-HTL** in acetonitrile containing 0.1% trifluoroacetic acid based on the DPBF assay under continuous illumination with green LED (620-630 nm, 50 mW/cm$^2$). The absolute singlet oxygen yield is marked on the graph; Units: × $10^{-4}$. Data points represent averaged DPBF bleaching curves and standard deviation of three independent replicates.

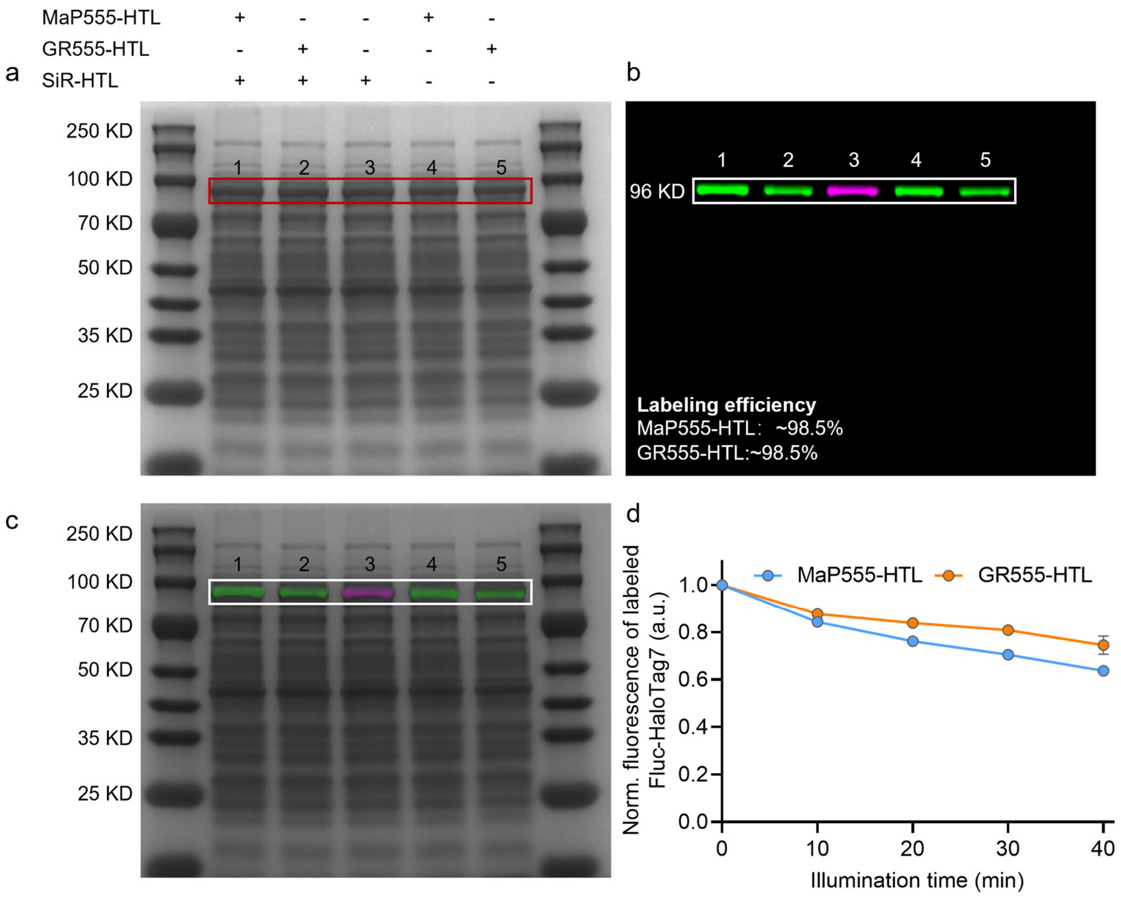


**Figure S18. Analysis of the labeling efficiency and photobleaching of firefly luciferase-HaloTag7 protein (Fluc-HaloTag7).** Fluc-HaloTag7 was first labeled with **MaP555-HTL** (lane 1,4), **GR555-HTL** (lane 2,5), or **SiR-HTL** (lane 3) respectively *in-vitro*, and then labeled with **SiR-HTL** (lane 1,2) again, as a pulse-chase experiment, for the measurement of labeling efficiency of **MaP555-HTL** or **GR555-HTL**. (a) In-gel Coomassie brilliant blue staining analysis of E. coli lysate, recombinant expression of Fluc-HaloTag7 protein. (b) In-gel fluorescence analysis of E. coli lysate. Recombinantly expressed Fluc-HaloTag7 was labeled with **MaP555-HTL** or **GR555-HTL** (green, Cy3 channel) and **SiR-HTL** (magenta, Cy5 channel). (c) Merged SDS-PAGE electrophoresis of (a) and (b). The absence of a fluorescent band in the Cy5 channel of lane 1,2 suggests that Fluc-HaloTag7 is already fully labeled with **MaP555-HTL** and **GR555-HTL**. (d) Photobleaching of Fluc-HaloTag7 labeled with **MaP555-HTL** or **GR555-HTL** respectively under continuous illumination with green LED (520-530 nm, 50 mW/cm$^2$). Data points represent averaged fluorescence bleaching curves of three independent replicates. Error bars indicate the standard error of the mean (SEM).

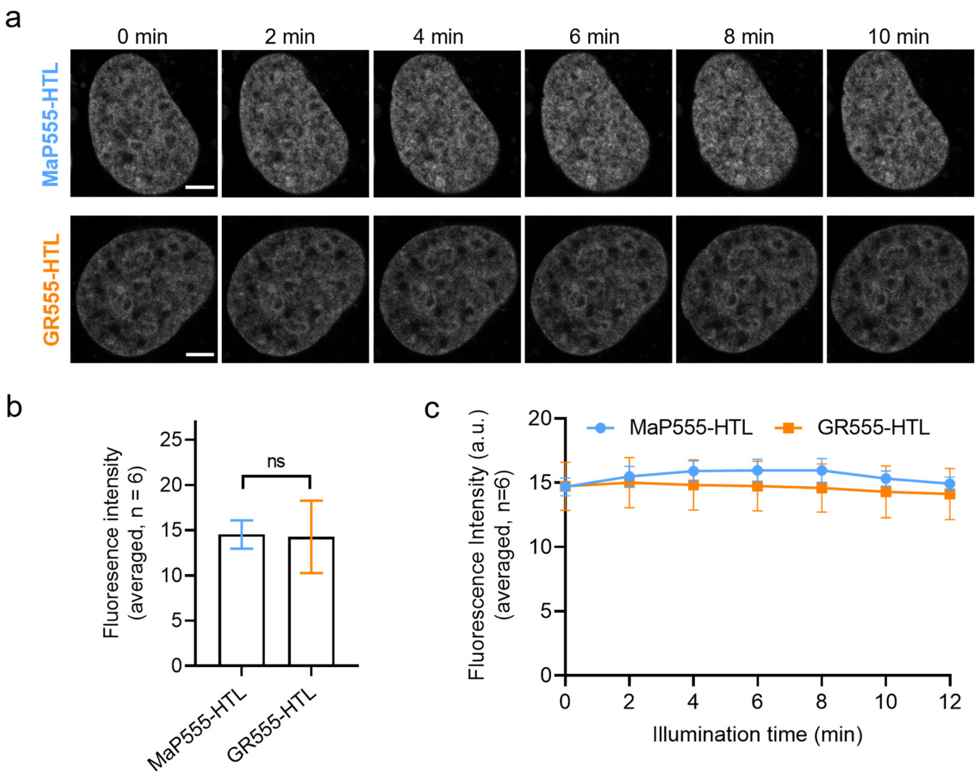


**Figure S19. Staining of MaP555-HTL and GR555-HTL on U-2 OS H2B-HaloTag7 expressing cells.** (a) Time-lapse confocal microscopy of U-2 OS H2B-HaloTag7 expressing cells labeled with **MaP555-HTL** or **GR555-HTL** (500 nM in DMEM, 30 min) with a time interval of 2 min. During the time interval, the cells in the field of view were continuously irradiated by a 561-nm pulsed laser. (b) Comparison of the cellular brightness of **MaP555-HTL** and **GR555-HTL** (500 nM in DMEM, 30 min) on U-2 OS H2B-HaloTag7 expressing cells. Bars indicate the mean of six samples. Error bars indicate standard deviation. Significance was determined using two- tailed unpaired t-test. *P* = n.s. (0.89). (c) Photobleaching curves of **MaP555-HTL** and **GR555-HTL** on U-2 OS H2B-HaloTag7 expressing cells in time-lapse recordings. Data points represent average bleaching curves of six cells from four independent biological replicates. Error bars indicate the standard error of the mean (SEM). Frame number = 7; Frame rate = 2 min/frame; Duration time = 12 min.

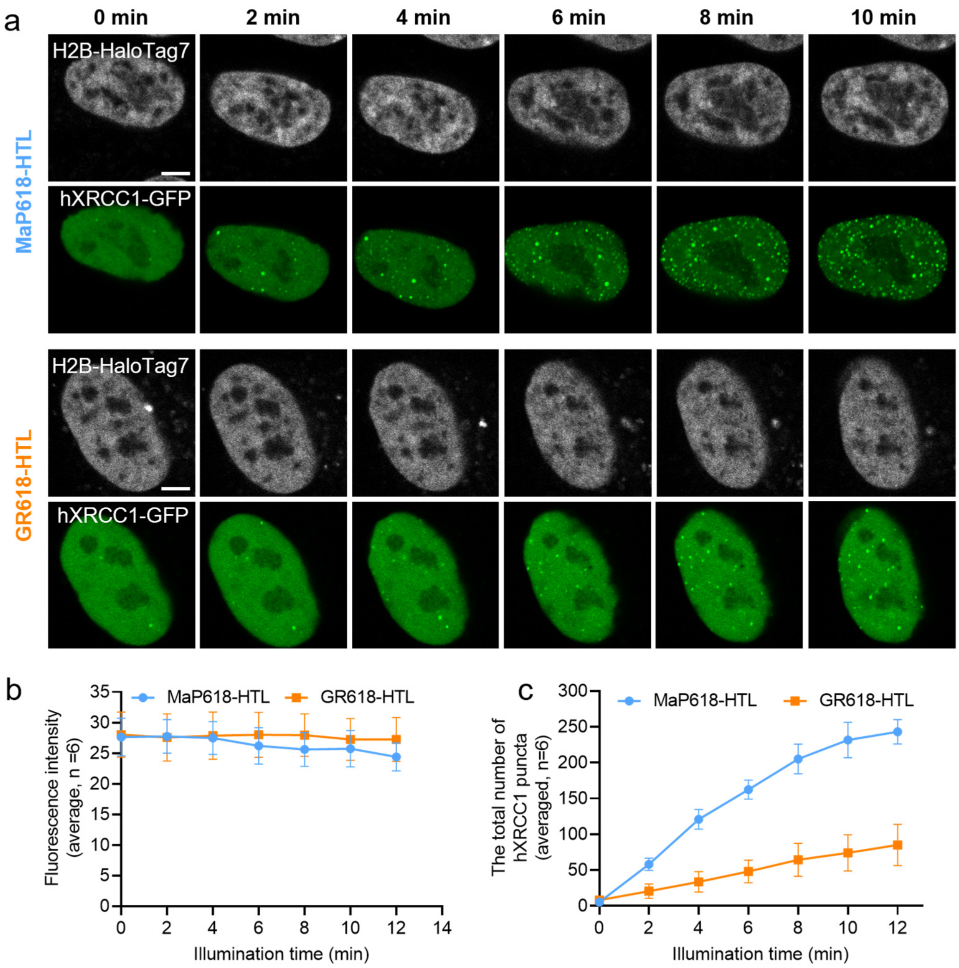


**Figure S20. Phototoxicity measurement of MaP618-HTL and GR618-HTL on U-2 OS EF1α-H2B-HaloTag7-expressing cells.** (a) Live-cell confocal recordings (gray) of U-2 OS EF1α H2B-HaloTag7-expressing cells labeled with **MaP618-HTL** (500 nM in DMEM) or **GR618-HTL** (500 nM in DMEM) and transiently expressing hXRCC1-GFP (green) for 30 min. The two-color confocal images were recorded under continuous 561-nm laser scanning. The snapshots were acquired with 2-min interval. Scale bars: 5 μm. (b) Photobleaching curves of **MaP618-HTL** and **GR618-HTL** on U-2 OS H2B-HaloTag7 expressing cells in time-lapse recordings. Data points represent average bleaching curves of six cells from five independent biological repeats. Error bars indicate the standard error of the mean (SEM). (c) Semi-quantitative analysis of cellular phototoxicity analysis of **GR618-HTL** and **MaP618-HTL** of U-2 OS hXRCC1-GFP cells, as measured by counting the total number of hXRCC1-GFP puncta. Data points represent averaged hXRCC1-GFP number of six cells from five independent experiments. Error bars indicate standard error of the mean. Frame number = 7; Frame rate = 2 min/frame; Duration time = 12 min.

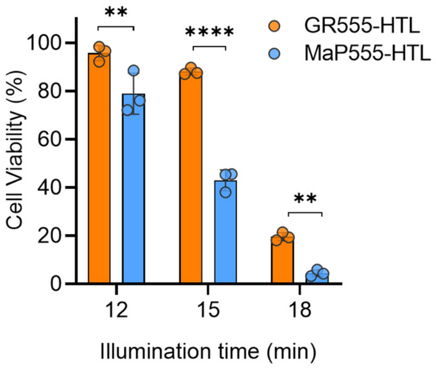


**Figure S21. Phototoxicity measurement of MaP555-HTL and GR555-HTL on HeLa pCMV-mEGFP-HaloTag7-PDGFR$^{tmb}$ cells.** Viability of **MaP555-HTL** and **GR5555-HTL**-stained (500 nM, 30 min) HeLa pCMV-mEGFP-HaloTag7-PDGFR$^{tmb}$ cells after green LED light illumination (532 nm, 2.6 W/cm$^2$), measured by Calcein AM staining. Bars indicate the mean of at least 1500 individual cells from three independent biological replicates. Error bars indicate standard deviation. Significance was determined using Two-way ANOVA followed by Sidak's multiple comparisons test. $P$ = ** <1.0 ×10$^{-2}$ (1.2 ×10$^{-3}$; 2.7 ×10$^{-3}$); **** < 1.0 ×10$^{-4}$.

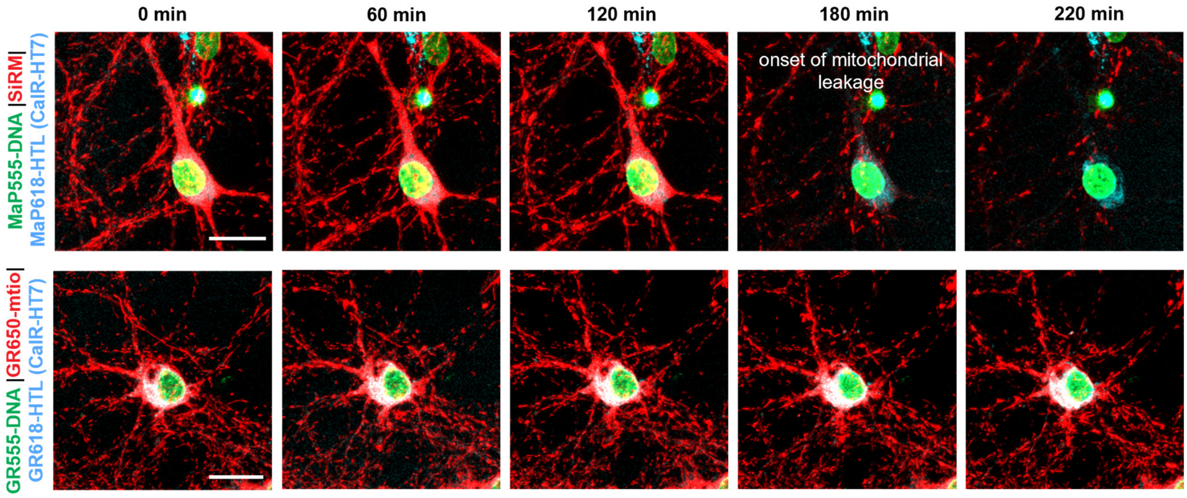


**Figure S22. Three-color time-lapse confocal microscopy recording of rat hippocampal neurons expressing CalR-HaloTag7-KDEL(CalR-HT7).** Live cultured neurons (10 DIV) were labeled with Gentle Rhodamine probes (**GR555-DNA**, 2 µM / **GR618-HTL**, 500 nM / **GR650M**, 50 nM) or established probes (**MaP555-DNA**, 500 nM / **MaP618-HTL,** 500 nM / **SiRM**, 50 nM) for 2h at 37 °C. Confocal recordings of 12 z-stacks per channel at a frame rate of 90 sec/frame were performed for 4 h. Neurons stained with gentle rhodamines showed no signs of phototoxicity and stable mitochondrial signal for the time of recording. Scale bars = 20 µm.

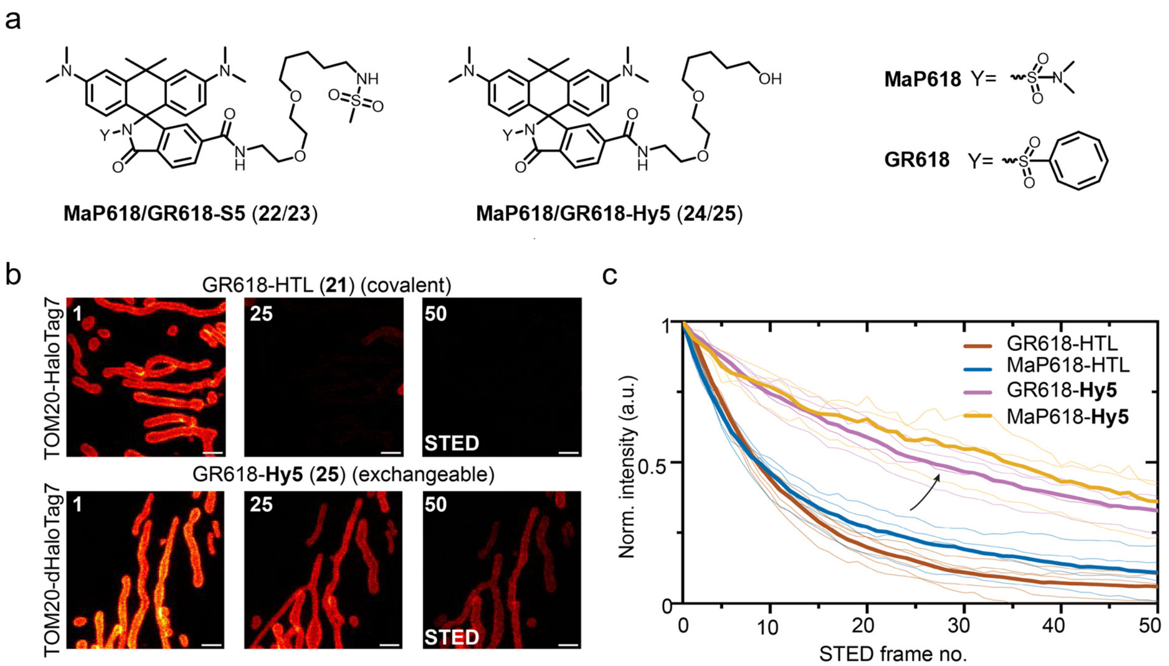


**Figure S23. Time-lapse STED imaging showcasing different photobleaching behavior of MaP/GR618 covalently conjugated to HaloTag (HTL) or its exchangeable counterpart (Hy5)[7].** (a) Chemical structures of compound **22-23**: **MaP618/GR618** derivates coupled to exchangeable HaloTag Ligands (**S5**, **Hy5**). (b) Time-lapse STED imaging showcasing different photobleaching behavior of **MaP/GR618** covalently conjugated to HaloTag (HTL) or its exchangeable counterpart (Hy5) of U-2 OS mitochondria outer membrane (TOM20-HaloTag7 or -dHaloTag7) labeled with **GR618-(x) HTLs** over 50 consecutive frames in a 10 × 10 µm ROI using **MaP618/GR618-HTL, -Hy5**. Frame numbers are indicated in the top left corner. Scale bars: 1 µm. (c) Bleaching curves (thick lines: mean value, thin lines: individual experiments) for at least 4 image series (n ≥ 4) from similar bleaching experiments as shown in b.

**Table S1. Absolute singlet oxygen yields of Rhodamine-COT derivatives in acetonitrile.**

| Compound | Absolute singlet oxygen yield ($\Phi_{\Delta(^1O_2)}$) Units: X $10^{-4}$ (Mean ± SD) | Reference compound |
|---|---|---|
| **1** (TMRM) | 218 ± 5 | TMRE[11,16] |
| **2** (GR555M) | 21.5 ± 0.2 | TMRE[11,16] |
| **3** | 187 ± 2 | TMRE[11,16] |
| **4** | 190 ± 4 | TMRE[16] |
| **5** | 120 ± 2 | TMRE[11,16] |
| **6** (Rho123) | 290 ± 7 | Rho123 in MeOH[16] |
| **7** (GR510M) | 35.1 ± 0.5 | Rhodamine 123 (Rho123)[16] |
| **8** (SiRM) | 32.0 ± 0.6 | Methylene blue (MB)[16] |
| **9** (GR650M) | 10.9 ± 0.3 | Methylene blue (MB)[16] |
| **10** ($JF_{549}$) | 302 ± 8 | TMRE[11,16] |
| **11** ($JF_{549}$-COT) | 20.2 ± 0.4 | TMRE[11,16] |
| **12** (Rho101) | 637 ± 11 | TMRE[11,16] |
| **13** (Rho101-COT) | 81.0 ± 0.9 | TMRE[11,16] |
| **14** (MaP555-DNA) | 140 ± 5 | **1** |
| **15** (GR555-DNA) | 16.9 ± 0.2 | **1** |
| **16** (MaP555-Actin) | 44.0 ± 0.9 | **1** |
| **17** (GR555-Actin) | 7.5 ± 0.2 | **1** |
| **18** (MaP555-HTL) | 321 ± 6 | **1** |
| **19** (GR555-HTL) | 32.2 ± 1.2 | **1** |
| **20** (MaP618-HTL) | 77.8 ± 1.6 | **1** |
| **21** (GR618-HTL) | 19.3 ± 0.4 | **1** |

**Table S2. Photophysical properties of mitochondrial dyes (Rhodamine-COT esters).**

| Compound | $\lambda_{ext}$ [nm] | $\lambda_{em}$ [nm] | Φ |
|---|---|---|---|
| **1** (TMRM) | 551[a] / 552[b] | 574[a] / 580[b] | 0.33[a] / 0.30[b] |
| **2** (GR555M) | 553[a] / 556[b] | 575[a] / 582[b] | 0.51[a] / 0.20[b] |
| **3** | 555[a] / 555[b] | 578[a] / 582[b] | 0.51[a] / 0.20[b] |
| **4** | 556[a] / 556[b] | 580[a] / 587[b] | 0.49[a] / 0.10[b] |
| **5** | 554[a] / 556[b] | 575[a] / 577[b] | 0.17[a] / 0.11[b] |
| **6** (Rho123) | 501[a] / 500[b] | 526[a] / 527[b] | 0.53[a] / 0.54[b] |
| **7** (GR510M) | 503[a] / 503[b] | 528[a] / 529[b] | 0.13[a] / 0.20[b] |
| **8** (SiRM) | 649[a] / 648[b] | 667[a] / 671[b] | 0.52[a] / 0.26[b] |
| **9** (GR650M) | 650[a] / 651[b] | 667[a] / 673[b] | 0.64[a] / 0.22[b] |
| **10** ($JF_{549}$) | 552[a] / 548[b] | 576[a] / 574[b] | n.d. |
| **11 ($JF_{549}$-COT)** | 554[a] / 556[b] | 579[a] / 582[b] | n.d. |
| **12** (Rho101) | 576[a] / 576[b] | 599[a] / 600[b] | n.d. |
| **13** (Rho101-COT) | 578[a] / 582[b] | 604[a] / 608[b] | n.d. |

[a]acetonitrile (ACN), [b]PBS (pH 7.4)

**Table S3. Photophysical properties of Rhodamine-COT derivates for protein and DNA labeling.**

| | Dye | $\lambda_{ext}$ [nm] | $\lambda_{em}$ [nm] | ε [$M^{-1}cm^{-1}$] | Φ | $F/F_0$ [d] | $D_{50}$ [e] | τ [ns][f] |
|---|---|---|---|---|---|---|---|---|
| HaloTag Ligand | **TMR** | 554[a] / 556[b] | 577[a] / 578[b] | 60,000[a] / 84,000[b] | 0.43[a] / 0.56[b] | 2 ± 1 | 15 ± 1.1 | n.d. |
| | **MaP555** (**18**) | 554[a]/ 558[b] | 576[a] / 578[b] | 22,000[a] / 85,000[b] | 0.38[a] / 0.52[b] | 15 ± 1 | 66 ± 2 | 2.3 ± 0.1 |
| | **GR555** (**19**) | 550[a] / 552[b,c] | 574[a] / 576[b,c] | 46,000[a] / 75,000[b] / 110,000[c] | 0.37[a] / 0.58[b] / 0.66[c] | 5 ± 1 | 40 ± 1 | 2.2 ± 0.1 |
| | **MaP618** (**20**) | 618[b] / 620[c] | 642[b] / 636[c] | *7,200*[b,g] / *107,000*[c,g] | 0.64[b] /0.63[c] | 335 ± 16 | >75 | 2.9 ± 0.1 |
| | **GR618** (**21**) | 618[b] / 620[c] | 640[b] / 636[c] | 16,200[b] / 48,000[c] | 0.70[b] / 0.69[c] | 41 ± 3 | >75 | 3.3 ± 0.1 |
| DNA | **MaP555** (**14**) | 552[a] / 554[h] | 582[a] / 584[h] | n.d. | 0.23[a] / 0.35[h] | 4 ± 1 | n.d. | n.d. |
| | **GR555** (**15**) | 554[a] / 556[h] | 584[a] / 586[h] | | 0.21[a] / 0.37[h] | 8 ± 1 | | |

[a]HEPES buffer (pH 7.3), [b]bound to HaloTag7, [c]0.1% SDS in HEPES buffer (pH 7.3), [d]ratio of max. fluorescence intensities in the presence and absence of HaloTag7; average data and standard deviation from three replicates, [e]dioxane-$H_2O$ mixture (v/v: 90/10 –10/90); average data and standard deviation from three replicates, [f] average fluorescence lifetimes (τ) using 1,000 photons per pixel from nuclear signal of U-2 OS FlpIn TRex[TM] H2B-HaloTag7-T2A-mEGFP expressing cells (n≥3 images, mean value and standard deviation). [g]taken from L. Wang *et a.l*[4] [h]bound to hpDNA.

**Table S4. Plasmids used in this study**

| Gene | Plasmid | Source |
|---|---|---|
| 6×His-firefly luciferase-HaloTag7 | T7 | |
| 10xHis-TEV-HaloTag7 | pET51b (+) | Addgene #167267 |
| H2B-HaloTag7-T2A-mEGFP | pcDNA5/FR/TO | Addgene #187070 |
| TOM20-HaloTag7 | pcDNA5/FR/TO | Addgene #169330 |
| TOM20-dHaloTag7 | pcDNA5/FR/TO | |
| hXRCC1-AcGFP | pRetroQ | |
| pCMV-mEGFP-HaloTag7- $PDGFR^{tmb}$ | pCMV | |
| PB-pCMV-mEGFP-HaloTag7- $PDGFR^{tmb}$ | pCMV | |
| Super PiggyBac Transposase | T7 | System Bioscience PB200PA-1 |
| CalR-HaloTag7-KDEL-WPRE-SV40 | pGP-AAV-hSyn1 | |
| pCMV-Voltron | pCMV | |

**Table S5. Crystal data and structure refinement for compound 2.**

| | |
|---|---|
| Empirical formula | $C_{33}H_{31}N_2O_3PF_6$ |
| Formula weight | 648.57 |
| Temperature/K | 293(2) |
| Crystal system | monoclinic |
| Space group | C2/c |
| a/Å | 28.7845(11) |
| b/Å | 15.4247(6) |
| c/Å | 14.2626(5) |
| α/° | 90 |
| β/° | 102.379(3) |
| γ/° | 90 |
| Volume/Å$^3$ | 6185.3(4) |
| Z | 8 |
| $\rho_{calc}$ (g/cm$^3$) | 1.393 |
| μ/mm$^{-1}$ | 0.162 |
| F(000) | 2688.0 |
| Crystal size/mm$^3$ | 0.18 × 0.12 × 0.06 |
| Radiation | MoKα (λ = 0.71073) |
| 2Θ range for data collection/° | 4.408 to 50.054 |
| Index ranges | -34 ≤ h ≤ 34, -18 ≤ k ≤ 18, -16 ≤ l ≤ 16 |
| Reflections collected | 17966 |
| Independent reflections | 5421 [$R_{int}$ = 0.0162, $R_{sigma}$ = 0.0194] |
| Data/restraints/parameters | 5421/97/460 |
| Goodness-of-fit on F$^2$ | 1.064 |
| Final R indexes [I>=2σ (I)] | $R_1$ = 0.0712, w$R_2$ = 0.2164 |
| Final R indexes [all data] | $R_1$ = 0.0847, w$R_2$ = 0.2316 |
| Largest diff. peak/hole / e Å$^{-3}$ | 0.52/-0.33 |

**Table S6. Fractional Atomic Coordinates (×10$^4$) and Equivalent Isotropic Displacement Parameters (Å$^2$×10$^3$) for compound 2. $U_{eq}$ is defined as 1/3 of the trace of the orthogonalized $U_{IJ}$ tensor.**

| Atom | *x* | *y* | *z* | U(eq) |
|---|---|---|---|---|
| P1 | 8681.1(3) | 3050.4(6) | 8343.6(7) | 69.1(3) |
| F1A | 8172.1(9) | 2971(2) | 7687(2) | 109.1(10) |
| F1B | 8968(7) | 2699(14) | 9347(11) | 109.1(10) |
| F2A | 8689.7(11) | 4047(2) | 8044(3) | 127.0(13) |
| F2B | 9174(6) | 3168(14) | 8052(18) | 127.0(13) |
| F3A | 8912.7(11) | 2801(3) | 7483(2) | 119.4(11) |
| F3B | 8703(8) | 2090(10) | 7909(15) | 119.4(11) |
| F4A | 9212.3(10) | 3132.7(17) | 8976(2) | 99.3(9) |
| F4B | 8696(8) | 3953(10) | 8751(16) | 99.3(9) |
| F5A | 8486.5(17) | 3316(3) | 9234(3) | 154.1(16) |
| F5B | 8217(6) | 2828(17) | 8667(17) | 154.1(16) |
| F6A | 8673.5(12) | 2069(2) | 8633(3) | 118.9(11) |
| F6B | 8422(8) | 3347(13) | 7340(11) | 118.9(11) |
| O1 | 4810.7(6) | 2585.7(11) | 6107.2(13) | 41.8(4) |
| O2 | 6171.8(7) | 2526.9(13) | 5323.5(14) | 54.1(5) |
| O3A | 6955.9(17) | 2699(3) | 5498(4) | 74.1(15) |
| O3B | 6956(7) | 2344(14) | 5524(17) | 74.1(15) |
| N1 | 4532.5(9) | -426.2(15) | 5938.5(17) | 49.4(6) |
| N2 | 4930.2(10) | 5638.5(16) | 6224.9(19) | 60.9(7) |
| C1 | 4022.5(11) | -270(2) | 5617(3) | 62.8(8) |
| C2 | 4677.5(12) | -1333.5(18) | 6086(2) | 57.0(8) |
| C3 | 4842.9(9) | 235.4(16) | 6135.6(17) | 41.1(6) |
| C4 | 5339.2(10) | 92.7(17) | 6500.5(18) | 44.9(6) |
| C5 | 5644.7(9) | 763.4(17) | 6740.2(17) | 41.9(6) |
| C6 | 5491.1(9) | 1648.4(16) | 6647.3(17) | 38.6(6) |
| C7 | 4998.0(9) | 1767.1(15) | 6249.9(17) | 37.4(6) |
| C8 | 4685.1(9) | 1103.3(17) | 5993.0(18) | 40.8(6) |
| C9 | 5786.8(9) | 2366.5(17) | 6891.6(18) | 40.3(6) |
| C10 | 5587.4(9) | 3205.3(17) | 6758.1(18) | 41.8(6) |
| C11 | 5848.3(10) | 3985.6(18) | 6980(2) | 49.3(7) |
| C12 | 5636.4(11) | 4770.1(19) | 6799(2) | 53.0(7) |
| C13 | 5139.4(11) | 4857.7(18) | 6392(2) | 51.2(7) |
| C14 | 4871.9(10) | 4084.9(18) | 6170.7(19) | 47.0(6) |
| C15 | 5093.8(9) | 3297.5(16) | 6350.7(18) | 41.0(6) |
| C16 | 4425.8(14) | 5723(2) | 5833(3) | 69.9(9) |
| C17 | 5210.7(16) | 6435(2) | 6352(3) | 78.8(11) |
| C18 | 6298.8(9) | 2230.8(19) | 7357(2) | 47.1(6) |
| C19 | 6412.9(11) | 2027(3) | 8325(2) | 70.4(10) |
| C20 | 6882.5(14) | 1847(3) | 8791(3) | 87.3(13) |
| C21 | 7240.1(12) | 1897(3) | 8296(3) | 81.2(12) |
| C22 | 7138.5(11) | 2124(3) | 7353(3) | 67.3(9) |
| C23 | 6670.4(9) | 2280.7(19) | 6862(2) | 49.8(7) |

| Atom | *x* | *y* | *z* | U(eq) |
|---|---|---|---|---|
| C24 | 6559.4(10) | 2485.7(18) | 5827(2) | 49.1(7) |
| C25A | 6889(3) | 2947(4) | 4492(5) | 76.2(18) |
| C26A | 6820.2(17) | 3913(4) | 4362(3) | 68.3(12) |
| C27A | 6815(2) | 4465(4) | 5075(4) | 87.8(15) |
| C28A | 6701(3) | 5389(4) | 5001(5) | 109(2) |
| C29A | 6901(3) | 5978(5) | 4537(5) | 132(3) |
| C30A | 7282(3) | 5851(6) | 4006(6) | 151(4) |
| C31A | 7302(3) | 5312(6) | 3320(6) | 130(3) |
| C32A | 6935(2) | 4704(4) | 2893(4) | 99.2(19) |
| C33A | 6720.8(18) | 4122(3) | 3343(4) | 74.9(13) |
| C25B | 6931(14) | 2668(15) | 4580(19) | 76.2(18) |
| C26B | 7063(6) | 3620(11) | 4503(10) | 68.3(12) |
| C27B | 7392(6) | 3965(11) | 5321(10) | 87.8(15) |
| C28B | 7355(7) | 4615(11) | 5775(13) | 109(2) |
| C29B | 6981(9) | 5247(13) | 5616(15) | 132(3) |
| C30B | 6821(10) | 5607(16) | 4846(17) | 151(4) |
| C31B | 6958(8) | 5526(14) | 3948(14) | 130(3) |
| C32B | 7008(7) | 4866(13) | 3500(13) | 99.2(19) |
| C33B | 6918(6) | 3994(10) | 3742(12) | 74.9(13) |

**Table S7. Anisotropic Displacement Parameters (Å$^2$×10$^3$) for compound 2. The Anisotropic displacement factor exponent takes the form: $-2\pi^2[h^2a^{*2}U_{11}+2hka^*b^*U_{12}+…]$.**

| Atom | $U_{11}$ | $U_{22}$ | $U_{33}$ | $U_{23}$ | $U_{13}$ | $U_{12}$ |
|---|---|---|---|---|---|---|
| P1 | 67.0(6) | 71.7(6) | 71.3(6) | -10.6(4) | 20.7(5) | -27.2(4) |
| F1A | 60.6(15) | 119(2) | 145(3) | -20.4(18) | 15.0(15) | -33.8(15) |
| F1B | 60.6(15) | 119(2) | 145(3) | -20.4(18) | 15.0(15) | -33.8(15) |
| F2A | 94(2) | 84(2) | 183(3) | 30(2) | -14(2) | -29.0(15) |
| F2B | 94(2) | 84(2) | 183(3) | 30(2) | -14(2) | -29.0(15) |
| F3A | 88(2) | 195(3) | 84.0(17) | -5(2) | 36.9(15) | -17(2) |
| F3B | 88(2) | 195(3) | 84.0(17) | -5(2) | 36.9(15) | -17(2) |
| F4A | 99(2) | 81.5(17) | 99.5(19) | -2.7(13) | -17.9(15) | -27.1(14) |
| F4B | 99(2) | 81.5(17) | 99.5(19) | -2.7(13) | -17.9(15) | -27.1(14) |
| F5A | 196(4) | 170(4) | 117(3) | -37(2) | 79(3) | 8(3) |
| F5B | 196(4) | 170(4) | 117(3) | -37(2) | 79(3) | 8(3) |
| F6A | 126(3) | 77.3(19) | 158(3) | 11.9(18) | 40(2) | -34.4(17) |
| F6B | 126(3) | 77.3(19) | 158(3) | 11.9(18) | 40(2) | -34.4(17) |
| O1 | 35.1(9) | 36.3(9) | 55.0(11) | 1.2(7) | 11.5(8) | 1.7(7) |
| O2 | 44.0(11) | 67.4(14) | 52.5(11) | 3.2(9) | 14.1(9) | 0.2(9) |
| O3A | 44.1(12) | 119(5) | 65.2(15) | 23(3) | 25.2(11) | 12(3) |
| O3B | 44.1(12) | 119(5) | 65.2(15) | 23(3) | 25.2(11) | 12(3) |
| N1 | 54.2(14) | 42.2(12) | 54.7(13) | -4.4(10) | 18.2(11) | -3.9(10) |
| N2 | 75.1(18) | 38.7(13) | 70.7(16) | 7.0(11) | 19.3(14) | 5.4(12) |
| C1 | 50.7(17) | 53.8(18) | 84(2) | -10.3(16) | 15.3(15) | -10.9(14) |
| C2 | 71(2) | 40.5(15) | 61.2(17) | -1.4(13) | 19.1(15) | -8.2(13) |
| C3 | 48.5(15) | 40.1(14) | 38.7(12) | -3.1(10) | 18.3(11) | -2.4(11) |
| C4 | 53.7(16) | 38.6(14) | 46.4(14) | 4.0(11) | 19.6(12) | 7.7(12) |
| C5 | 38.8(13) | 45.0(14) | 44.2(13) | 6.3(11) | 14.0(11) | 7.1(11) |
| C6 | 36.2(13) | 43.0(14) | 39.6(12) | 4.9(10) | 14.8(10) | 2.4(10) |
| C7 | 38.4(13) | 36.9(13) | 40.3(12) | 3.2(10) | 15.7(10) | 4.0(10) |
| C8 | 35.9(13) | 44.1(14) | 44.7(13) | -0.7(11) | 13.7(10) | 0.1(11) |
| C9 | 37.1(13) | 46.7(14) | 40.3(13) | 5.3(11) | 15.1(10) | -0.3(11) |
| C10 | 40.8(14) | 43.9(14) | 43.2(13) | 2.6(11) | 14.6(11) | -2.6(11) |
| C11 | 47.3(15) | 50.2(16) | 53.0(15) | -0.2(12) | 16.8(12) | -6.0(12) |
| C12 | 59.4(18) | 46.3(16) | 57.2(16) | -2.4(13) | 21.1(14) | -10.4(13) |
| C13 | 68.8(19) | 40.9(15) | 49.1(15) | 3.9(12) | 24.1(14) | 2.1(13) |
| C14 | 50.3(15) | 44.5(15) | 48.5(15) | 4.3(11) | 15.4(12) | 5.3(12) |
| C15 | 42.8(14) | 39.8(14) | 43.7(13) | 2.2(11) | 16.5(11) | -0.5(11) |
| C16 | 90(3) | 49.3(18) | 73(2) | 11.9(15) | 22.2(19) | 18.2(17) |
| C17 | 106(3) | 41.8(18) | 91(3) | 2.8(17) | 27(2) | -2.5(17) |
| C18 | 37.6(14) | 50.1(15) | 54.5(15) | 9.7(12) | 11.8(11) | -3.4(12) |
| C19 | 45.0(17) | 104(3) | 62.2(19) | 25.9(18) | 11.1(14) | -8.7(17) |
| C20 | 58(2) | 127(4) | 71(2) | 42(2) | 0.9(17) | -6(2) |
| C21 | 41.6(17) | 115(3) | 82(2) | 32(2) | 1.4(16) | 2.4(18) |
| C22 | 36.6(15) | 85(2) | 81(2) | 16.6(18) | 14.4(15) | 7.3(15) |
| C23 | 37.0(14) | 53.4(16) | 60.3(17) | 7.9(13) | 13.3(12) | 1.9(12) |

| Atom | $U_{11}$ | $U_{22}$ | $U_{33}$ | $U_{23}$ | $U_{13}$ | $U_{12}$ |
|---|---|---|---|---|---|---|
| C24 | 39.1(15) | 54.4(16) | 58.1(16) | 1.6(13) | 19.9(13) | 4.7(12) |
| C25A | 56(3) | 121(6) | 58(2) | 22(3) | 27(2) | 16(4) |
| C26A | 52(3) | 95(4) | 61(2) | -5(2) | 18(2) | -14(2) |
| C27A | 91(4) | 94(4) | 73(3) | 1(3) | 4(3) | -24(3) |
| C28A | 133(6) | 84(4) | 104(5) | -20(4) | 14(4) | -21(4) |
| C29A | 191(8) | 100(5) | 88(4) | 4(4) | -8(5) | -41(5) |
| C30A | 164(7) | 155(7) | 117(6) | 33(5) | -10(5) | -82(6) |
| C31A | 117(5) | 159(7) | 109(5) | 38(5) | 12(4) | -45(5) |
| C32A | 92(4) | 131(5) | 72(3) | 27(4) | 11(3) | -15(3) |
| C33A | 62(3) | 86(3) | 72(3) | 4(3) | 4(2) | 4(3) |
| C25B | 56(3) | 121(6) | 58(2) | 22(3) | 27(2) | 16(4) |
| C26B | 52(3) | 95(4) | 61(2) | -5(2) | 18(2) | -14(2) |
| C27B | 91(4) | 94(4) | 73(3) | 1(3) | 4(3) | -24(3) |
| C28B | 133(6) | 84(4) | 104(5) | -20(4) | 14(4) | -21(4) |
| C29B | 191(8) | 100(5) | 88(4) | 4(4) | -8(5) | -41(5) |
| C30B | 164(7) | 155(7) | 117(6) | 33(5) | -10(5) | -82(6) |
| C31B | 117(5) | 159(7) | 109(5) | 38(5) | 12(4) | -45(5) |
| C32B | 92(4) | 131(5) | 72(3) | 27(4) | 11(3) | -15(3) |
| C33B | 62(3) | 86(3) | 72(3) | 4(3) | 4(2) | 4(3) |

**Table S8. Bond Lengths for compound 2.**

| Atom | Atom | Length/Å | Atom | Atom | Length/Å |
|---|---|---|---|---|---|
| P1 | F1A | 1.566(3) | C9 | C18 | 1.496(4) |
| P1 | F1B | 1.586(13) | C10 | C11 | 1.418(4) |
| P1 | F2A | 1.597(3) | C10 | C15 | 1.422(4) |
| P1 | F2B | 1.573(13) | C11 | C12 | 1.355(4) |
| P1 | F3A | 1.566(3) | C12 | C13 | 1.430(4) |
| P1 | F3B | 1.612(13) | C13 | C14 | 1.417(4) |
| P1 | F4A | 1.605(3) | C14 | C15 | 1.371(4) |
| P1 | F4B | 1.506(12) | C18 | C19 | 1.385(4) |
| P1 | F5A | 1.549(3) | C18 | C23 | 1.405(4) |
| P1 | F5B | 1.542(13) | C19 | C20 | 1.400(5) |
| P1 | F6A | 1.570(3) | C20 | C21 | 1.370(6) |
| P1 | F6B | 1.535(13) | C21 | C22 | 1.359(5) |
| O1 | C7 | 1.371(3) | C22 | C23 | 1.400(4) |
| O1 | C15 | 1.367(3) | C23 | C24 | 1.476(4) |
| O2 | C24 | 1.192(3) | C25A | C26A | 1.509(7) |
| O3A | C24 | 1.363(5) | C26A | C27A | 1.328(7) |
| O3A | C25A | 1.457(6) | C26A | C33A | 1.456(7) |
| O3B | C24 | 1.321(14) | C27A | C28A | 1.461(8) |
| O3B | C25B | 1.423(18) | C28A | C29A | 1.324(9) |
| N1 | C1 | 1.461(4) | C29A | C30A | 1.473(10) |
| N1 | C2 | 1.463(4) | C30A | C31A | 1.295(10) |
| N1 | C3 | 1.346(3) | C31A | C32A | 1.447(9) |
| N2 | C13 | 1.345(4) | C32A | C33A | 1.329(7) |
| N2 | C16 | 1.445(4) | C25B | C26B | 1.527(17) |
| N2 | C17 | 1.460(4) | C26B | C27B | 1.438(18) |
| C3 | C4 | 1.429(4) | C26B | C33B | 1.220(18) |
| C3 | C8 | 1.414(4) | C27B | C28B | 1.209(19) |
| C4 | C5 | 1.353(4) | C28B | C29B | 1.435(19) |
| C5 | C6 | 1.432(4) | C29B | C30B | 1.23(2) |
| C6 | C7 | 1.423(3) | C30B | C31B | 1.425(19) |
| C6 | C9 | 1.395(4) | C31B | C32B | 1.23(2) |
| C7 | C8 | 1.361(4) | C32B | C33B | 1.427(17) |
| C9 | C10 | 1.412(4) | | | |

**Table S9. Bond Angles for compound 2.**

| Atom | Atom | Atom | Angle/˚ | Atom | Atom | Atom | Angle/˚ |
|---|---|---|---|---|---|---|---|
| F1A | P1 | F2A | 88.96(17) | C6 | C9 | C18 | 119.3(2) |
| F1A | P1 | F4A | 177.51(17) | C10 | C9 | C18 | 121.4(2) |
| F1A | P1 | F6A | 91.01(18) | C9 | C10 | C11 | 124.6(2) |
| F1B | P1 | F3B | 88.7(9) | C9 | C10 | C15 | 119.2(2) |
| F2A | P1 | F4A | 90.38(15) | C11 | C10 | C15 | 116.2(2) |
| F2B | P1 | F1B | 87.3(9) | C12 | C11 | C10 | 121.4(3) |
| F2B | P1 | F3B | 83.8(9) | C11 | C12 | C13 | 122.1(3) |
| F3A | P1 | F1A | 90.72(18) | N2 | C13 | C12 | 121.8(3) |
| F3A | P1 | F2A | 89.6(2) | N2 | C13 | C14 | 120.8(3) |
| F3A | P1 | F4A | 86.87(17) | C14 | C13 | C12 | 117.3(2) |
| F3A | P1 | F6A | 90.1(2) | C15 | C14 | C13 | 119.6(3) |
| F4B | P1 | F1B | 90.1(9) | O1 | C15 | C10 | 120.8(2) |
| F4B | P1 | F2B | 92.5(9) | O1 | C15 | C14 | 115.8(2) |
| F4B | P1 | F3B | 176.2(10) | C14 | C15 | C10 | 123.4(2) |
| F4B | P1 | F5B | 92.6(10) | C19 | C18 | C9 | 118.4(2) |
| F4B | P1 | F6B | 93.0(9) | C19 | C18 | C23 | 118.1(3) |
| F5A | P1 | F1A | 93.2(2) | C23 | C18 | C9 | 123.6(2) |
| F5A | P1 | F2A | 89.6(2) | C18 | C19 | C20 | 121.1(3) |
| F5A | P1 | F3A | 176.0(2) | C21 | C20 | C19 | 120.0(3) |
| F5A | P1 | F4A | 89.3(2) | C22 | C21 | C20 | 119.9(3) |
| F5A | P1 | F6A | 90.6(2) | C21 | C22 | C23 | 121.3(3) |
| F5B | P1 | F1B | 88.4(9) | C18 | C23 | C24 | 119.3(2) |
| F5B | P1 | F2B | 173.3(11) | C22 | C23 | C18 | 119.5(3) |
| F5B | P1 | F3B | 91.0(9) | C22 | C23 | C24 | 121.1(3) |
| F6A | P1 | F2A | 179.7(2) | O2 | C24 | O3A | 121.7(3) |
| F6A | P1 | F4A | 89.64(17) | O2 | C24 | O3B | 125.1(11) |
| F6B | P1 | F1B | 176.1(11) | O2 | C24 | C23 | 126.0(2) |
| F6B | P1 | F2B | 90.3(9) | O3A | C24 | C23 | 112.2(3) |
| F6B | P1 | F3B | 88.1(9) | O3B | C24 | C23 | 105.9(10) |
| F6B | P1 | F5B | 93.8(10) | O3A | C25A | C26A | 111.5(5) |
| C15 | O1 | C7 | 120.5(2) | C27A | C26A | C25A | 124.2(5) |
| C24 | O3A | C25A | 117.2(6) | C27A | C26A | C33A | 126.1(5) |
| C24 | O3B | C25B | 112(2) | C33A | C26A | C25A | 109.5(4) |
| C1 | N1 | C2 | 116.0(2) | C26A | C27A | C28A | 127.5(5) |
| C3 | N1 | C1 | 121.2(2) | C29A | C28A | C27A | 126.2(8) |
| C3 | N1 | C2 | 122.6(2) | C28A | C29A | C30A | 128.0(8) |
| C13 | N2 | C16 | 121.6(3) | C31A | C30A | C29A | 129.3(8) |
| C13 | N2 | C17 | 121.1(3) | C30A | C31A | C32A | 125.9(8) |
| C16 | N2 | C17 | 117.1(3) | C33A | C32A | C31A | 127.4(6) |
| N1 | C3 | C4 | 121.8(2) | C32A | C33A | C26A | 128.3(5) |
| N1 | C3 | C8 | 120.6(2) | O3B | C25B | C26B | 116(2) |
| C8 | C3 | C4 | 117.6(2) | C27B | C26B | C25B | 115.4(16) |
| C5 | C4 | C3 | 121.3(2) | C33B | C26B | C25B | 118.4(14) |

| Atom | Atom | Atom | Angle/˚ | Atom | Atom | Atom | Angle/˚ |
|---|---|---|---|---|---|---|---|
| C4 | C5 | C6 | 122.3(2) | C33B | C26B | C27B | 125.9(12) |
| C7 | C6 | C5 | 114.9(2) | C28B | C27B | C26B | 128.5(13) |
| C9 | C6 | C5 | 125.0(2) | C27B | C28B | C29B | 128.8(13) |
| C9 | C6 | C7 | 120.0(2) | C30B | C29B | C28B | 125.1(13) |
| O1 | C7 | C6 | 120.3(2) | C29B | C30B | C31B | 129.8(15) |
| C8 | C7 | O1 | 115.9(2) | C32B | C31B | C30B | 128.8(13) |
| C8 | C7 | C6 | 123.8(2) | C31B | C32B | C33B | 127.5(13) |
| C7 | C8 | C3 | 120.0(2) | C26B | C33B | C32B | 128.0(12) |
| C6 | C9 | C10 | 119.0(2) | | | | |

**Table S10. Torsion Angles for compound 2.**

| A | B | C | D | Angle/˚ | A | B | C | D | Angle/˚ |
|---|---|---|---|---|---|---|---|---|---|
| O1 | C7 | C8 | C3 | -178.4(2) | C15 | O1 | C7 | C8 | 179.3(2) |
| O3A | C25A | C26A | C27A | -0.8(10) | C15 | C10 | C11 | C12 | -0.8(4) |
| O3A | C25A | C26A | C33A | -175.9(6) | C16 | N2 | C13 | C12 | -178.6(3) |
| O3B | C25B | C26B | C27B | 28(3) | C16 | N2 | C13 | C14 | 1.2(4) |
| O3B | C25B | C26B | C33B | -159(2) | C17 | N2 | C13 | C12 | 6.9(4) |
| N1 | C3 | C4 | C5 | -176.7(2) | C17 | N2 | C13 | C14 | -173.2(3) |
| N1 | C3 | C8 | C7 | 176.0(2) | C18 | C9 | C10 | C11 | 4.6(4) |
| N2 | C13 | C14 | C15 | -179.8(2) | C18 | C9 | C10 | C15 | -177.2(2) |
| C1 | N1 | C3 | C4 | 176.5(2) | C18 | C19 | C20 | C21 | 1.9(7) |
| C1 | N1 | C3 | C8 | -3.0(4) | C18 | C23 | C24 | O2 | -8.6(5) |
| C2 | N1 | C3 | C4 | 0.3(4) | C18 | C23 | C24 | O3A | 166.4(3) |
| C2 | N1 | C3 | C8 | -179.2(2) | C18 | C23 | C24 | O3B | -169.6(12) |
| C3 | C4 | C5 | C6 | 0.2(4) | C19 | C18 | C23 | C22 | 0.1(5) |
| C4 | C3 | C8 | C7 | -3.5(3) | C19 | C18 | C23 | C24 | 179.2(3) |
| C4 | C5 | C6 | C7 | -2.4(3) | C19 | C20 | C21 | C22 | 0.3(7) |
| C4 | C5 | C6 | C9 | 179.2(2) | C20 | C21 | C22 | C23 | -2.3(7) |
| C5 | C6 | C7 | O1 | -178.6(2) | C21 | C22 | C23 | C18 | 2.1(5) |
| C5 | C6 | C7 | C8 | 1.6(3) | C21 | C22 | C23 | C24 | -177.0(4) |
| C5 | C6 | C9 | C10 | 179.7(2) | C22 | C23 | C24 | O2 | 170.5(3) |
| C5 | C6 | C9 | C18 | -5.0(4) | C22 | C23 | C24 | O3A | -14.5(5) |
| C6 | C7 | C8 | C3 | 1.4(4) | C22 | C23 | C24 | O3B | 9.5(12) |
| C6 | C9 | C10 | C11 | 179.8(2) | C23 | C18 | C19 | C20 | -2.0(6) |
| C6 | C9 | C10 | C15 | -2.0(4) | C24 | O3A | C25A | C26A | 91.0(7) |
| C6 | C9 | C18 | C19 | -77.7(4) | C24 | O3B | C25B | C26B | 86(3) |
| C6 | C9 | C18 | C23 | 100.8(3) | C25A | O3A | C24 | O2 | -2.4(6) |
| C7 | O1 | C15 | C10 | -0.1(3) | C25A | O3A | C24 | C23 | -177.7(4) |
| C7 | O1 | C15 | C14 | 179.5(2) | C25A | C26A | C27A | C28A | -173.0(7) |
| C7 | C6 | C9 | C10 | 1.4(3) | C25A | C26A | C33A | C32A | -128.2(7) |
| C7 | C6 | C9 | C18 | 176.7(2) | C26A | C27A | C28A | C29A | -53.6(10) |
| C8 | C3 | C4 | C5 | 2.8(4) | C27A | C26A | C33A | C32A | 56.8(8) |
| C9 | C6 | C7 | O1 | -0.2(3) | C27A | C28A | C29A | C30A | -0.9(12) |
| C9 | C6 | C7 | C8 | -179.9(2) | C28A | C29A | C30A | C31A | 54.0(13) |
| C9 | C10 | C11 | C12 | 177.5(2) | C29A | C30A | C31A | C32A | 1.6(14) |
| C9 | C10 | C15 | O1 | 1.4(4) | C30A | C31A | C32A | C33A | -53.0(12) |
| C9 | C10 | C15 | C14 | -178.2(2) | C31A | C32A | C33A | C26A | -4.0(11) |
| C9 | C18 | C19 | C20 | 176.6(4) | C33A | C26A | C27A | C28A | 1.3(9) |
| C9 | C18 | C23 | C22 | -178.5(3) | C25B | O3B | C24 | O2 | 30(2) |
| C9 | C18 | C23 | C24 | 0.6(4) | C25B | O3B | C24 | C23 | -168.9(17) |
| C10 | C9 | C18 | C19 | 97.5(3) | C25B | C26B | C27B | C28B | -129(2) |
| C10 | C9 | C18 | C23 | -83.9(4) | C25B | C26B | C33B | C32B | -177.8(19) |
| C10 | C11 | C12 | C13 | 1.0(4) | C26B | C27B | C28B | C29B | -2(3) |
| C11 | C10 | C15 | O1 | 179.7(2) | C27B | C26B | C33B | C32B | -5(3) |

| A | B | C | D | Angle/° | A | B | C | D | Angle/° |
|---|---|---|---|---|---|---|---|---|---|
| C11 | C10 | C15 | C14 | 0.2(4) | C27B | C28B | C29B | C30B | -51(3) |
| C11 | C12 | C13 | N2 | 179.2(3) | C28B | C29B | C30B | C31B | -2(3) |
| C11 | C12 | C13 | C14 | -0.6(4) | C29B | C30B | C31B | C32B | 51(3) |
| C12 | C13 | C14 | C15 | 0.0(4) | C30B | C31B | C32B | C33B | 4(3) |
| C13 | C14 | C15 | O1 | -179.4(2) | C31B | C32B | C33B | C26B | -52(3) |
| C13 | C14 | C15 | C10 | 0.2(4) | C33B | C26B | C27B | C28B | 58(3) |
| C15 | O1 | C7 | C6 | -0.5(3) | | | | | |

**Table S11. Hydrogen Atom Coordinates (Å×10[4]) and Isotropic Displacement Parameters (Å[2]×10[3]) for compound 2.**

| Atom | *x* | *y* | *z* | U(eq) |
|---|---|---|---|---|
| H1A | 3961.61 | -0.58 | 4996.06 | 94 |
| H1B | 3915.74 | 103.83 | 6065.8 | 94 |
| H1C | 3855.01 | -811.83 | 5577.82 | 94 |
| H2A | 4678.76 | -1497.78 | 6735.88 | 85 |
| H2B | 4990.86 | -1404.44 | 5965.69 | 85 |
| H2C | 4457.96 | -1693.88 | 5653.58 | 85 |
| H4 | 5454.58 | -471.97 | 6574.7 | 54 |
| H5 | 5965.58 | 645.63 | 6973.4 | 50 |
| H8 | 4367.56 | 1220.45 | 5723.68 | 49 |
| H11 | 6171.53 | 3958.21 | 7254.63 | 59 |
| H12 | 5819.99 | 5267.85 | 6944.74 | 64 |
| H14 | 4547.61 | 4111.99 | 5904.89 | 56 |
| H16A | 4355.89 | 5537.25 | 5175.62 | 105 |
| H16B | 4333.29 | 6318.33 | 5867.69 | 105 |
| H16C | 4253.23 | 5370.28 | 6196.37 | 105 |
| H17A | 5352.53 | 6511.81 | 7019.61 | 118 |
| H17B | 5008.03 | 6919.37 | 6127.14 | 118 |
| H17C | 5456.12 | 6396.81 | 5991.6 | 118 |
| H19 | 6173.59 | 2008.12 | 8671.52 | 84 |
| H20 | 6951.65 | 1694.88 | 9436.93 | 105 |
| H21 | 7551.95 | 1774.77 | 8603.68 | 97 |
| H22 | 7384.82 | 2176.11 | 7028.39 | 81 |
| H25A | 6613.29 | 2648.42 | 4122.39 | 91 |
| H25B | 7164.62 | 2769.23 | 4250.27 | 91 |
| H27A | 6892.59 | 4237.06 | 5692.03 | 105 |
| H28A | 6466.73 | 5582.57 | 5308.27 | 130 |
| H29A | 6786.43 | 6540.88 | 4548.3 | 159 |
| H30A | 7545.09 | 6210.12 | 4189.27 | 181 |
| H31A | 7578.3 | 5315.66 | 3079.9 | 157 |
| H32A | 6836.52 | 4722.75 | 2228.62 | 119 |
| H33A | 6476.64 | 3807.88 | 2958.21 | 90 |
| H25C | 6609.54 | 2583.33 | 4212.29 | 91 |
| H25D | 7140.32 | 2320.8 | 4281.34 | 91 |
| H27B | 7669.37 | 3647.33 | 5528.13 | 105 |
| H28B | 7603.73 | 4718.35 | 6298.54 | 130 |
| H29B | 6849.82 | 5396.35 | 6135.32 | 159 |
| H30B | 6571.62 | 5990.45 | 4850.33 | 181 |
| H31B | 7017.74 | 6044.63 | 3661.92 | 157 |
| H32B | 7118.4 | 4943.09 | 2938.53 | 119 |
| H33B | 6725.03 | 3670.3 | 3261.39 | 90 |

**Table S12. Atomic Occupancy for compound 2.**

| Atom | *Occupancy* | Atom | *Occupancy* | Atom | *Occupancy* |
|---|---|---|---|---|---|
| F1A | 0.905(2) | F1B | 0.095(2) | F2A | 0.905(2) |
| F2B | 0.095(2) | F3A | 0.905(2) | F3B | 0.095(2) |
| F4A | 0.905(2) | F4B | 0.095(2) | F5A | 0.905(2) |
| F5B | 0.095(2) | F6A | 0.905(2) | F6B | 0.095(2) |
| O3A | 0.777(3) | O3B | 0.223(3) | C25A | 0.777(3) |
| H25A | 0.777(3) | H25B | 0.777(3) | C26A | 0.777(3) |
| C27A | 0.777(3) | H27A | 0.777(3) | C28A | 0.777(3) |
| H28A | 0.777(3) | C29A | 0.777(3) | H29A | 0.777(3) |
| C30A | 0.777(3) | H30A | 0.777(3) | C31A | 0.777(3) |
| H31A | 0.777(3) | C32A | 0.777(3) | H32A | 0.777(3) |
| C33A | 0.777(3) | H33A | 0.777(3) | C25B | 0.223(3) |
| H25C | 0.223(3) | H25D | 0.223(3) | C26B | 0.223(3) |
| C27B | 0.223(3) | H27B | 0.223(3) | C28B | 0.223(3) |
| H28B | 0.223(3) | C29B | 0.223(3) | H29B | 0.223(3) |
| C30B | 0.223(3) | H30B | 0.223(3) | C31B | 0.223(3) |
| H31B | 0.223(3) | C32B | 0.223(3) | H32B | 0.223(3) |
| C33B | 0.223(3) | H33B | 0.223(3) | | |

### Chemical Synthesis and Characterization of New Compounds

### General Information

Unless otherwise mentioned, all reactions were carried out under a nitrogen atmosphere with dry solvents under anhydrous conditions. All the chemicals were purchased at the highest commercial quality and used without further purification unless otherwise stated. Reactions were monitored by Thin Layer Chromatography on silica plates (GF254 by Yantai Chemicals or POLYGRAM® SIL G/UV254, Roth) using UV light as a visualizing agent and an ethanolic solution of phosphomolybdic acid and cerium sulfate, and heat as developing agents or by LC/MS (4.6 mm × 150 mm 5 µm $C_{18}$ column; 2 µL injection; 10-95% or 50-95% $CH_3CN/H_2O$, linear-gradient, with constant 0.1% v/v TFA additive; 20 min run; 1 mL/min flow; ESI; positive ion mode; UV detection at 254 nm).
Flash column chromatography uses silica gel (200-300 mesh) supplied by Tsingtao Haiyang Chemicals (Qingdao, Shandong, China) or was carried out on a Biotage (IsoleraTM One) flash system equipped and pre-packed silica columns (SiliaSep™ Flash Cartridges, 40 – 63 µm, 60 Å). Preparative reversed-phase high-performance liquid chromatography (RP-HPLC) was carried out on an UltiMate 3000 system (Thermo Fisher Scientific) equipped with a $C_{18}$ column (5 µm, 21.2 × 250 mm, Supelco) and a 2998 PDA. Solvents: 0.1% TFA in MiliQ® water (dd$H_2O$), MeCN.
Analytical LC/MS was performed on a LCMS2020 (Shimadzu) connected to a Nexera UHPLC system with a $C_{18}$ 1.7 µm, 50 × 2.1 mm (ACQUITY UPLC BEH, Waters) column or LCMSD iQ (Agilent Technologies) connected 1290 Infinity II system with RRHD Eclipse Plus $C_{18}$ column 1.8 µm, 50 × 2.1 mm (Agilent Technologies). Typical gradient was from 10% dd$H_2O$ water w. 0.1% formic acid to 90% MeCN within 6 min with 0.5 mL/min flow or 5% dd$H_2O$ water w. 0.1% formic acid to 95% MeCN w. 0.1% formic acid within 6 min with 0.4 mL/min flow.
NMR spectra were recorded on Bruker Advance 400 ($^1$H 400 MHz, $^{13}$C 101 MHz) and are calibrated using residual undeuterated solvent ($CDCl_3$ at 7.26 ppm $^1$H NMR, 77.16 ppm $^{13}$C NMR; $CD_3OD$ at 3.31 ppm $^1$H NMR, 49.00 ppm $^{13}$C NMR; DMSO-$d_6$ at 2.50 ppm $^1$H NMR, 39.52 ppm $^{13}$C NMR). Data for $^1$H NMR spectra are reported as follows: chemical shift (δ ppm), multiplicity (s = singlet, d = doublet, t = triplet, q = quartet, dd = doublet of doublets, dt = triplet of doublets, m = multiplet, br = broad), coupling constant (Hz), integration. Data for $^{13}$C NMR are reported by chemical shift (δ ppm).

### Synthetic Procedure and Characterization

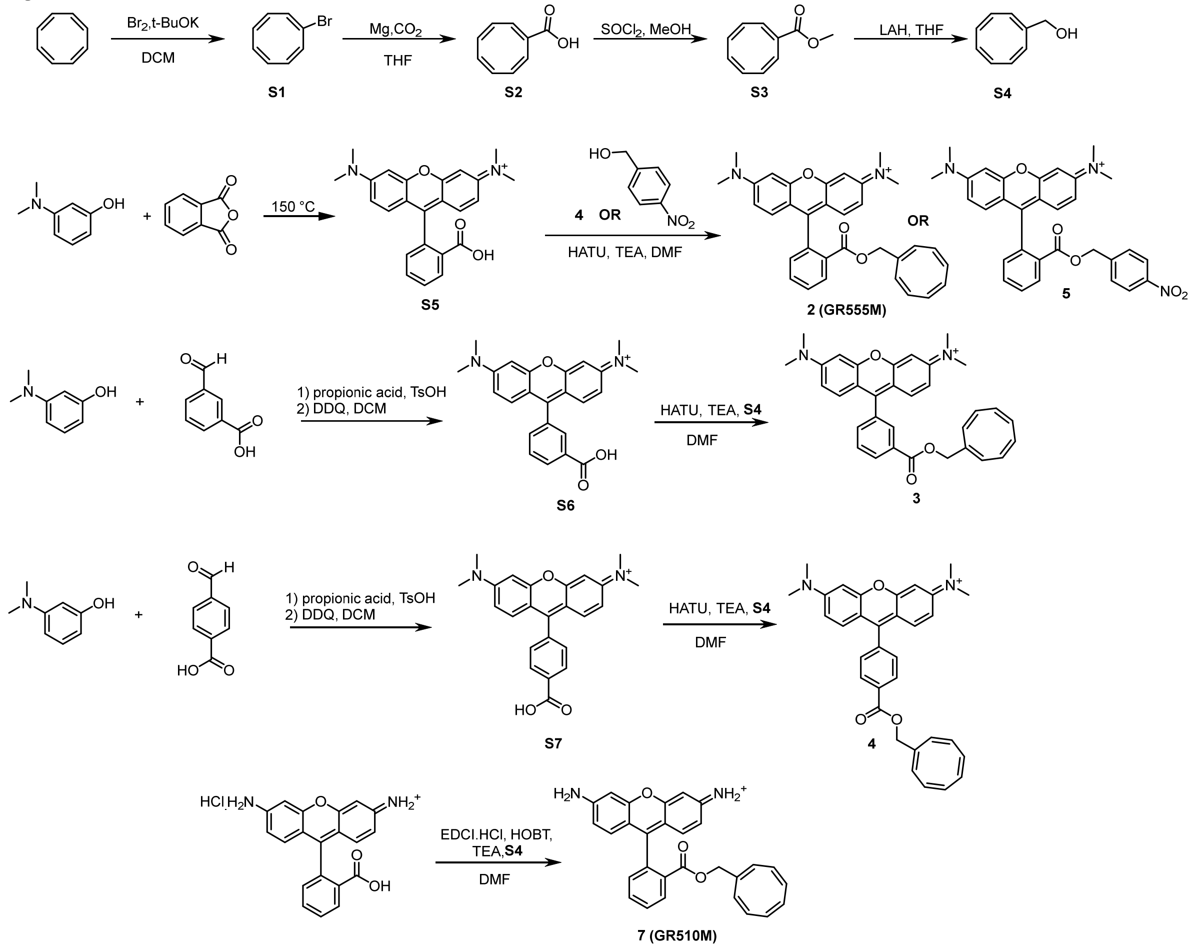


**Supplementary Scheme S1**. Synthetic route of GR mitochondria probes Compound **1-7**.

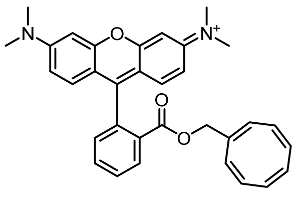

Synthesis of compound **S1** to **S5** were referred to in previously reported work[17,18].

**Compound 2 (GR555M)**: Compound **S5** (80 mg, 0.21 mmol), HATU (118 mg, 0.310 mmol), and TEA (42.5 mg, 0.42 mmol) were dissolved in 3.0 mL DMF. Compound **S4** (41.6 mg, 0.310 mmol) was added. The reaction mixture was stirred at r.t. for 24 h, concentrated under reduce pressure and purified by silica gel chromatography column, eluting with DCM: MeOH=20:1 to afford compound **1** (36 mg, impure). 36 mg compound **2** was dissolved in DCM (2.5 mL) and added dropwise into MTBE (25 mL) slowly, the solid was precipitated and filtered. The filtered cake was collected and dried under reduce pressure to obtain compound **2** (20 mg, 19%) as a red solid.

**HRMS** (m/z): calculated for $C_{33}H_{31}N_2O_3^+[M]^+$ 503.2329, found 503.2414.

**$^1$H NMR** (400 MHz, Chloroform-d) δ 8.27 (d, *J* = 7.6 Hz, 1H), 7.81 (t, *J* = 7.5 Hz, 1H), 7.74 (t, *J* = 7.5 Hz, 1H), 7.35 (d, *J* = 7.7 Hz, 1H), 7.09 7.09 (m, 2H), 6.87 (m, 4H), 5.74 (m, 7H), 4.39 (s, 2H), 3.31 (s, 12H).

**$^{13}$C NMR** (75 MHz, C Chloroform-d) δ 165.93, 160.07, 158.46, 158.37, 138.78, 134.35, 133.90, 132.71, 132.02, 131.86, 131.44, 131.29, 131.19, 131.13, 115.33, 114.50, 97.22, 68.93, 41.28, 30.28.

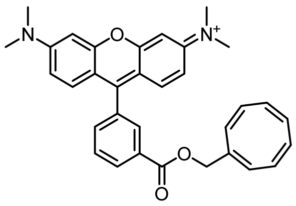

Synthesis of compound **S6** was carried out similar as previously reported[19]. The NMR is in accordance with published result, see reference[20].

**Compound 3**: Compound **S6** (40 mg, 0.10 mmol), HATU (58.7 mg, 0.15 mmol), and TEA (20.8 mg, 0.206 mmol) were dissolved in 4.0 mL DMF. Compound **S4** (16.6 mg, 0.12 mmol) was added. The reaction mixture was stirred at r.t. for 18 h, concentrated under reduce and purified by silica gel chromatography column, eluting with DCM: MeOH=30:1 to obtain compound **3** (33 mg, impure). The crude compound was dissolved in DCM (2.0 mL) and added dropwise into $Et_2O$ (20 mL) slowly, the solid was precipitated and filtered. The filtered cake was collected, dried under reduce pressure to afford the compound **3** (20 mg, 39%) as a red solid.

**HRMS** (m/z): calculated for $C_{33}H_{31}N_2O_3^+[M]^+$ 503.2329, found 503.2410.

**$^1$H NMR** (300 MHz, Acetonitrile-$d_3$) δ 8.27 (d, *J* = 7.9Hz, 1H), 8.01 (s, 1H), 7.79 (t, *J* = 7.7 Hz, 1H), 7.66 (d, *J* = 7.2Hz, 1H), 7.25 (d, *J* = 9.5 Hz, 2H), 6.94 (dd, *J* = 9.5, 2.4 Hz, 2H), 6.79 (d, *J* = 2.4 Hz, 2H), 6.06 – 5.68 (m, 7H), 4.75 (s, 2H), 3.24 (s, 12H).

**$^{13}$C NMR** (75 MHz, Acetonitrile-$d_3$) δ 166.05, 158.58, 158.22, 157.31, 139.48, 134.80, 134.27, 133.49, 133.13, 132.77, 132.69, 132.21, 132.00, 131.92, 131.78, 131.73, 131.08, 130.99, 130.37, 115.28, 114.25, 97.19, 68.59, 41.29.

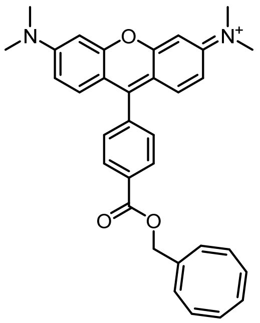

Synthesis of compound **S7** was referred to in previously reported work [19].

**Compound 4**: Compound **S7** (24.5 mg, 0.06 mmol), HATU (36.1 mg, 0.10 mmol), and TEA (12.8 mg, 0.13 mmol) were dissolved in 2.0 mL DMF. Compound **S4** (10.2 mg, 0.08 mmol) was added. The reaction mixture was stirred at r.t. for 18 h before concentrated under reduce pressure to obtain the residue, which was purified by silica gel chromatography column, eluting with DCM: MeOH= from 100:1 to 30:1 to obtain the compound **3** (20 mg, impure). The 20 mg compound **4** was dissolved in DCM (1.5 mL) and added dropwise into $Et_2O$ (20 mL) slowly, the solid was precipitated and filtered. The filtered cake was collected, dried under reduce pressure to afford the compound **4** (15 mg, 47%) as a red solid.

**HRMS** (m/z): calculated for $C_{33}H_{31}N_2O_3^+[M]^+$ 503.2329, found 503.2408.

**$^1$H NMR** (400 MHz, Chloroform-d) δ 8.27 (d, *J* = 7.9 Hz, 1H), 7.48 (d, *J* = 8.2 Hz, 1H), 7.33 – 7.19 (m, 1H), 6.92 (dd, *J* = 9.4, 2.2 Hz, 1H), 6.84 (d, *J* = 2.3 Hz, 1H), 5.92 (m,7H), 4.81 (s, 2H), 3.32 (s, 12H).

**$^{13}$C NMR** (75 MHz, Chloroform-d) δ 165.35, 157.79, 157.44, 156.44, 138.30, 136.55, 133.69, 132.46, 132.11, 132.09, 131.89, 131.37, 131.21, 131.02, 130.70, 130.64, 130.21, 129.75, 114.63, 113.23, 97.06, 68.19, 53.59, 41.08.

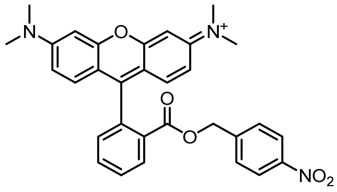


**Compound 5**: Compound **S5** (100 mg, 0.26 mmol), HATU (147 mg, 0.35 mmol), and TEA (78.3 mg, 0.77 mmol) were dissolved in 5.0 mL DMF. P-nitrobenzyl alcohol (CAS: 619-73-8; 51.4 mg, 0.35 mmol) was added. The reaction mixture was stirred at r.t. for 18 h, concentrated under reduce pressure and purified by silica gel chromatography column, eluting with DCM: MeOH=50:1 to afford the compound **5** (40 mg, 41%) as a red solid.

**HRMS** (m/z): calculated for $C_{31}H_{28}N_3O_5^+$[M]$^+$ 522.2023, found 522.2070.

**$^1$H NMR** (400 MHz, DMSO-*d*$_6$) δ 8.31 (dd, *J* = 7.7, 1.5 Hz, 1H), 8.04 (d, *J* = 8.7 Hz, 2H), 7.92 (td, *J* = 7.5, 1.5 Hz, 1H), 7.87 (td, *J* = 7.6, 1.5 Hz, 1H), 7.49 (dd, *J* = 7.4, 1.4 Hz, 1H), 7.22(d, *J* = 8.7 Hz, 2H), 7.04 (dd, *J* = 9.4, 2.4 Hz, 2H), 6.97 (d, *J* = 9.4 Hz, 2H), 6.80 (d, *J* = 2.3 Hz, 2H), 5.08 (s, 2H), 3.26 (s, 12H).

**$^{13}$C NMR** (101 MHz, DMSO-*d*$_6$) δ 164.68, 157.53, 156.77, 156.56, 147.05, 142.22, 133.29, 132.81, 130.96, 130.56, 130.48, 130.38, 129.43, 129.14, 123.36, 114.70, 112.86, 96.03, 65.53, 40.44, 40.44, 38.24.

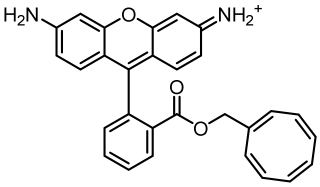


**Compound 7 (GR510M)**: Rhodamine 110 (CAS: 13558-31-1; 100 mg, 0.27 mmol), EDCI.HCl (CAS: 7084-11-9; 67.9 mg, 0.35 mmol), HOBT (47.9 mg, 0.35 mmol), and TEA (110 mg, 1.09 mmol) were dissolved in 5.0 mL DMF. Compound **S4** (48.3 mg, 0.33 mmol) was added. The reaction mixture was stirred at r.t. for 24 h before concentrated under reduce pressure to obtain the residue, which was purified by reversed-HPLC (0.5% HCl aqueous, v/v) to obtain the compound **7** (8.0 mg, 7%) as a red solid.

**HRMS** (m/z): calculated for $C_{29}H_{23}N_2O_3^+$[M]$^+$ 447.1703, found 447.1876.

**$^1$H NMR** (400 MHz, Methanol-d4) δ 8.29 (dd, *J* = 7.7, 1.4 Hz, 1H), 7.86 (td, *J* = 7.5, 1.5 Hz, 1H), 7.81 (td, *J* = 7.6, 1.5 Hz, 1H), 7.43 (dd, *J* = 7.7, 1.4 Hz, 1H), 7.05 (d, *J* = 8.9 Hz, 2H), 6.86 – 6.79 (m, 4H), 4.34 (s, 2H).

**$^{13}$C NMR** (101 MHz, Methanol-d4) δ 166.44, 161.54, 161.33, 161.10, 159.71, 138.89, 135.22, 134.75, 134.06, 132.87, 132.79, 132.34, 132.21, 131.73, 131.55, 118.00, 117.89, 114.95, 114.88, 98.56, 98.39, 69.57.

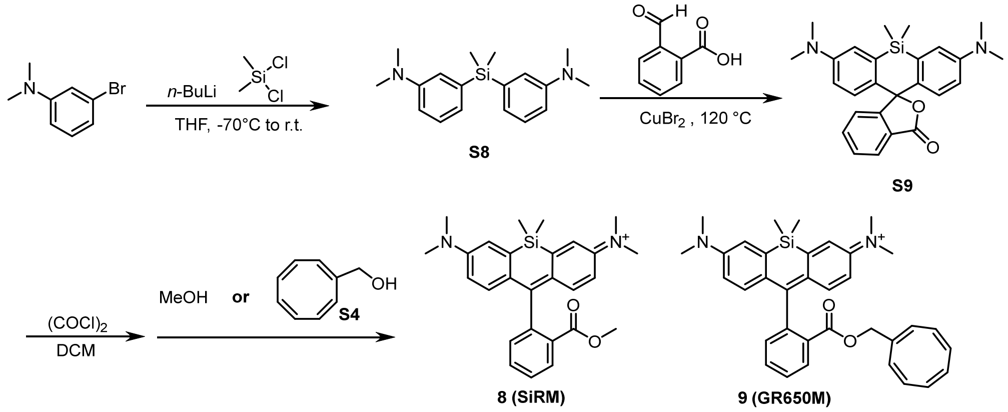


**Supplementary Scheme S2.** Synthetic route of Compound **6 (SiRM)** and **7 (GR650M)**.

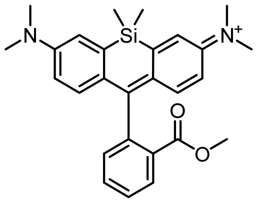

**Compound 8 (SiRM)**: Compound **S9** was synthesized according to J. Yang *et al.*[21] 5.5 mg **S9** (0.01 mmol) were dissolved in DCM (1.0 mL), oxalyl chloride (74 mL, 0.58 mmol, 50 eq.) was added at 0 °C and the solution was left stirring for 8h at r.t. Subsequently, MeOH (1 mL) was added at 0 °C under nitrogen. The mixture was allowed to warm up to r.t. and left stirring for 30 min, concentrated under reduce pressure and purified by prep-TLC (DCM: MeOH=10:1) to obtain the compound **8** (1.8 mg, 34%) as a blue solid.

**HRMS** (m/z): calculated for $C_{27}H_{31}N_2O_2Si^+[M]^+$ 443.2149 found 443.2296.

**$^1$H NMR** (400 MHz, Methanol-d4) δ 8.25 (d, *J* = 7.8 Hz, 1H), 7.78 (t, *J* = 7.4 Hz, 1H), 7.71 (t, *J* = 7.6 Hz, 1H), 7.34 (d, *J* = 2.8 Hz, 2H), 7.30 (dd, *J* = 7.5, 1.4 Hz, 1H), 6.97 (d, *J* = 9.6 Hz, 2H), 6.73 (dd, *J* = 9.6, 2.8 Hz, 2H), 3.62 (s, 3H), 3.33 (s, 12H), 0.64 (s, 3H), 0.60 (s, 3H).

**$^{13}$C NMR** (101 MHz, Methanol-d4) δ 172.32, 166.97, 155.59, 149.44, 142.13, 141.90, 133.46, 131.94, 131.84, 131.49, 130.31, 129.04, 121.71, 114.90, 52.73, 40.90, -0.64, -1.74.

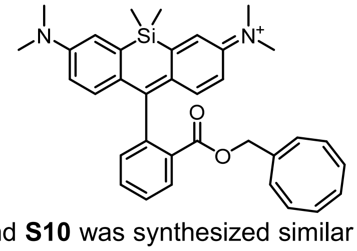

**Compound 9 (GR650M)**: Compound **S10** was synthesized similar as compound **S9**. Instead of MeOH, compound **S4** (1.90 mg, 0.01mmol) was added at 0 °C under nitrogen. The mixture was stirred at r.t. for 3 h, concentrated under reduce pressure and purified by prep-TLC (DCM: MeOH=10:1) to obtain the compound **9** (1.2 mg, 18%) as a blue solid.

**HRMS** (m/z): calculated for $C_{27}H_{31}N_2O_2Si^+[M]^+$ 545.2619 found 545.2684.

**$^1$H NMR** (400 MHz, Acetonitrile-d3) δ 8.20 (d, *J* = 7.8 Hz, 1H), 7.76 (t, *J* = 7.4 Hz, 1H), 7.70 (t, *J* = 7.5 Hz, 1H), 7.29 (d, *J* = 7.6 Hz, 1H), 7.27 (d, *J* = 2.8 Hz, 2H), 6.94 (d, *J* = 9.6 Hz, 2H), 6.65 (dd, *J* = 9.6, 2.7 Hz, 2H), 5.83 – 5.48 (m, 7H), 4.41 (s, 2H), 3.27 (s, 12H), 0.59 (s, 6H).

$^{13}$**C NMR** (101 MHz, Methanol-d4) δ 171.90, 166.41, 155.63, 149.56, 142.11, 141.80, 139.21, 134.19, 133.45, 132.95, 132.88, 132.64, 132.06, 131.97, 131.86, 131.75, 131.67, 130.81, 130.33, 129.11, 121.78, 114.98, 68.71, 40.84, -0.84, -1.41.

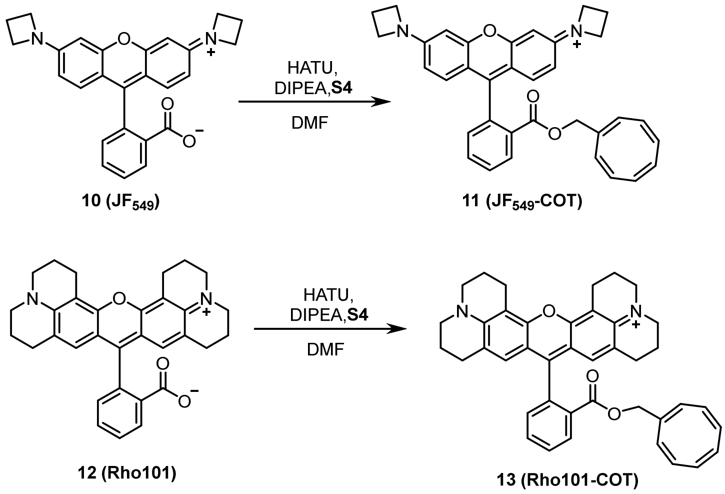


**Supplementary Scheme S3.** Synthetic route of Compound **11 ($JF_{549}$-COT)** and **13 (Rho101-COT)**.

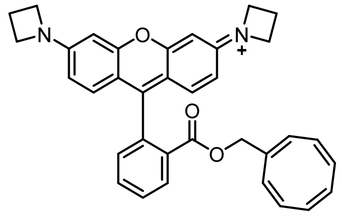

**Compound 11 ($JF_{549}$-COT)**: Compound **10** was synthesized according to J.Grimm *et.al.*[22]. Compound **10** (10 mg, 0.02 mmol), HATU (15.2 mg, 0.04 mmol), and DIPEA (15.5 mg, 0.12 mmol) were dissolved in 2.0 mL DMF. Compound **S4** (10.7 mg, 0.08 mmol) was added and stirred at r.t. for 24 h. The mixture was concentrated under reduce pressure and purified by prep-HPLC to obtain compound **11** (2.6 mg, 21%) as a purple solid.
**LC/MS** (5% to 95% MeCN/$H_2O$, 6 min, 0.4 mL/min) 3.14 min, 527.4 m/z, >99%.
**HRMS** (m/z): calculated for $C_{35}H_{31}N_2O_3^+[M]^+$ 527.2329, found 527.2371.
$^{1}$**H NMR** (400 MHz, Methanol-d4) δ 8.27 (dd, *J* = 7.8, 1.1 Hz, 1H), 7.86 (td, *J* = 7.5, 1.4 Hz, 1H), 7.80 (td, *J* = 7.6, 1.3 Hz, 1H), 7.45 – 7.38 (m, 1H), 7.06 (d, *J* = 9.2 Hz, 2H), 6.62 (dd, *J* = 9.2, 2.1 Hz, 2H), 6.56 (d, *J* = 2.1 Hz, 2H), 5.77 – 5.32 (m, 7H), 4.35 – 4.27 (m,10H), 2.57 (p, *J* = 7.6 Hz, 4H).

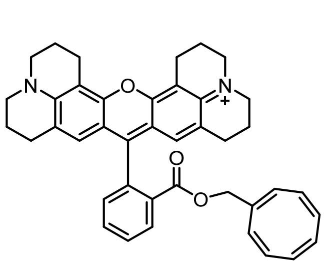

**Compound 13 (Rho101-COT)**: Compound **12** (CAS: 116450-56-7; 10 mg, 0.02 mmol), HATU (15.2 mg, 0.04 mmol), and DIPEA (15.5 mg, 0.12 mmol) were dissolved in 2.0 mL DMF. Compound **S4** (10.7 mg, 0.08 mmol) was added and stirred at r.t. for 3 h. The mixture was concentrated under reduce pressure and purified by prep-HPLC to obtain compound **13** (1.6 mg, 15.9%) as a purple solid.

**LC/MS** (5% to 95% MeCN/$H_2O$, 6 min, 0.4 mL/min) 3.50 min, 607.4 m/z, >99%.
**HRMS** (m/z): calculated for $C_{41}H_{39}N_2O_3^+[M]^+$ 607.2955, found 607.2880.
**$^1$H NMR** (400 MHz, Methanol-d4) δ 8.24 (dd, *J* = 7.7, 1.2 Hz, 1H), 7.83 (td, *J* = 7.5, 1.4 Hz, 1H), 7.78 (td, *J* = 7.6, 1.3 Hz, 1H), 7.39 – 7.35 (m, 1H), 6.63 (s, 2H), 5.69 – 5.02 (m, 7H), 4.29 (s, 2H), 3.59 – 3.54 (m, 4H), 3.54 – 3.50 (m, 4H), 3.10 (q, *J* = 6.1 Hz, 4H), 2.72 – 2.65 (m, 4H), 2.12 (p, *J* = 6.4 Hz, 4H), 1.95 (p, *J* = 6.4 Hz, 4H).

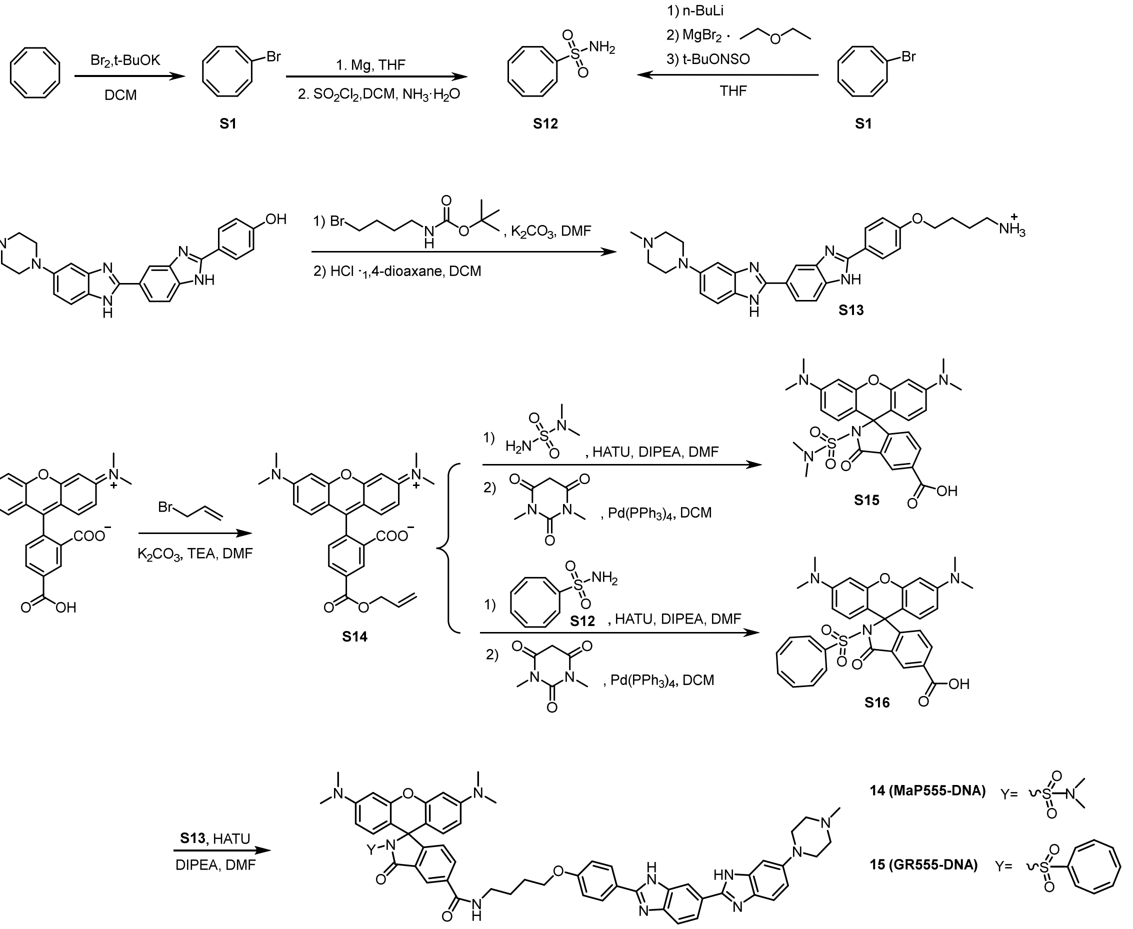


**Supplementary Scheme S4.** Synthetic route of Compound **14** (**MaP555-DNA)** and **15 (GR555-DNA)**.

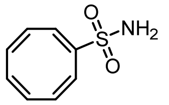

**Compound S12**: A solution of **S1** (100 mg, 0.55 mmol) in THF (1 mL) was added dropwise into a mixture of magnesium ribbon in THF (1 mL) at 0 °C under argon gas. The mixture was stirred at r.t for 2 h until a dark blue-green colored solution was generated indicating formation of the Grignard reagent (compound **S11**). This resulting solution was added dropwise into sulfonyl chloride (CAS: 7791-25-5; 1 mL) diluted with dry DCM (5-6 mL) at −78 °C and the mixture was stirred at −78 °C for 10 min. The reaction was allowed to warmed up to r.t. and stirred at r.t. for another 30 min. After then the solvent and excess sulfonyl chloride were concentrated *in vacuo* to obtain brown oil for next step without further purification. A solution of crude brown oil in dry DCM was added to pre-cooled 7M ammonia in methanol (CAS: 7664-41-7) diluted with dry THF (1 mL). The mixture was stirred at r.t. for 2 h before concentrated under reduce pressure to obtain the residue, which was purified by silica gel chromatography column, eluting with PE: EA= from 20:1 to 3:1 to obtain the compound **S12** (21 mg, 21%) as a brown oil.

Alternative route: 1.6 M n-BuLi (1.7 mL, 3 mmol, 1.1 eq.) was added dropwise to a solution of **S1** (500 mg, 2.7 mmol) in THF (10 mL) at -78 °C under argon gas. The mixture was stirred at 78 °C for 30 min, warmed to -10 °C and magnesium bromide ethyl etherate (776 mg, 3 mmol, 1.1 eq.) in 5 mL THF were added. Meanwhile, N-sulfinyl-O-(tert-butyl) hydroxylamine (t-BuONSO[23], TCI, 500 mg, 3.7 mmol, 1.4 eq.) was cooled to -78 °C and added to the reaction mixture slowly. The reaction was allowed to warm to r.t. and left stirring for 18 h. The solvent was removed under reduced pressure and the crude was purified by flash column purification ($SiO_2$. 0 to 60% EA in n-hexane in 12 CV). Compound **S12** (180 mg, 1.0 mmol, 36 %) as a brown oil.
**$^1$H NMR** (400 MHz, Chloroform-d) δ 6.90 (d, *J* = 3.2 Hz, 1H), 6.20 (dd, *J* = 11.3, 3.0 Hz, 1H), 6.06 (d, *J* = 11.3 Hz, 1H), 6.00 – 5.80 (m, 4H), 4.73 (s, 2H).
**$^{13}$C NMR** (101 MHz, Chloroform-d) δ 143.59, 138.29, 138.23, 134.46, 132.49, 131.47, 128.78, 125.13.

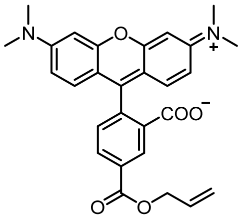

**Compound S14**: 5-carboxytetramethylrhodamine (CAS: 91809-66-4; 100 mg, 0.23 mmol, 1.0 equiv.) was dissolved in DMF (2 mL). $K_2CO_3$ (64 mg, 0.46 mmol, 2.0 equiv.) and $Et_3N$ (65 µL, 0.46 mmol, 2.0 equiv.) were added. The reaction was cooled down in an ice bath and allyl bromide (30 µL, 0.35 mmol, 1.5 equiv.) was added slowly. The reaction mixture was allowed to warm up to r.t. and stirred overnight. Then, the reaction was diluted with water and extracted with DCM. The combined organics were washed with brine, dried with $Na_2SO_4$, filtered, and concentrated *in vacuo* to obtain the residue, which was purified by prep-TLC (DCM: MeOH=10:1) to obtain the compound **S14** (90 mg, 83%) as a red solid.
**HRMS** (m/z): calculated for $C_{28}H_{27}N_2O_5$ [M+H] $^+$ 471.1842 found 471.2082.
**$^1$H NMR** (400 MHz, Methanol-d4) δ 8.93 (d, *J* = 1.8 Hz, 1H), 8.46 (dd, *J* = 7.9, 1.8 Hz, 1H), 7.57 (d, *J* = 8.0 Hz, 1H), 7.14 (d, *J* = 9.5 Hz, 2H), 7.06 (dd, *J* = 9.4, 2.5 Hz, 2H), 6.99 (d, *J* = 2.4 Hz, 2H), 6.22 – 6.08 (m, 1H), 5.49 (dd, *J* = 17.3, 1.6 Hz, 1H), 5.36 (dd, *J* = 10.5, 1.5 Hz, 1H), 4.95 (d, *J* = 5.8 Hz, 1H), 3.31 (s, 12H).
**$^{13}$C NMR** (101 MHz, Methanol-d4) δ 167.08, 166.00, 160.33, 159.02, 158.97, 139.73, 134.26, 133.50, 133.36, 133.28, 133.00, 132.20, 131.90, 119.09, 115.61, 114.62, 97.47, 67.35, 40.94.

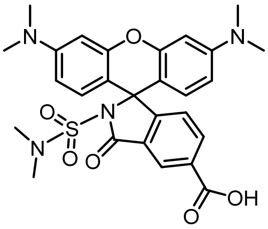

**Compound S15**: Compound **S14** (74 mg, 0.16 mmol, 1.0 equiv.), HATU (120 mg, 0.32 mmol, 2.0 equiv.), and DIPEA (548 µL, 3.28 mmol, 20.0 equiv.) were dissolved in dry DMF (2 mL) and stirred at r.t. for 5 min. Then N, N-dimethylsulfamide (CAS: 3984-14-3; 585 mg, 4.74 mmol, 30.0 equiv.) was added and stirred at r.t. for 36 h. Then, the reaction was diluted with water and extracted with DCM. The combined organics were washed with brine, dried with $Na_2SO_4$, filtered, and concentrated *in vacuo* to obtain the residue. The products were mixed with 1,3-dimethylbarbituric acid (41 mg, 0.26 mmol, 3.0 equiv.), and tetrakis (triphenyl-phosphine) palladium (0) (50 mg, 0.04 mmol, 0.5 equiv.) in DCM (2 mL) and stirred at r.t. for 30 min. The mixture was concentrated under reduce pressure and purified by prep-HPLC to obtain compound **S15** (26 mg, 38%) as a red solid.
**LC/MS** (5% to 95% MeCN/$H_2O$, 6 min, 0.4 mL/min) 3.48 min, 537.3 m/z, >99%.
**HRMS** (m/z): calculated for $C_{27}H_{29}N_4O_6S$ [M+H] $^+$ 537.1730 found 537.1883.

**$^{1}$H NMR** (400 MHz, Methanol-d4) δ 8.53 (d, *J* = 1.2 Hz, 1H), 8.40 (d, *J* = 7.8 Hz, 1H), 7.52 (s, 1H), 7.20 – 6.71 (m, 6H), 3.24 (s, 12H), 2.63 (s, 6H).
**$^{13}$C NMR** (101 MHz, Methanol-*d*$_4$) δ 166.21, 165.65, 156.48, 133.42, 133.04, 132.47, 130.53, 130.31, 130.11, 128.42, 128.17, 113.10, 112.94, 112.86, 96.79, 39.46, 36.83.

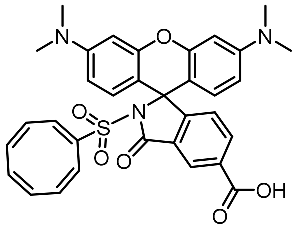


**Compound S16**: Compound **S15** was prepared similar as **S15**, where N, N-dimethylsulfamide was replaced with compound **S12** (43 mg, 0.24 mmol, 1.5 equiv.) to obtain compound **S16** (38 mg, 37%) as a light red solid.
**LC/MS** (5% to 95% MeCN/$H_2O$, 6 min, 0.4 mL/min) 3.55 min, 596.3 m/z, >99%.
**HRMS** (m/z): calculated for $C_{33}H_{30}N_3O_6S$ $[M+H]^+$ 596.1777 found 596.1801.
**$^{1}$H NMR** (400 MHz, Methanol-d4) δ 8.61 (s, 1H), 8.29 (dd, *J* = 8.0, 1.1 Hz, 1H), 7.18 (d, *J* = 8.0 Hz, 1H), 6.67 (d, J = 8.3 Hz, 2H), 6.59 (d, *J* = 7.4 Hz, 4H), 6.43 (s, 1H), 5.95 – 5.42 (m, 6H), 3.05 (s, 12H).
**$^{13}$C NMR** (101 MHz, Methanol-d4) δ 163.55, 163.19, 155.73, 154.82, 143.21, 142.94, 142.86, 138.43, 136.10, 135.78, 133.15, 132.62, 130.66, 129.74, 127.59, 127.27, 126.22, 119.49, 116.58, 111.15, 109.56, 99.23, 40.61, 30.73.

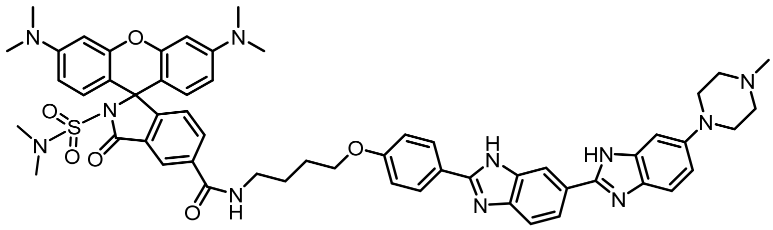

**Compound 14 (MaP555-DNA)**: Compound **S15** (5.0 mg, 0.093 mmol, 1.0 equiv.), HATU (6.2 mg, 0.014 mmol, 1.5 equiv.), compound **S13** (5.0 mg, 0.010 mmol, 1.1 equiv.) and DIPEA (5 µL, 0.028 mmol, 3.0 equiv.) were dissolved in DMF (2 mL) and the mixture was stirred at r.t. for 12 h, concentrated under reduce pressure and purified by prep-HPLC to obtain **MaP555-DNA** (2 mg, 21%) as a red solid.
**LC/MS** (5% to 95% MeCN/$H_2O$, 6 min, 0.4 mL/min) 3.34 min, 508.3 m/z, 99%.
**HRMS** (m/z): calculated for $C_{56}H_{61}N_{11}O_6$ S [M+2H] $^{2+}$ 507.7263 found 507.7233.
**$^1$H NMR** (400 MHz, Methanol-d4) δ 8.39 (s, 2H), 8.19 (dd, J = 8.0, 1.3 Hz, 1H), 8.12 (d, J = 8.8 Hz, 2H), 8.03 (dd, J = 8.5, 1.5 Hz, 1H), 7.88 (d, *J* = 8.5 Hz, 1H), 7.72 (d, *J* = 9.0 Hz, 1H), 7.46 – 7.37 (m, 2H), 7.32 (d, *J* = 1.8 Hz, 1H), 7.20 (d, *J* = 8.9 Hz, 2H), 6.93 (d, *J* = 8.1 Hz, 2H), 6.82 –6.79 (m, 4H), 4.21 (t, *J* = 5.8 Hz, 2H), 3.96 (br, 2H), 3.68 (br, 2H), 3.57 (t, *J* = 6.6 Hz, 2H), 3.48 (t, *J* = 1.6 Hz, 2H), 3.17 (s, 12H), 3.13 (*J* = 1.6 Hz, 2H), 3.02 (s, 3H), 2.62 (s, 6H), 2.02 – 1.89 (m, 4H).

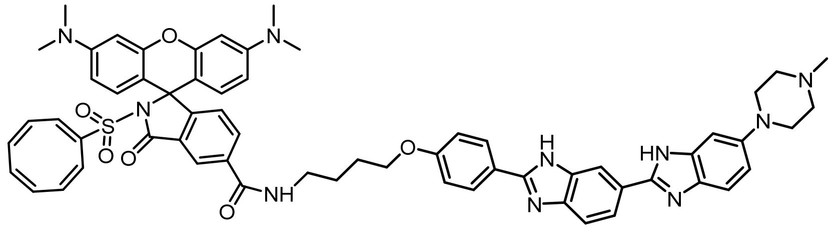

**Compound 15 (GR555-DNA)**: The procedure was same as the synthesis procedure of compound **MaP555-DNA**, where compound **S15** was replaced with compound **S16** (5.6 mg, .093 mmol, 1.0 equiv.) to obtain **GR555-DNA** (3 mg, 30%) as a red solid.
**LC/MS** (5% to 95% MeCN/$H_2O$, 6 min, 0.4 mL/min) 2.99 min, 537.8 m/z, 99%.
**HRMS** (m/z): calculated for $C_{62}H_{62}N_{10}O_6$ S [M+2H] $^{2+}$ 537.2287 found 537.2240.
**$^1$H NMR** (400 MHz, Methanol-d4) δ 8.42 (d, *J* = 1.1 Hz, 1H), 8.36 (s, 1H), 8.20 (dd, *J* = 8.0, 1.3 Hz, 1H), 8.11 (d, *J* = 8.7 Hz, 2H), 8.02 (dd, *J* = 8.6, 0.9 Hz, 1H), 7.86 (d, *J* = 8.5 Hz, 1H), 7.71 (d, *J* = 9.0 Hz, 1H), 7.47 (d, *J* = 8.0 Hz, 1H), 7.38 (dd, *J* = 9.0, 1.7 Hz, 1H), 7.31 (d, *J* = 1.6 Hz, 1H), 7.19 (d, *J* = 8.8 Hz, 2H), 6.98 (d, J = 9.2 Hz, 2H), 6.89 (d, J = 9.3 Hz, 2H), 6.82 (d, J = 1.8 Hz, 2H), 6.64 (s, 1H), 5.98 (d, *J* = 9.3 Hz, 1H), 5.88 (d, *J* = 9.7 Hz, 1H), 5.77 – 5.59 (m, 4H), 4.21 (t, *J* = 5.7 Hz, 2H), 3.94 (br, 2H), 3.67 (br, 2H), 3.58 (t, *J* = 6.4 Hz, 2H), 3.48 (t, *J* = 1.5 Hz, 2H), 3.21 (s, 12H), 3.13 (t, *J* = 1.6 Hz, 2H), 3.02 (s, 3H), 2.02 – 1.87 (m, 4H).

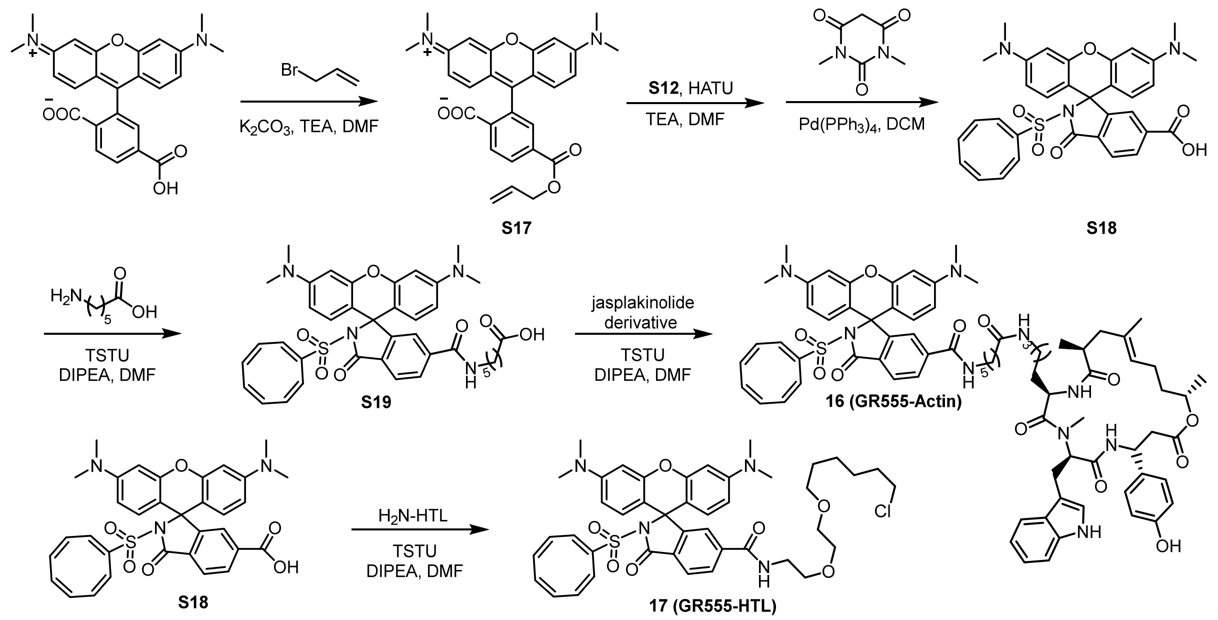


**Supplementary Scheme S5.** Synthetic route of Compound **10 (GR555-Actin)** and **11 (GR555-HTL)**.

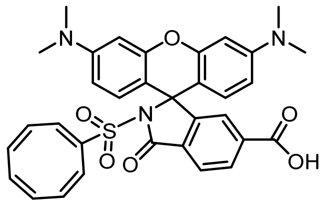

**Compound S18**: Compound S18 was prepared similar as **S14**, where 5-carboxytetramethylrhodamine was replaced with 6-carboxytetramethylrhodamine (100 mg, 0.23 mmol, 1.0 equiv.) to obtain compound **S18** (30 mg, 22%) as a red solid.

**MS** (m/z): calculated for $C_{33}H_{30}N_3O_6S$ [M+H] $^+$ 596.1777 found 596.1801.

**$^1$H NMR** (400 MHz, Acetonitrile-*d*$_3$ + $D_2O$) δ 8.30 (d, *J* = 8.3 Hz, 1H), 7.98 (dd, *J* = 8.1, 1.9 Hz, 1H), 7.92 (s, 1H), 7.05 (d, *J* = 9.4 Hz, 2H), 6.93 (dd, *J* = 9.4, 2.5 Hz, 2H), 6.80 (d, *J* = 2.3 Hz, 2H), 6.64 (s, 1H), 5.99 (d, *J* = 11.2 Hz, 1H), 5.84 (d, *J* = 11.4 Hz, 1H), 5.72 – 5.47 (m, 4H), 3.19 (s, 12H).

**$^{13}$C NMR** (101 MHz, Acetonitrile-d3+ $D_2O$) δ 166.87, 165.05, 157.16, 156.94, 143.82, 143.79, 139.82, 138.25, 136.54, 135.12, 134.59, 132.31, 131.46, 130.91, 130.84, 129.07, 128.56, 128.54, 124.36, 114.54, 113.35, 97.07, 54.56, 40.69.

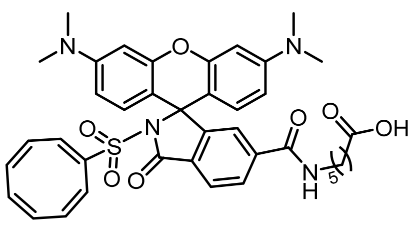

**Compound S19**: Compound **S18** TFA salt (7.1 mg, 10.0 µmol, 1 eq.) was dissolved in DMSO (0.5 ml) and treated with DIPEA (20 µL, 116 µmol, 12 eq.) and TSTU (3 mg, 10.0 µmol, 1 eq.). After 5 minutes, a solution of 6-aminohexanoic acid (2.6 mg, 40 µmol, 4.0 eq) in methanol (0.5 ml) was added. After 15 min the product was purified by preparative HPLC and yielded 6.2 mg (7.5 µmol 75%) as a red solid.

**$^1$H NMR** (400 MHz, Methanol-d4) δ 8.73 (t, *J* = 5.6 Hz, 1H), 8.19 (dd, *J* = 8.1, 1.7 Hz, 1H), 8.02 (d, *J* = 8.1 Hz, 1H), 7.83 (d, *J* = 1.3 Hz, 1H), 7.07 (d, *J* = 9.2 Hz, 2H), 6.99 (dd, *J* = 9.3, 2.2 Hz, 2H), 6.91 (d, *J* = 2.2 Hz, 2H), 6.70 (s, 1H), 6.06 – 5.55 (m, 6H), 3.39 (q, *J* = 6.9 Hz, 2H), 3.27 (s, 12H), 2.29 (t, *J* = 7.4 Hz, 2H), 1.69 – 1.56 (m, 4H), 1.46 – 1.35 (m, 2H).

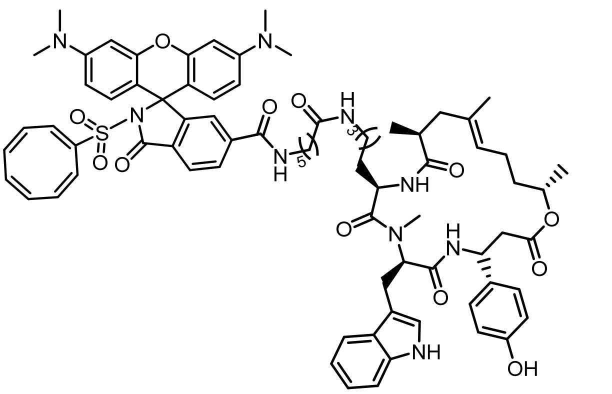

**Compound 17** (**GR555-Actin**): Compound **S19** (5.6 mg, 6.8 µmol, 1 eq.) was dissolved in 0.4 mL DMSO. DIPEA (20 µL, 116 umol, 17 eq.) and TSTU (2.4 mg, 8.2 µmol, 1.2 eq.) were added. After 5 min, Jasplakinolide-amine derivative[8] (6.4 mg, 8.2 umol, 1.2 eq.) dissolved in DMSO (0.2 mL) was added to the reaction. After 1 h purification by prep-HPLC yield compound **16** (**GR555-Actin**, 5.3 mg, 3.4 µmol, 53%) as a red solid.

**LC/MS** (5% to 95% MeCN/$H_2O$, 6 min, 0.4 mL/min) 3.88 min, 683.4 m/z, 98%.

**HRMS** (m/z): calculated for $C_{77}H_{91}N_9O_{12}S$ $[M+2H]^{2+}$ 682.8254 found 682.8216.

**$^1$H NMR** (400 MHz, Methanol-d4) δ 8.75 (t, *J* = 5.6 Hz, 1H), 8.35 (d, *J* = 8.5 Hz, 1H), 8.19 (dd, *J* = 8.1, 1.6 Hz, 1H), 8.00 (d, *J* = 8.1 Hz, 1H), 7.89 – 7.84 (m, 1H), 7.56 (d, *J* = 7.8 Hz, 2H), 7.27 (d, *J* = 8.0 Hz, 1H), 7.10 – 6.86 (m, 12H), 6.72 (d, *J* = 8.6 Hz, 2H), 5.99 (d, *J* = 9.7 Hz, 1H), 5.87 (d, *J* = 9.5 Hz, 1H), 5.74 – 5.51 (m, 5H), 5.22 (td, *J* = 8.9, 3.8 Hz, 1H), 5.01 (t, *J* = 6.8 Hz, 1H), 4.79 – 4.69 (m, 1H), 3.41 (q, *J* = 6.6 Hz, 2H), 3.25 (s, 12H), 3.13 (d, *J* = 8.2 Hz, 2H), 3.07 (s, 3H), 2.98 (dt, *J* = 14.1, 7.2 Hz, 1H), 2.89 (dt, *J* = 13.4, 7.1 Hz, 1H), 2.78 – 2.55 (m, 4H), 2.26 (d, *J* = 13.1 Hz, 1H), 2.17 (t, *J* = 7.3 Hz, 2H), 1.93 – 1.82 (m, 3H), 1.71 – 1.59 (m, 5H), 1.60 – 1.54 (m, 1H), 1.51 (s, 3H), 1.46 – 1.34 (m, 2H), 1.16 (t, *J* = 5.5 Hz, 5H), 1.06 (d, *J* = 6.8 Hz, 3H), 1.03 – 0.84 (m, 3H), 0.78 (s, 1H).

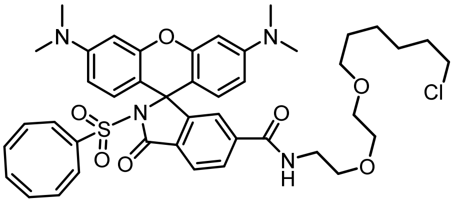

**Compound 19** (**GR555-HTL**): Compound **S18** (10 mg, 0.017 mmol, 1.0 equiv.), HATU (8.9 mg, 0.020 mmol,1.2 equiv.), $H_2N$-HaloTag Ligand[24] (5.8 mg, 0.026 mmol, 1.5 equiv.), and DIPEA (8 µL, 0.051 mmol, 3.0 equiv.) were dissolved in DMF. The mixture was stirred at r.t. for 12 h. Purified by prep-HPLC yielded **GR555-HTL** (3 mg, 22%) as a red solid.

**HRMS** (m/z): calculated for $C_{43}H_{50}ClN_4O_7S$ $[M+H]^+$ 801.3089 found 801.3034

**LC/MS** (5% to 95% MeCN/$H_2O$, 6 min, 0.4 mL/min) 4.22 min, 801.3 m/z, 99%.

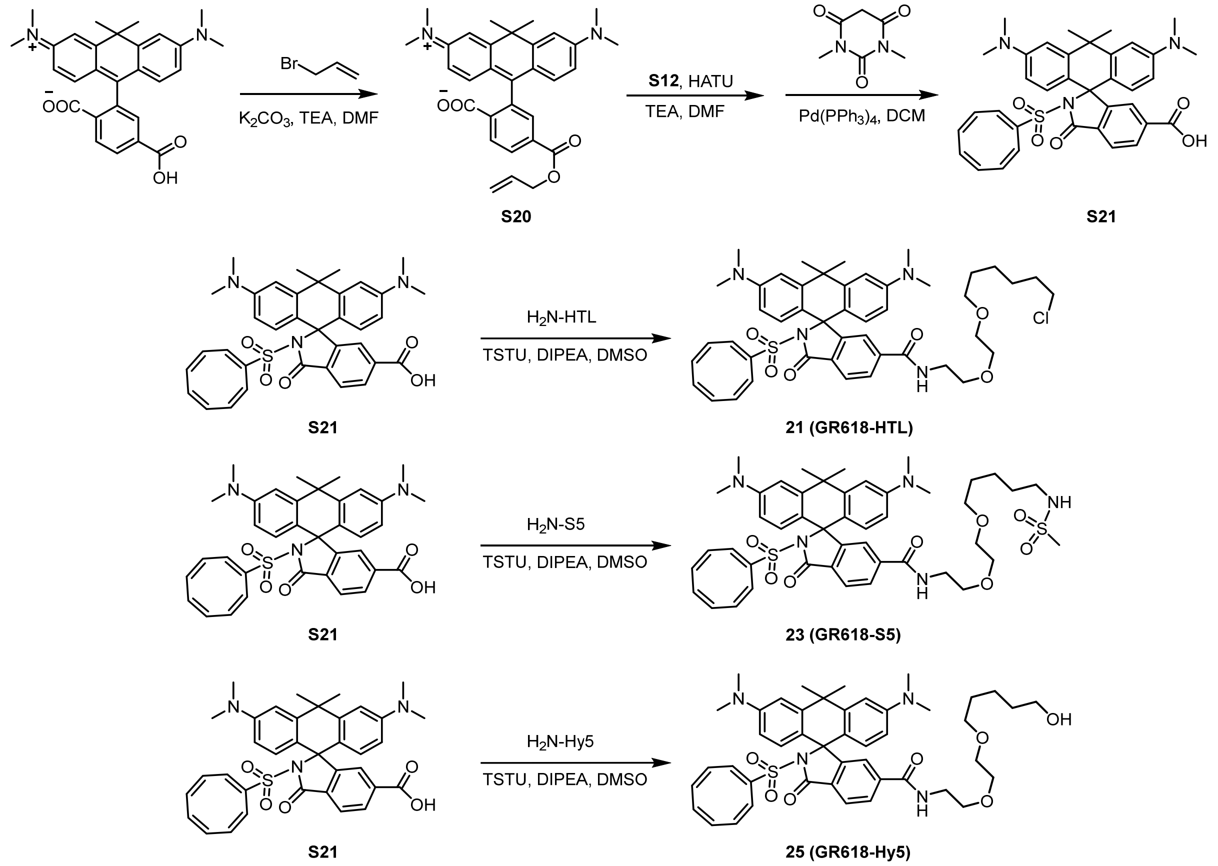


**Supplementary Scheme S6.** Synthetic route of Compound **GR618** (exchangeable) HaloTag Ligands (**HTL, S5, Hy5**).

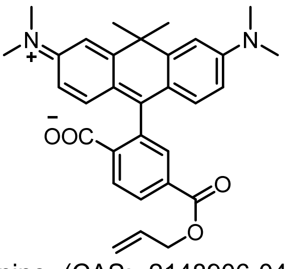

**Compound S20**: 6-carboxycarbopyronine (CAS: 2148906-04-9, commercially acquired from AAT Bioquest; 50 mg, 0.11 mmol, 1.0 equiv.) was dissolved in DMF (4 mL). $K_2CO_3$ (64 mg, 0.22 mmol, 2.0 equiv.) and $Et_3N$ (31 µL, 0.22 mmol, 2.0 equiv.) were added. The reaction was cooled down in an ice bath and allyl bromide (15 µL, 0.16 mmol, 1.5 equiv.) was slowly added. The reaction mixture was allowed to warm up to r.t. and stirred for 2 h. The reaction was monitored by LC/MS. Then, the reaction was concentrated *in vacuo* and purified by flash column purification ($SiO_2$. 0 to 10% MeOH in DCM in 12 CV) afforded the desired compound **S20** (53 mg, 0.10 mmol, 97 %) as a blue solid.

**$^1$H NMR** (400 MHz, Chloroform-d) δ 8.26 (dd, J = 8.0, 1.3 Hz, 1H), 8.07 (dd, J = 8.0, 0.8 Hz, 1H), 7.75 (t, J = 1.0 Hz, 1H), 7.05 – 6.83 (m, 2H), 6.72 – 6.37 (m, 4H), 5.97 (ddt, J = 17.2, 10.4, 5.9 Hz, 1H), 5.42 – 5.22 (m, 2H), 4.77 (dt, J = 5.9, 1.3 Hz, 2H), 3.01 (s, 12H), 1.91 (s, 3H), 1.79 (s, 3H).

**$^{13}$C NMR** (101 MHz, Chloroform-d) δ 169.89, 165.18, 155.52, 150.84, 147.10, 135.89, 131.78, 131.01, 130.29, 129.03, 125.43, 125.01, 119.24, 111.76, 109.35, 77.47, 77.15, 76.84, 66.41, 40.56, 38.74, 35.80, 32.67.

**HRMS** (m/z): calculated for $C_{31}H_{33}N_2O_4$ [M+H] $^+$ 497.2435 found 497.2435.

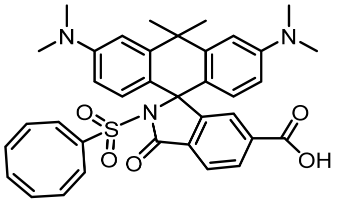

**Compound S21**: A dry Schlenk flask was charged with compound 53 mg **S20** (53 mg, 0.10 mmol) and dissolved in dry DCM (4 mL). Phosphorus oxychloride (150 µL, 1.6 mmol, 15 eq.) was added, the mixture was heated to 50°C and left stirring for 2h. Subsequently, **S12** (78 mg, 0.43 mmol, 4 eq.) in 8 mL dry MeCN and DIPEA (530 µL, 3.2 mmol, 30 eq) was added and the mixture was stirred at 70°C for 10 min. The solvent was evaporated, $H_2O$ (1 mL) was added and the aqueous layer was dried over $MgSO_4$, filtered and concentrated. The crude reaction mixture was purified by flash column purification ($SiO_2$. 20 to 80% EtOAc in n-hexane in 16 CV) to afford the 5.4 mg of the intermediate compound as a blue solid.
**LC/MS** (10% to 90% MeCN/$H_2O$, 6 min, 0.5 mL/min) 3.18 min, 662 m/z, 98%.
It was taken up in MeOH/$CH_2Cl_2$ (5:1, 5 mL) in a dry Schlenk flask together with 1,3-Dimethylbarbituric acid (7.5 mg, 48 µmol, 6 eq.) tetrakis (triphenyl-phosphine) palladium (0) (2.8 mg, 2.4 µmol, 0.3 eq.). The reaction mixture was stirred at rt for 30 min. After the solvent was evaporated, the crude product was dissolved in DMSO (0.5 mL) and purified by preparative HPLC (8 mL/min, 40 % to 99 % MeCN/$H_2O$ + 0.1 % TFA in 55 min) to obtain **S21** (4.2 mg, 6.7 µmol, 84 %) as a blue solid.
**LC/MS** (10% to 90% MeCN/$H_2O$, 6 min, 0.5 mL/min) 2.62 min, 622 m/z, 97%.
**$^1$H NMR** (400 MHz, DMSO-$d_6$) $^1$H NMR (400 MHz, DMSO) δ 8.02 (s, 2H), 7.11 (s, 1H), 6.90 (d, J = 2.7 Hz, 2H), 6.68 – 6.49 (m, 4H), 6.31 (s, 1H), 6.01 – 5.56 (m, 6H), 2.94 (s, 12H), 1.78 (s, 6H).
**$^{13}$C NMR** (101 MHz, DMSO-$d_6$) δ 166.15, 165.95, 155.48, 149.56, 145.18, 141.24, 136.94, 132.42, 129.66, 129.44, 128.47, 124.56, 124.18, 112.33, 110.19, 71.74, 37.63, 36.11, 32.46.
**HRMS** (m/z): calculated for $C_{36}H_{36}N_3O_5S$ $[M+H]^+$ 622.2370 found 622.2368.

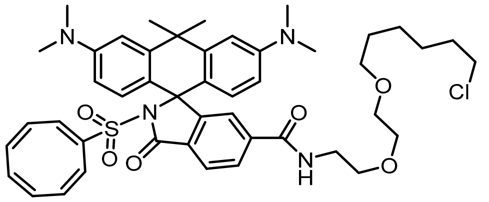

**Compound 21 (GR618-HTL)**: Compound **S21** (2 mg, 3.2 µmol, 1.0 equiv.), TSTU (2 mg, 6.4 µmol, 2 eq.), $H_2N$-HaloTag Ligand[24] (8.9 mg, 4.8 µmol, 1.5 equiv.), and DIPEA (3 µL, 18.5 µmol, 4 eq.) were dissolved in 200 µL DMSO. The mixture was stirred at r.t. for 1 h. Purification by prep-HPLC yielded **GR618-HTL** (~1 mg, 38%) as a blue solid.
**LC/MS** (10% to 90% MeCN/$H_2O$, 6 min, 0.5 mL/min) 3.10 min, 827 m/z, 97%.
**HRMS** (m/z): calculated for $C_{46}H_{56}N_4O_6ClS$ $[M+H]^+$ 827.3604 found 827.3607.

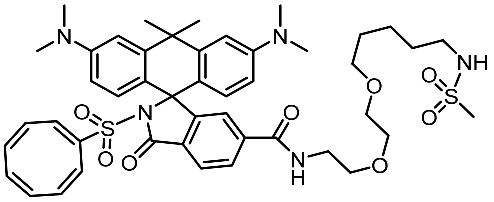

**Compound 23 (GR618-S5)**: $H_2N$-S5 was synthesized as previously reported[7]. Compound **S21** (1 mg, 1.6 µmol, 1.0 equiv.), TSTU (1 mg, 3.2 µmol, 2 eq.), $H_2N$-S5 (3.4 mg, 9.6 µmol, 8 equiv.), and DIPEA (3 µL, 18.5 µmol, 8 eq.) were dissolved in 100 µL DMSO. The mixture was stirred at r.t. for 1 h. Purification by prep-HPLC yielded **GR618-S5** (<1 mg) as a blue solid.
**LC/MS** (10% to 90% MeCN/$H_2O$, 6 min, 0.5 mL/min) 2.64 min, 872 m/z, 99%.

**HRMS** (m/z): calculated for $C_{46}H_{58}N_5O_8S_2^{2+}$ [M+2H] $^{2+}$ 436.1947 found 436.1918.

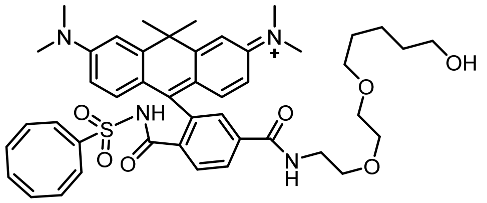

**Compound 25 (GR618-Hy5)**: $H_2N$-Hy5 was synthesized as previously reported[7]. Compound **S21** (1 mg, 1.6 µmol, 1.0 equiv.), TSTU (1 mg, 3.2 µmol, 2 eq.), $H_2N$-Hy5 (2.0 mg, 9.6 µmol, 8 equiv.), and DIPEA (3 µL, 18.5 µmol, 8 eq.) were dissolved in 100 µL DMSO. The mixture was stirred at r.t. for 1 h. Purification by prep-HPLC yielded **GR618-Hy5** (<1 mg) as a blue solid.
**LC/MS** (10% to 90% MeCN/$H_2O$, 6 min, 0.5 mL/min) 2.28 min, 722 m/z, 98%.
**HRMS** (m/z): calculated for $C_{45}H_{55}N_4O_7S^+$ [M+H] $^+$ 795.3786 found 795.3784.

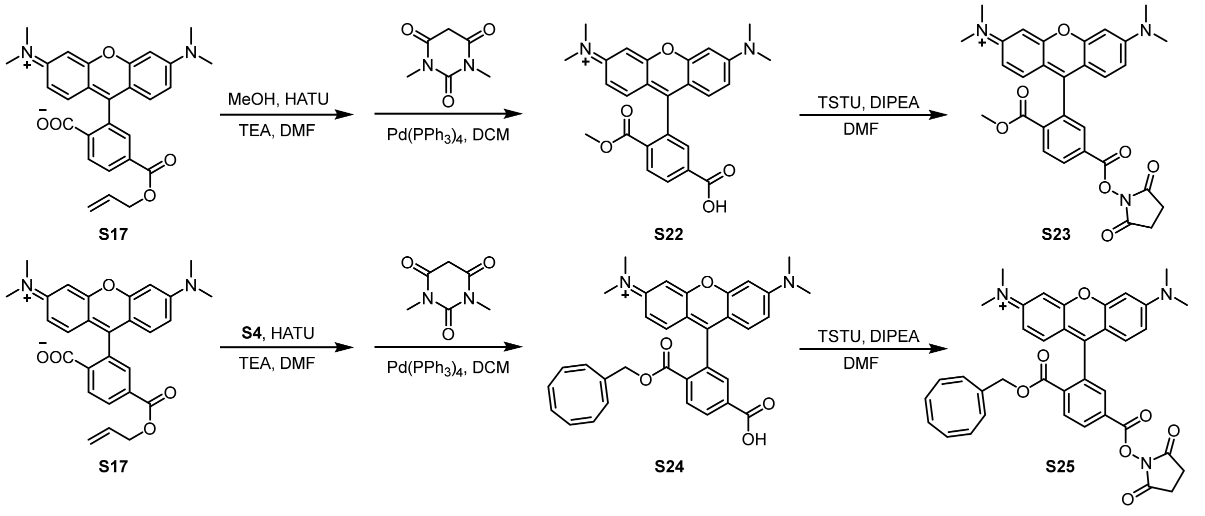


**Supplementary Scheme S7.** Synthetic route of Compound **S23** and **S25** for WGA-conjugation.

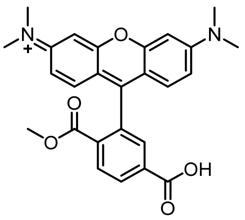

**Compound S22**: Compound **S17** (5 mg, 0.01 mmol, 1.0 equiv.), HATU (7.6 mg, 0.02 mmol, 2.0 equiv.), and DIPEA (10 µL, 0.06 mmol, 6.0 equiv.) were dissolved in dry DMF (0.5 mL) and stirred at r.t. for 5 min. Then, MeOH (2 µL, 0.05 mmol, 5.0 equiv.) was added and stirred at r.t. for 2 h. The reaction mixture was diluted with water and extracted with DCM. The combined organics were washed with brine, dried with $Na_2SO_4$, filtered, and concentrated *in vacuo* to obtain the residue. The products were mixed with 1,3-dimethylbarbituric acid (3.1 mg, 0.02 mmol, 5.0 equiv.), and tetrakis (triphenyl-phosphine) palladium (0) (9.5 mg, 0.008 mmol, 2.0 equiv.) in DCM (2 mL) and stirred at r.t. for 30 min. The mixture was concentrated under reduce pressure and purified by prep-HPLC to obtain compound **S22** (2.5 mg, 56%) as a purple solid.
**LC/MS** (10% to 90% MeCN/$H_2O$, 6 min, 0.4 mL/min) 2.83 min, 445.3 m/z, >99%.

**HRMS** (m/z): calculated for $C_{26}H_{25}N_2O_5^+$ [M] $^+$ 445.1758 found 445.2363.
**$^1$H NMR** (400 MHz, Methanol-d4) δ 8.41 (d, *J* = 1.0 Hz, 2H), 8.01 (t, *J* = 1.0 Hz, 1H), 7.13 (d, *J* = 9.5 Hz, 2H), 7.07 (dd, *J* = 9.5, 2.4 Hz, 2H), 7.01 (d, *J* = 2.4 Hz, 2H), 3.63 (s, 3H), 3.32 (s, 12H).

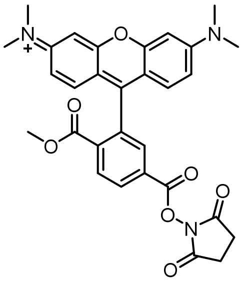

**Compound S23**: Compound **S22** (2.5 mg, 0.006 mmol, 1.0 equiv.), TSTU (3.3 mg, 0.01 mmol, 2.0 equiv.), and DIPEA (10 µL, 0.06 mmol, 10.0 equiv.) were dissolved in dry DMF (0.5 mL) and stirred at r.t. for 4 h. The mixture was concentrated *in vacuo* and purified by flash column (DCM: MeOH=10:1 +0.1%TFA) to obtain compound **S23** (2 mg, 61.5%) as a purple solid. Next, the product was used to label wheat germ agglutinin (WGA).
**HRMS** (m/z): calculated for $C_{30}H_{28}N_3O_7^+$ [M] $^+$ 542.1922 found 542.1951

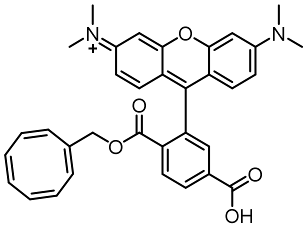

**Compound S24**: Compound **S17** (5 mg, 0.01 mmol, 1.0 equiv.), HATU (7.6 mg, 0.02 mmol, 2.0 equiv.), and DIPEA (10 µL, 0.06 mmol, 6.0 equiv.) were dissolved in dry DMF (0.5 mL) and stirred at r.t. for 5 min. Then compound **S4** (7 mg, 0.05 mmol, 5.0 equiv.) was added and stirred at r.t. for 2 h. Then, the reaction was diluted with water and extracted with DCM. The combined organics were washed with brine, dried with $Na_2SO_4$, filtered, and concentrated *in vacuo* to obtain the residue. The products were mixed with 1,3-dimethylbarbituric acid (3.1 mg, 0.02 mmol, 5.0 equiv.), and tetrakis (triphenyl-phosphine) palladium (0) (6 mg, 0.005 mmol, 1.3 equiv.) in DCM (2 mL) and stirred at r.t. for 30 min. The mixture was concentrated under reduce pressure and purified by prep-HPLC to obtain compound **S24** (2.0 mg, 36%) as a purple solid.
**LC/MS** (10% to 90% MeCN/$H_2O$, 6 min, 0.4 mL/min) 3.40 min, 547.5 m/z, >99%.
**HRMS** (m/z): calculated for $C_{26}H_{25}N_2O_5^+$ [M] $^+$ 547.2227 found 547.2275.
**$^1$H NMR** (400 MHz, Methanol-d4) δ 8.53 (d, *J* = 1.4 Hz, 1H), 8.43 (dd, *J* = 7.9, 1.6 Hz, 1H), 7.58 (d, *J* = 7.9 Hz, 1H), 7.08 (d, *J* = 9.3 Hz, 2H), 7.01 (dd, *J* = 9.4, 2.1 Hz, 2H), 6.94 (d, *J* = 2.2 Hz, 2H), 6.72 (s, 1H), 6.10 – 5.60 (m, 6H), 3.28 (s, 12H).

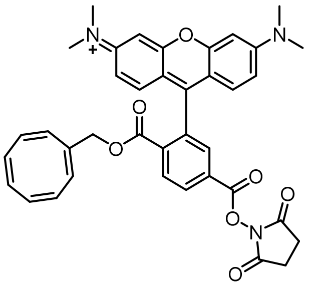

**Compound S25**: Compound **S23** (2.0 mg, 0.003 mmol, 1.0 equiv.), TSTU (2.0 mg, 0.007 mmol, 2.0 equiv.), and DIPEA (5 µL, 0.03 mmol, 10.0 equiv.) were dissolved in dry DMF (0.5 mL) and stirred at r.t. for 4 h. The mixture was concentrated *in vacuo* and purified by flash column (DCM: MeOH=10:1 +0.1%TFA) to obtain compound **S25** (1.1 mg, 56.9%) as a purple solid. Next, the product was used to label wheat germ agglutinin (WGA).
**HRMS** (m/z): calculated for $C_{38}H_{34}N_3O_7^+$ [M] $^+$ 644.2391 found 644.2381.

### Additional data

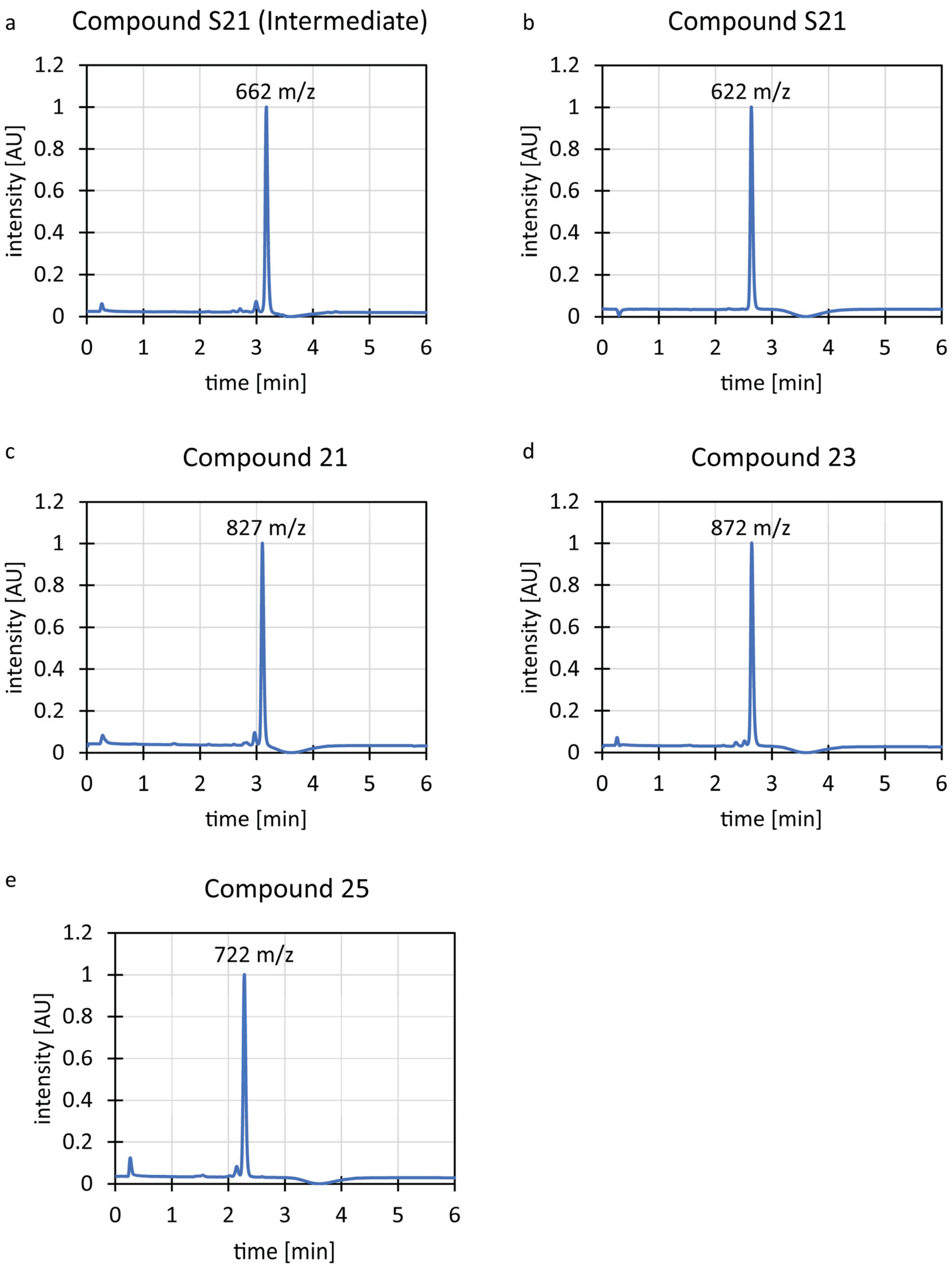


**LC/MS traces of GR618-(x)HTLs.** Normalized absorption intensity at 280 nm and main low-resolution mass spectrometry signal corresponding to the main peak. Signal integration was used to verify that compounds used for further experiments are ≥95% pure.

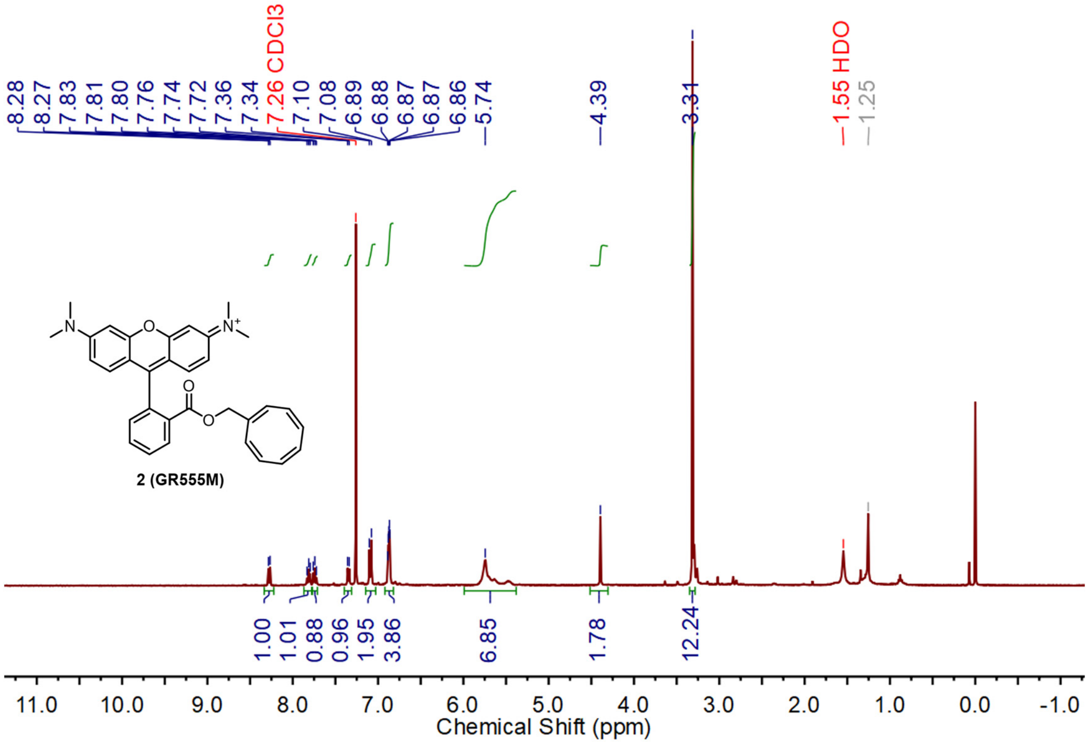


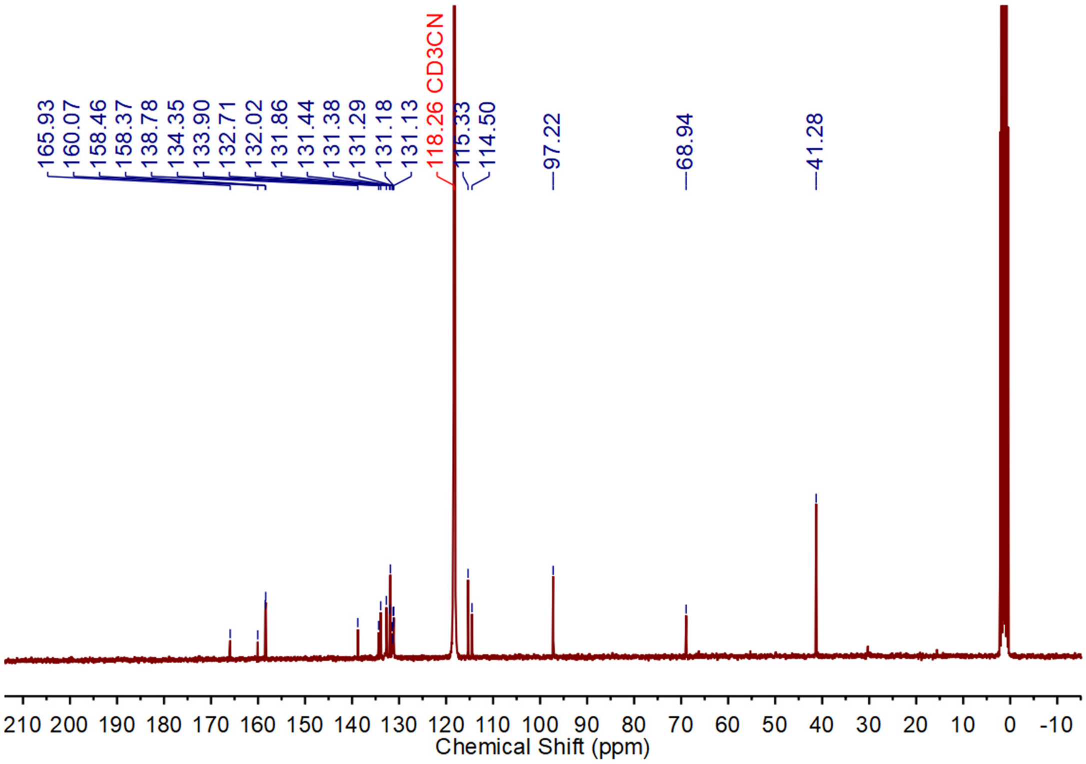


$^{1}H$ NMR and $^{13}C$ NMR spectra of compound **2** (GR555M)

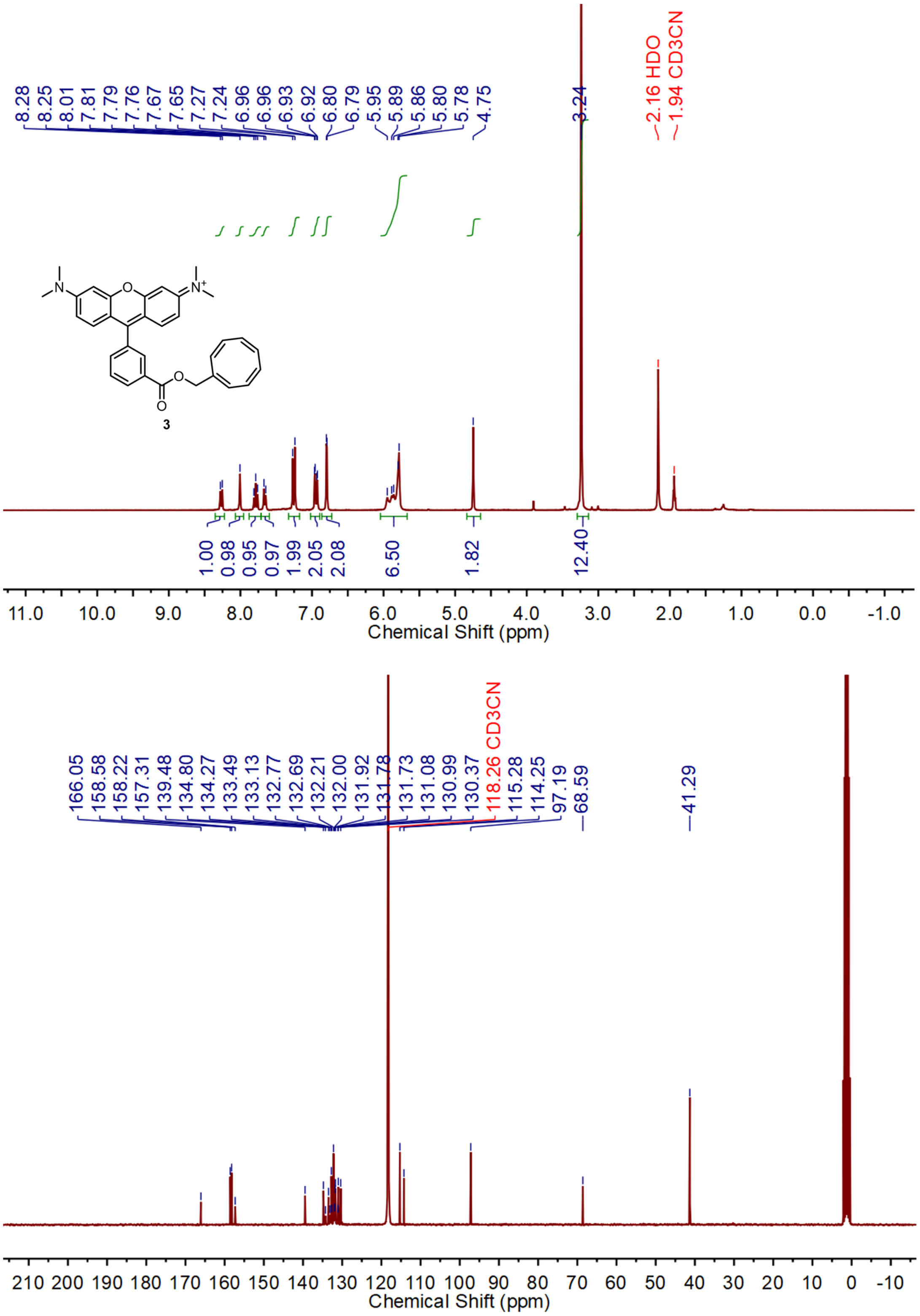


[1]H NMR and [13]C NMR spectra of compound **3**

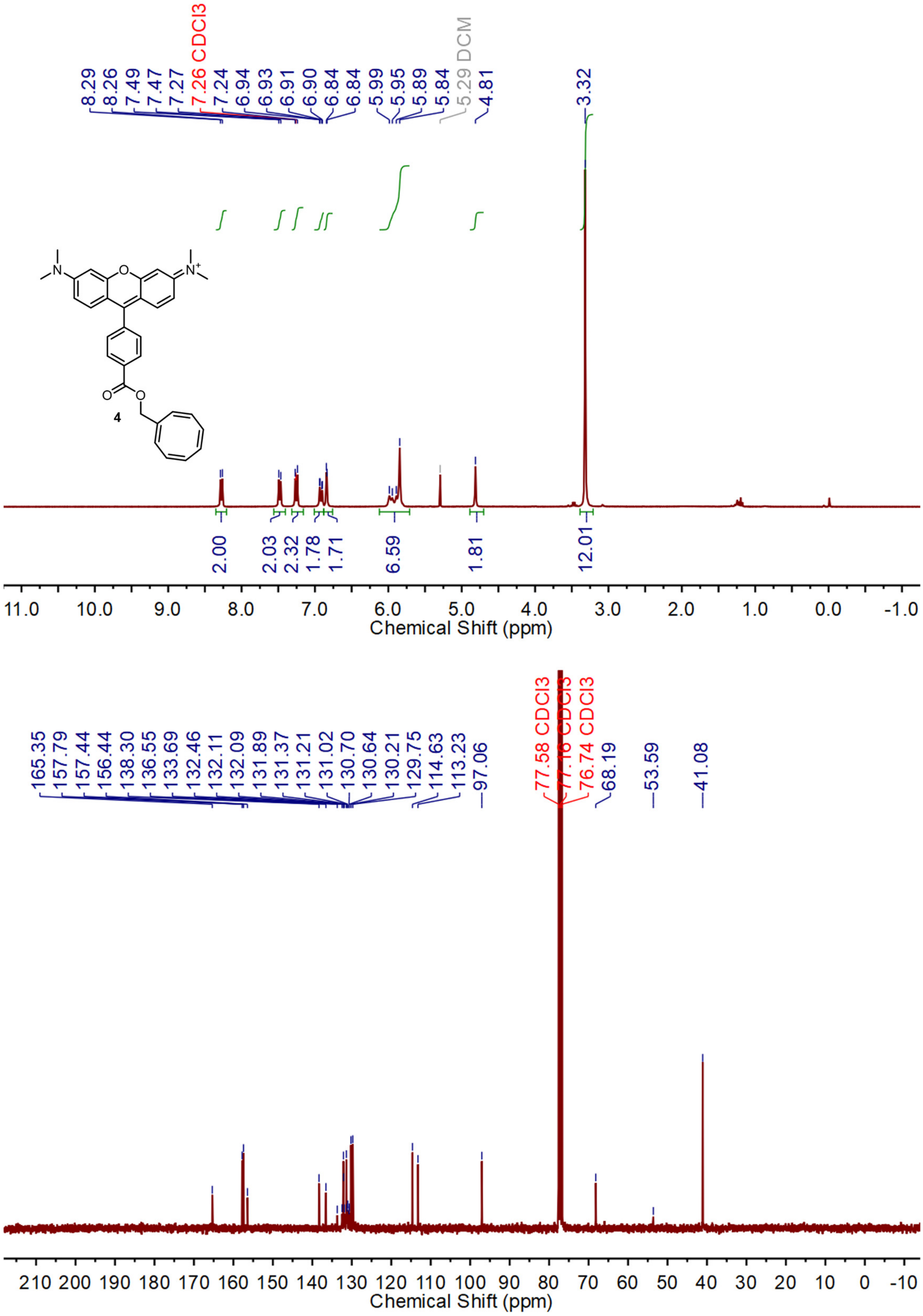


$^1$H NMR and $^{13}$C NMR spectra of compound **4**

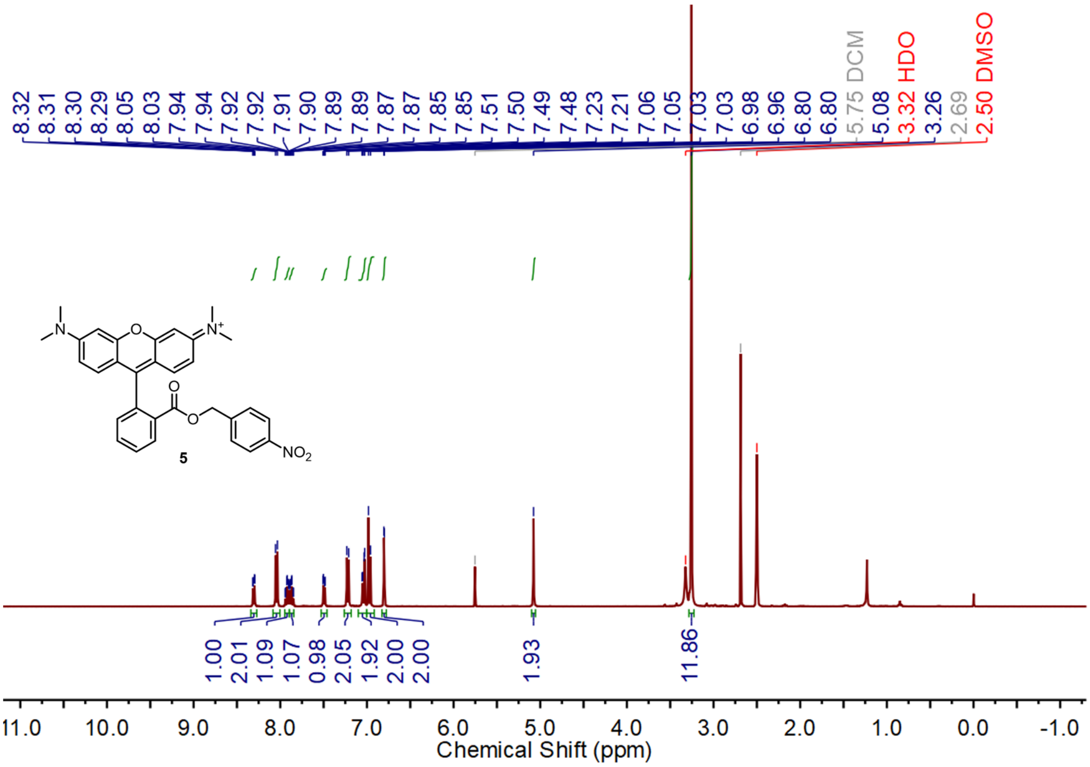


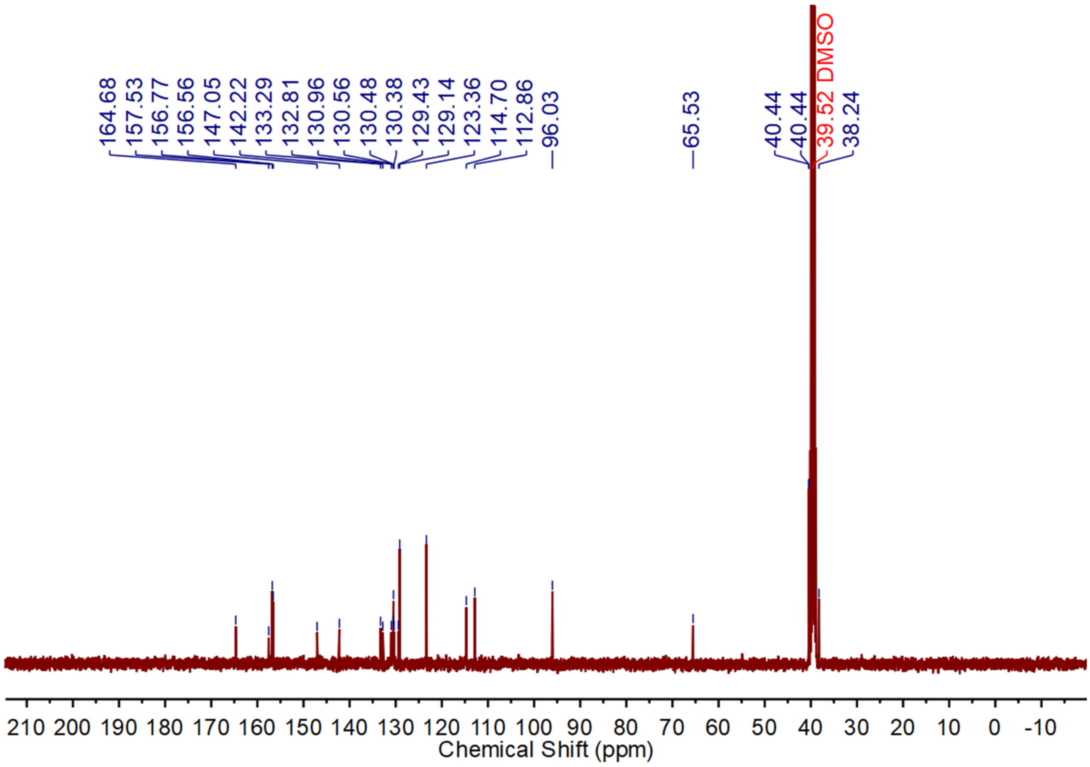


$^{1}$H NMR and $^{13}$C NMR spectra of compound **5**

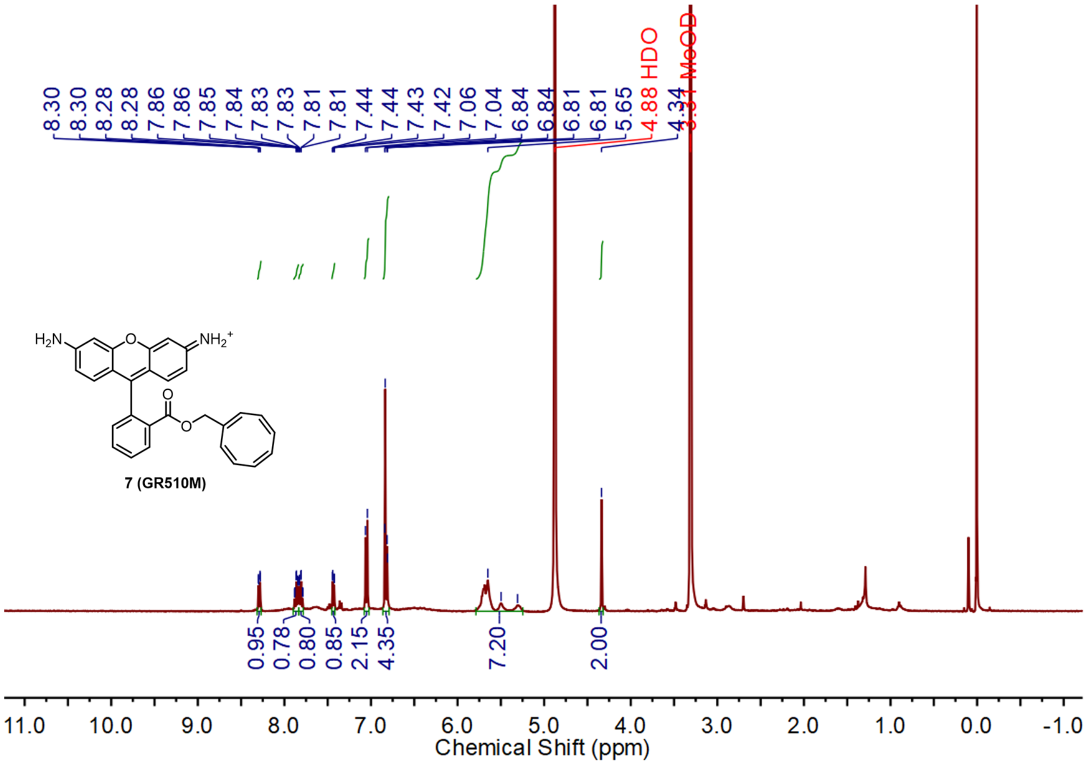


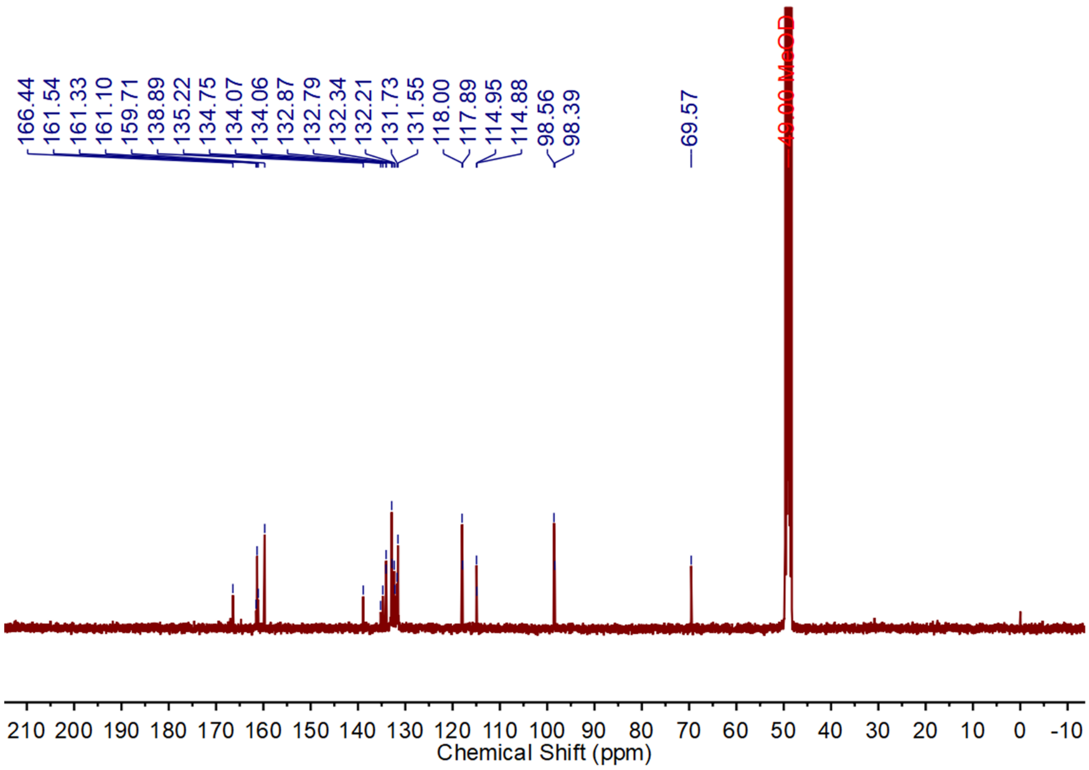


$^{1}$H NMR and $^{13}$C NMR spectra of compound **7** (GR510M)

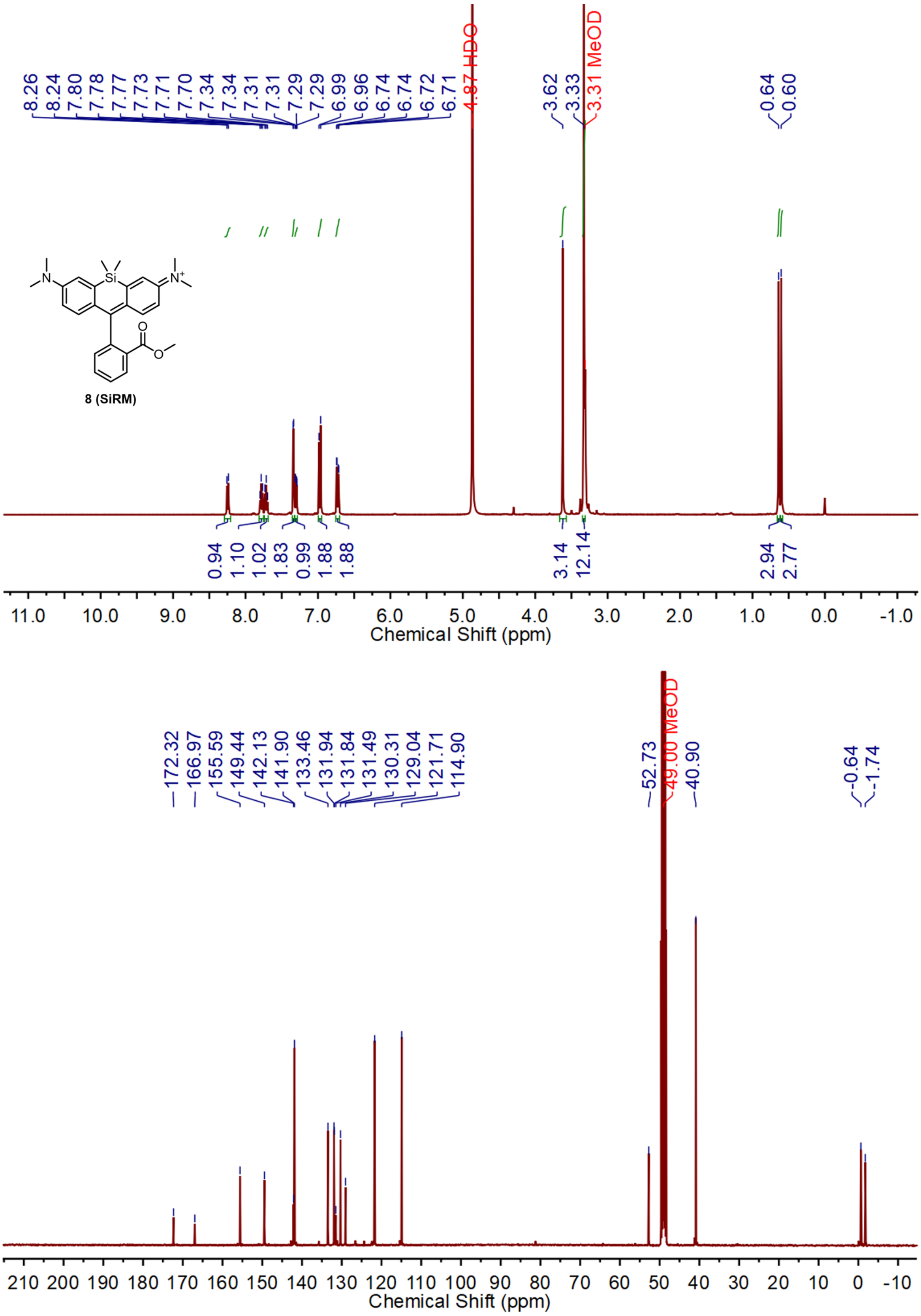


$^{1}H$ NMR and $^{13}C$ NMR spectra of compound **8** (SiRM)

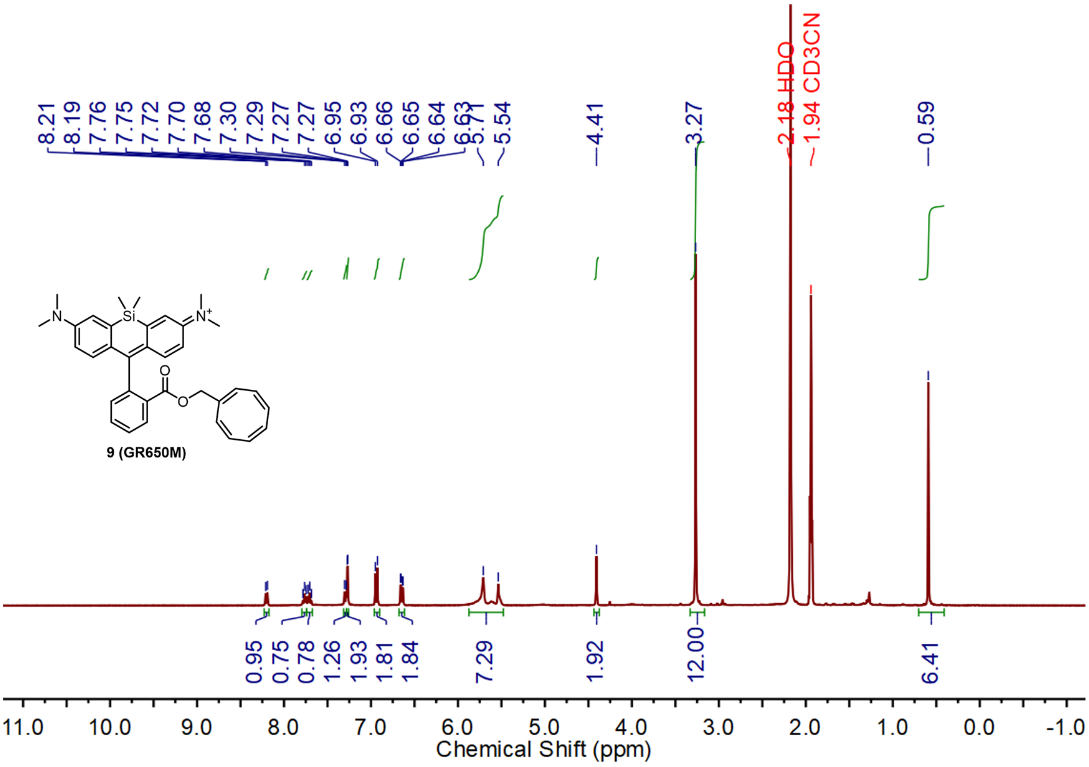

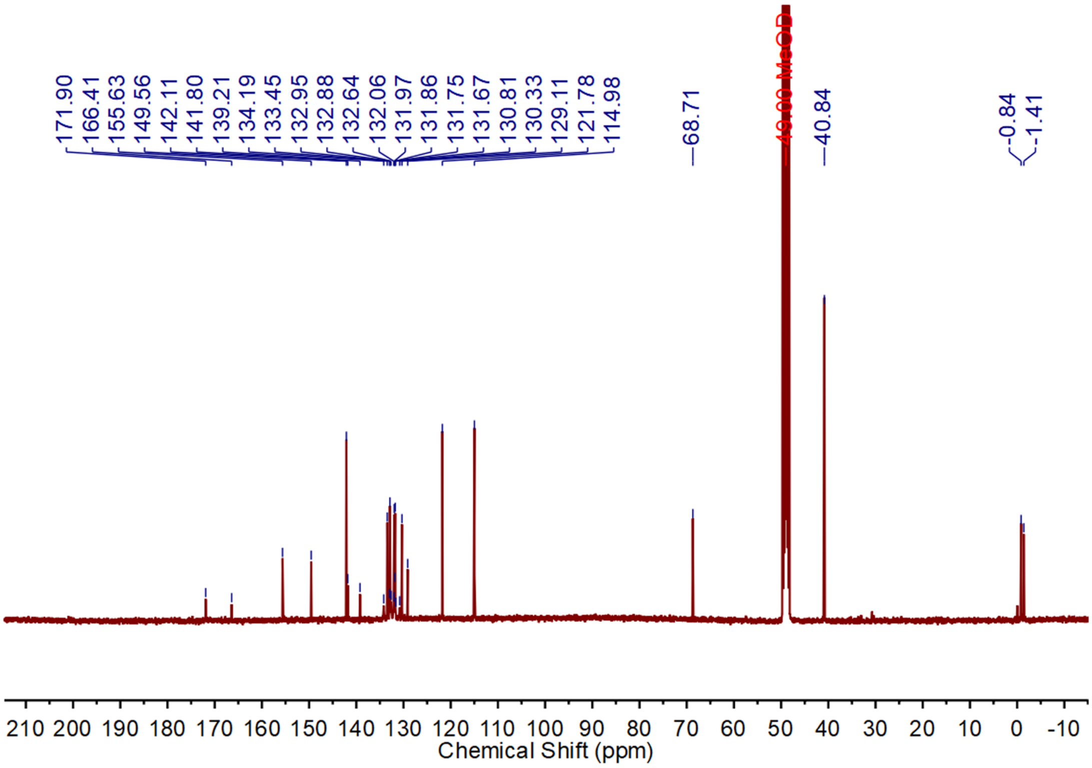

$^{1}H$ NMR and $^{13}C$ NMR spectra of compound **9** (GR650M)

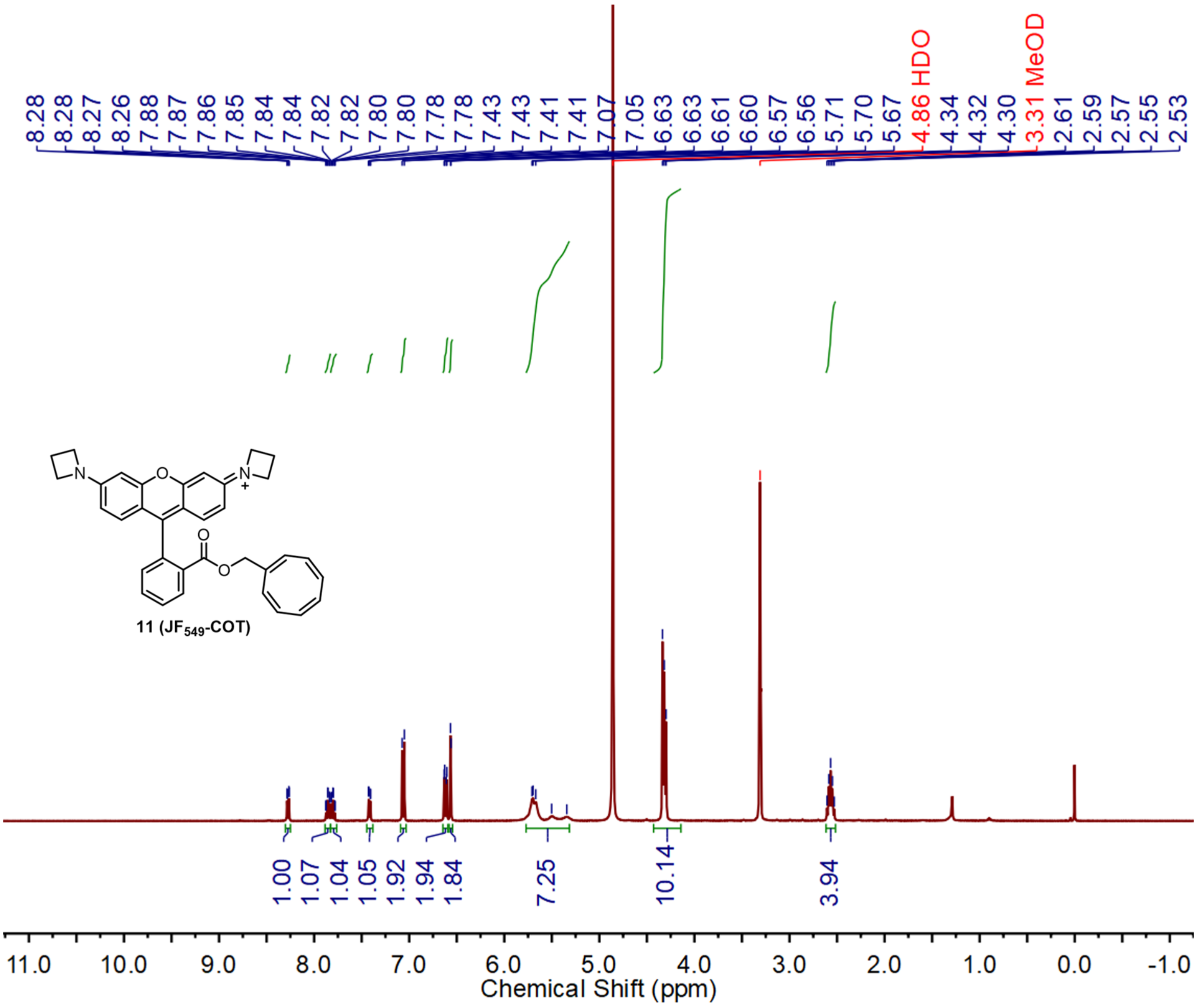


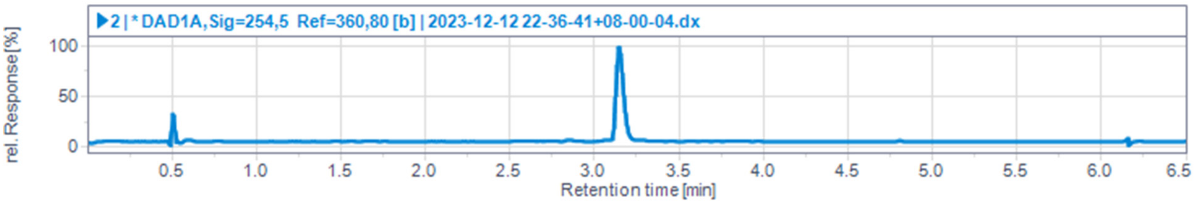


$^{1}$H NMR spectrum and LC/MS trace of compound **11** ($JF_{549}$-COT)

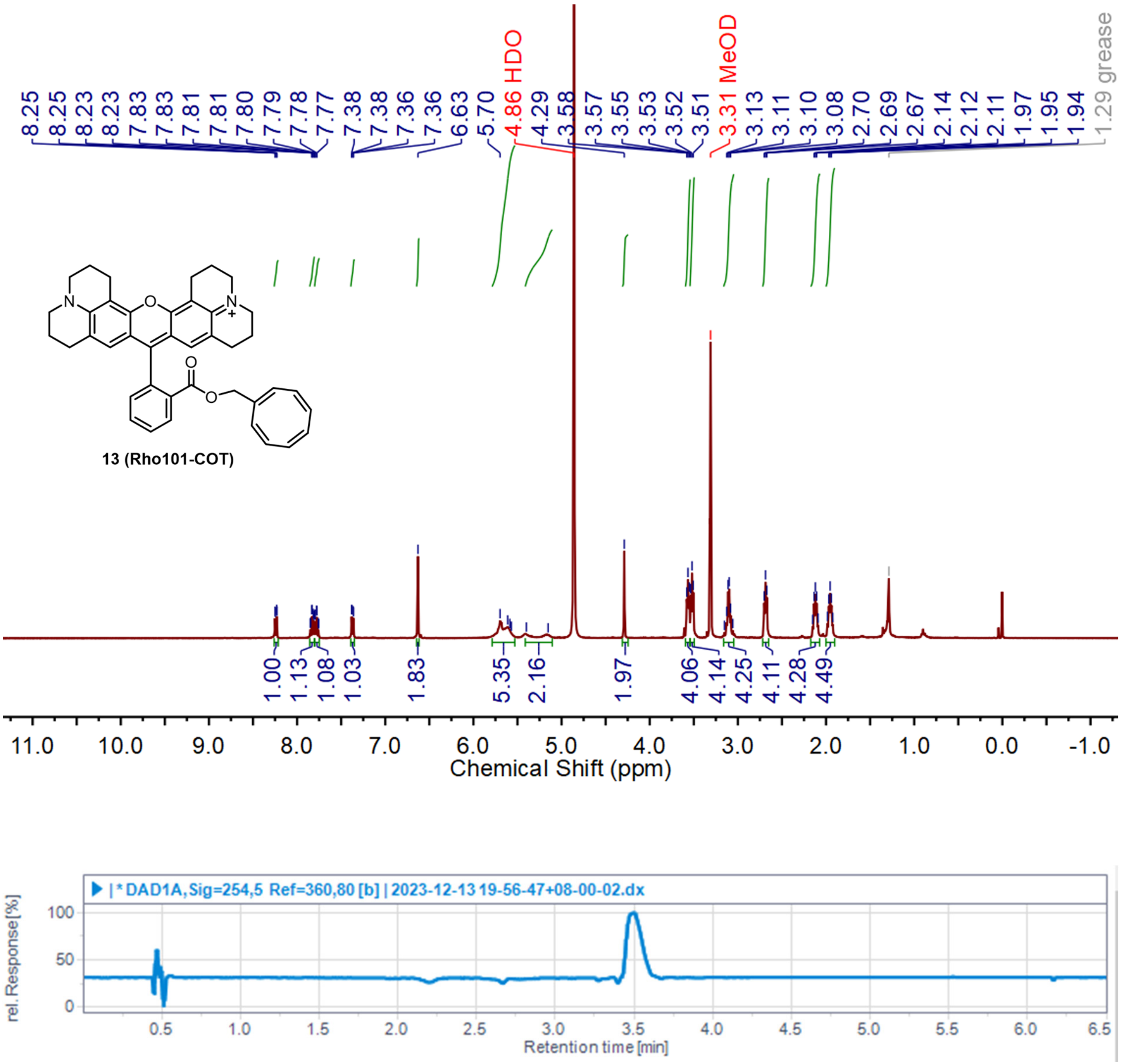


$^{1}H$ NMR spectrum and LC/MS trace of compound **13** (Rho101-COT)

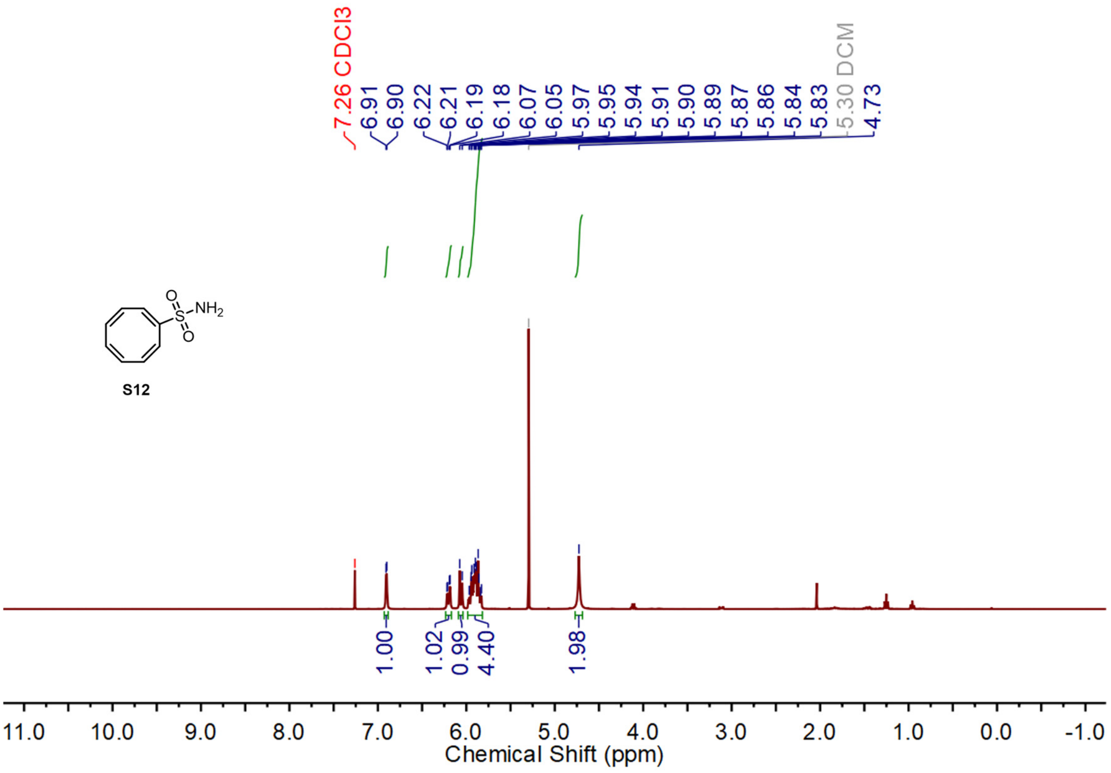



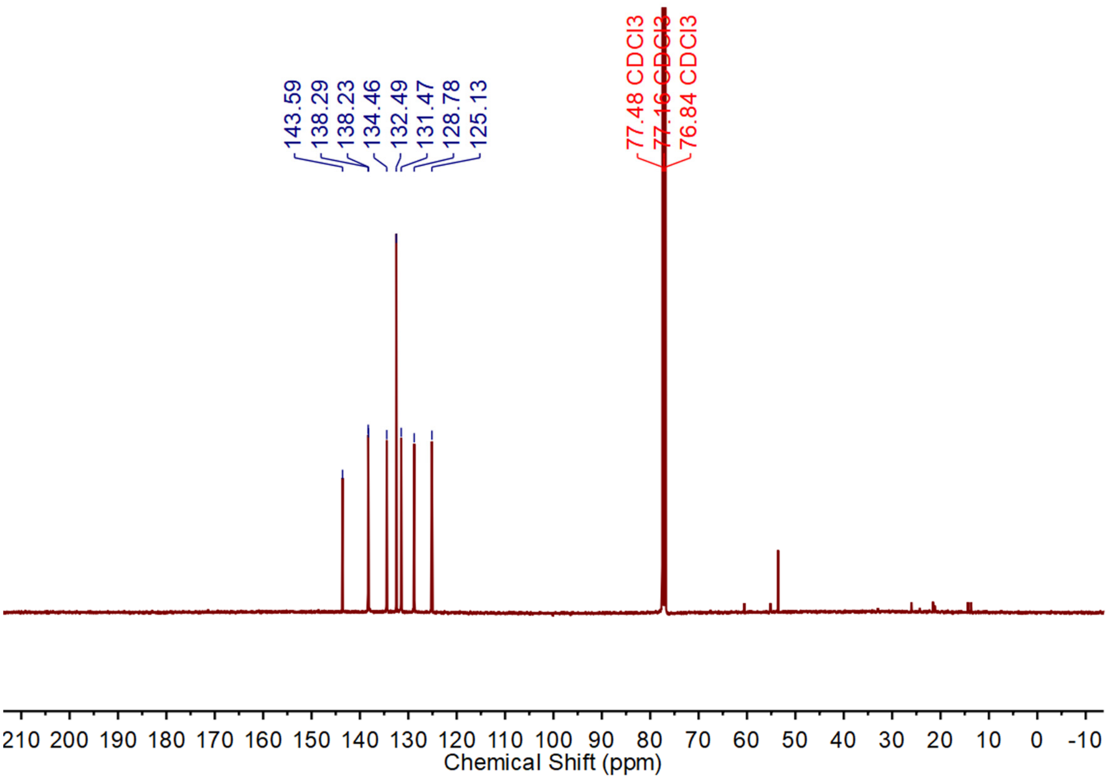


$^{1}H$ NMR and $^{13}C$ NMR spectra of compound **S12**

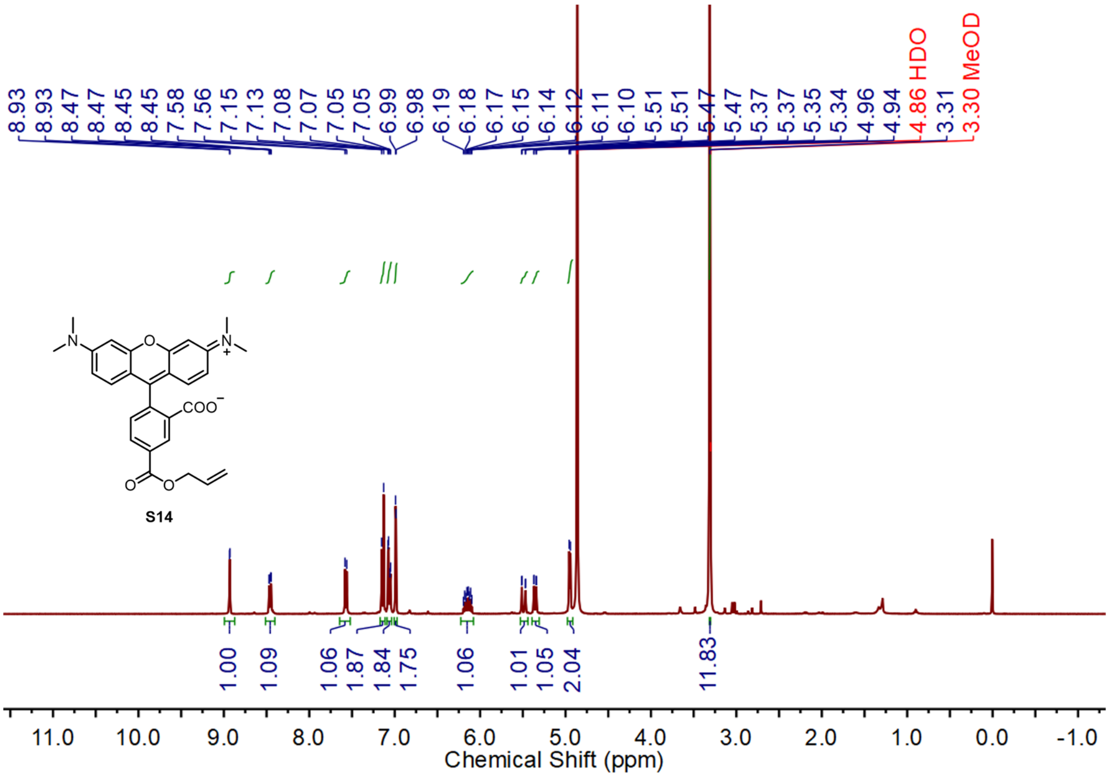


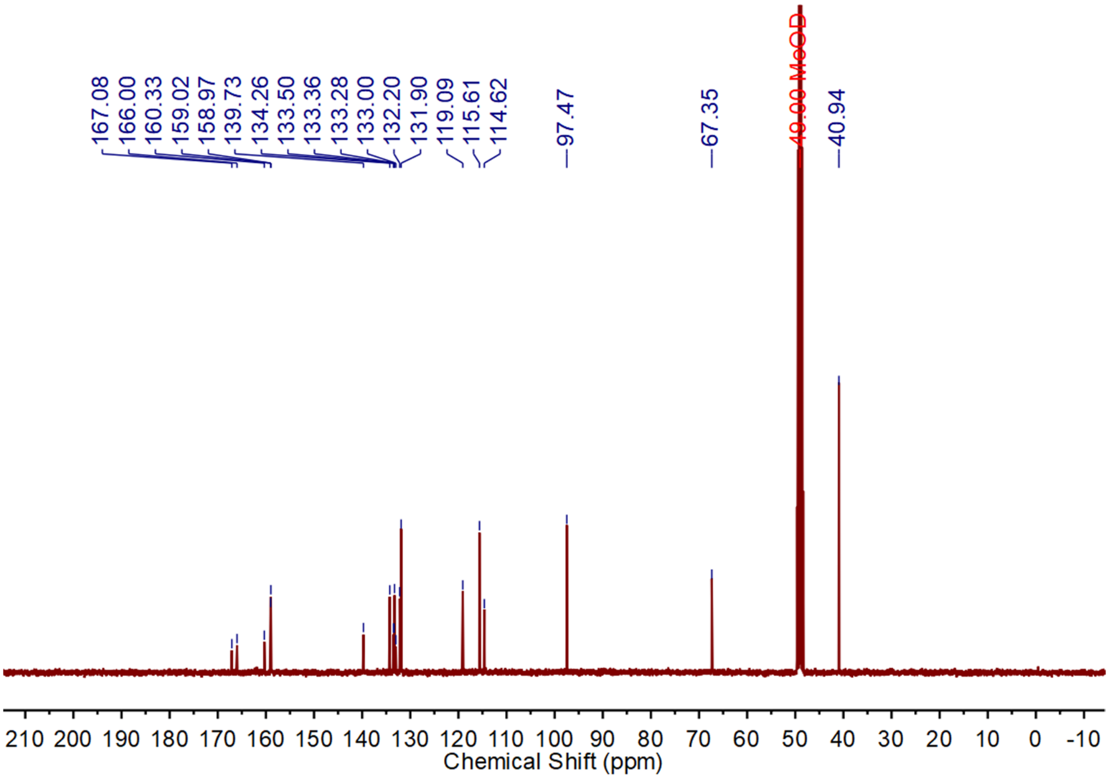


$^1$H NMR and $^{13}$C NMR spectra of compound **S14**

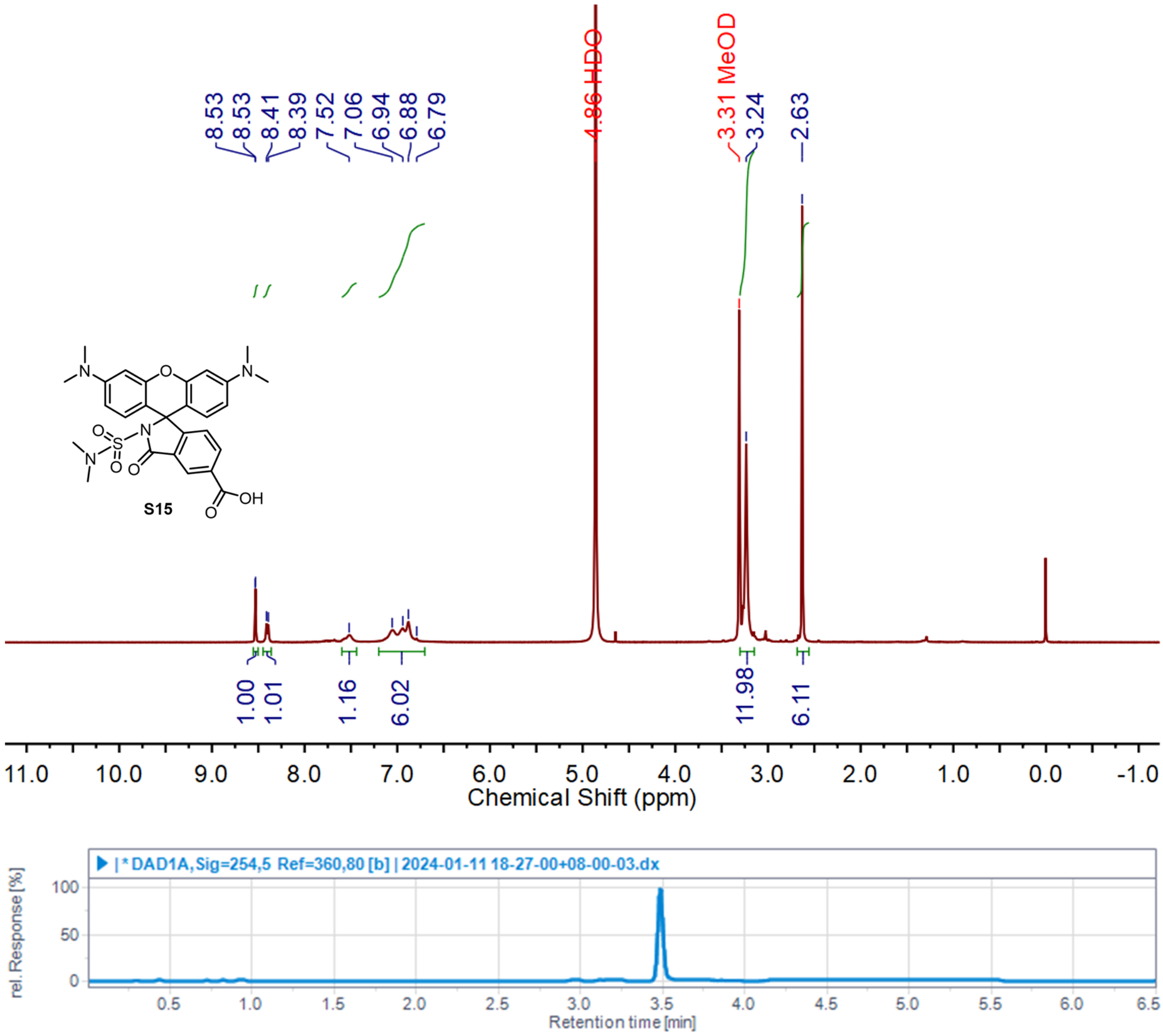


$^{1}$H NMR spectrum and LC/MS trace of compound **S15**

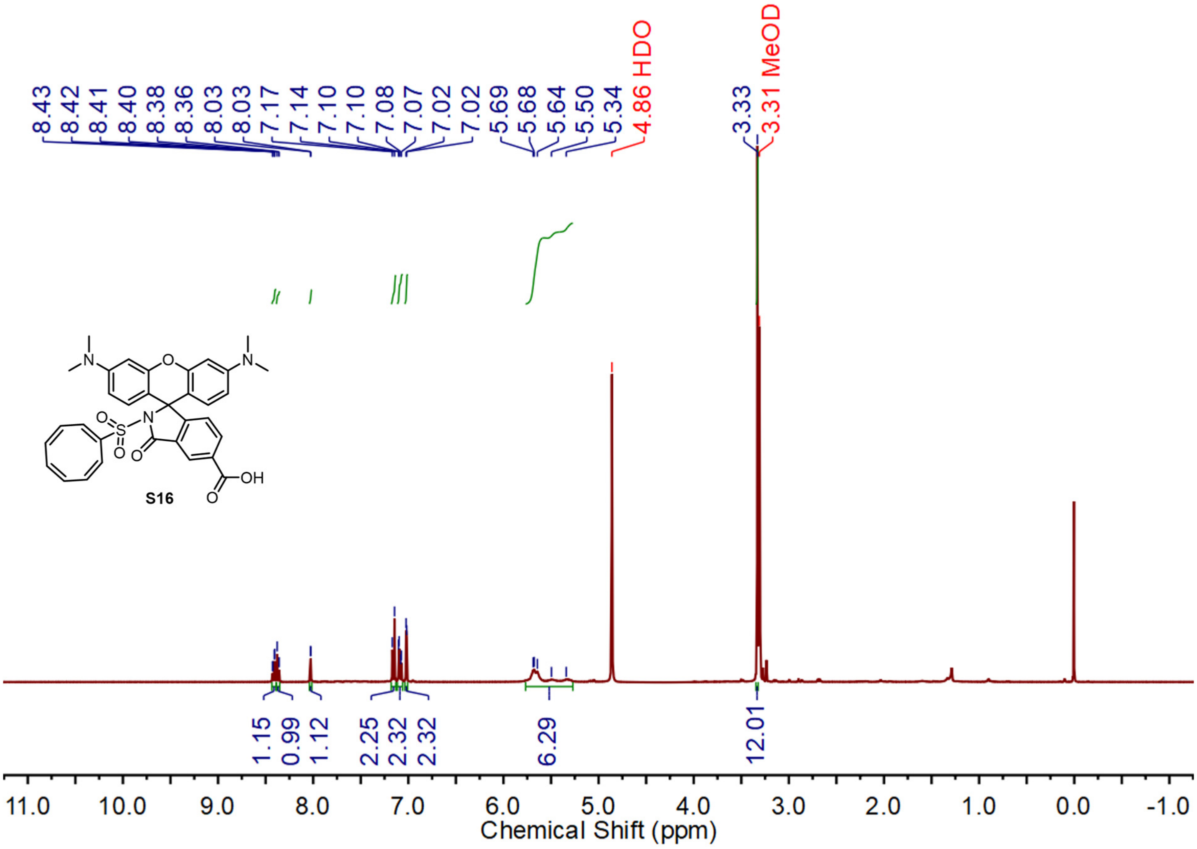


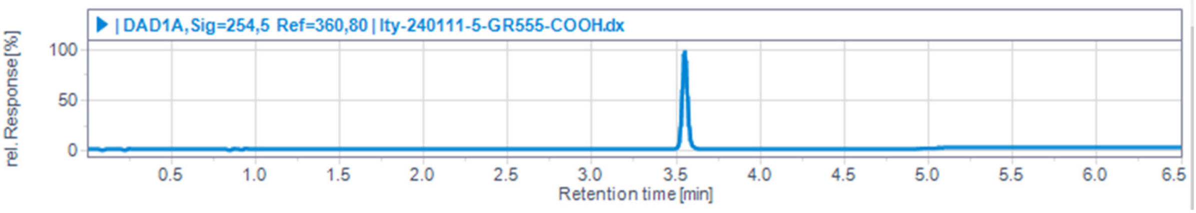


$^{1}$H NMR spectrum and LC/MS trace of compound **S16**

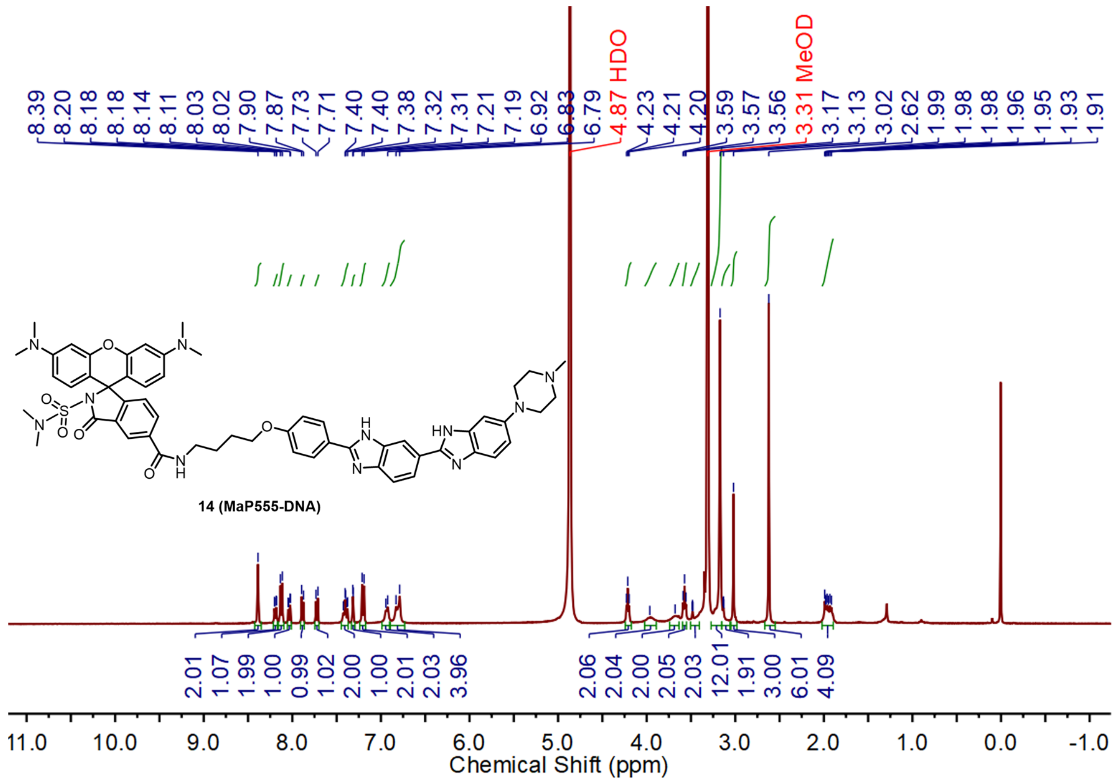

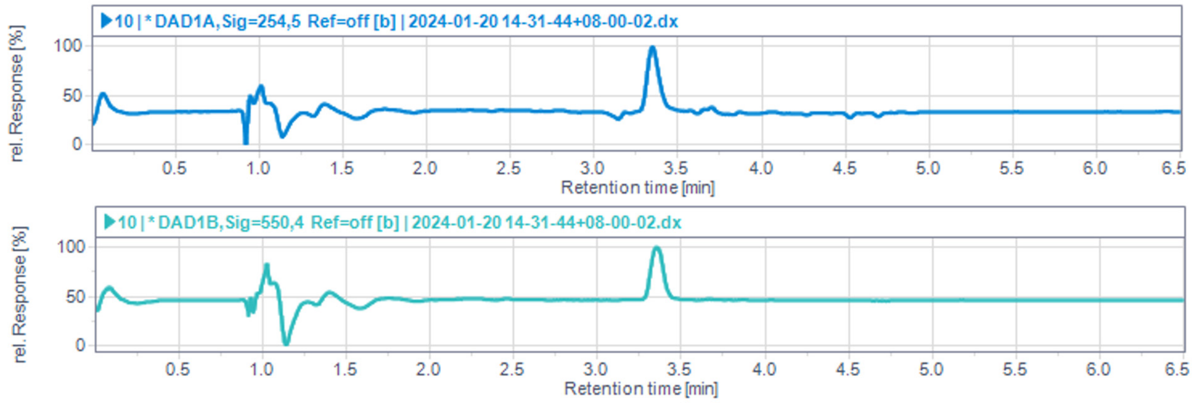


$^{1}$H NMR spectrum and LC/MS trace of compound **14** (MaP555-DNA)

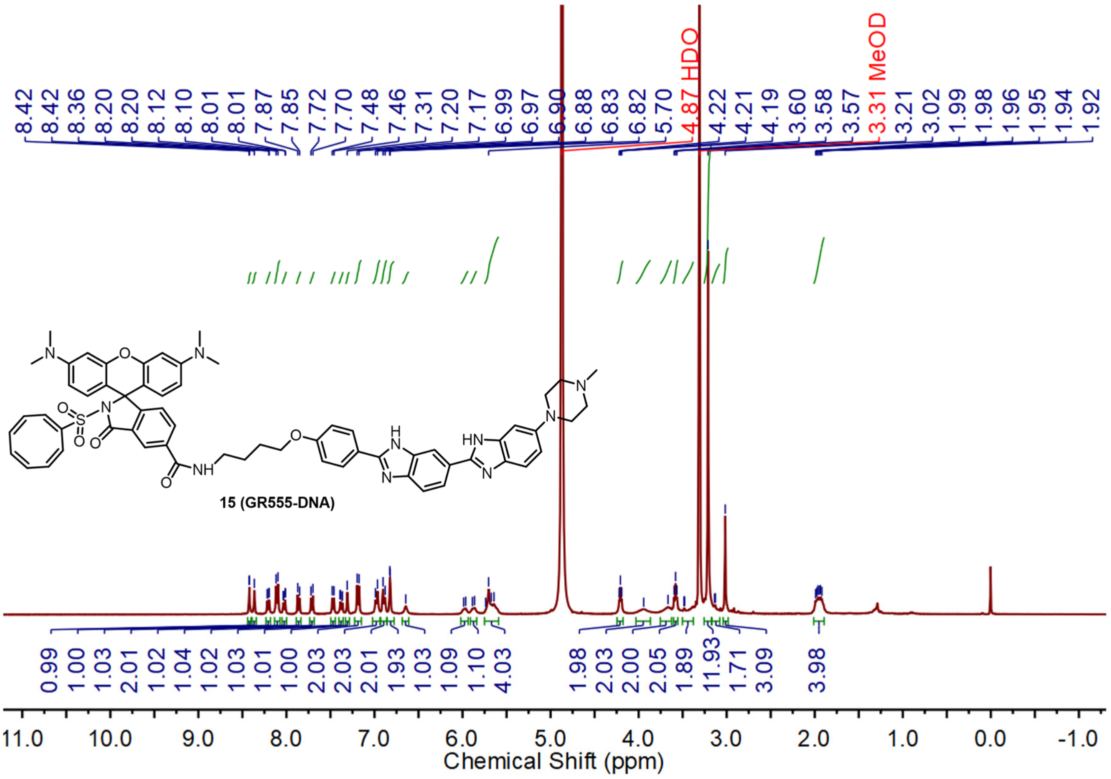


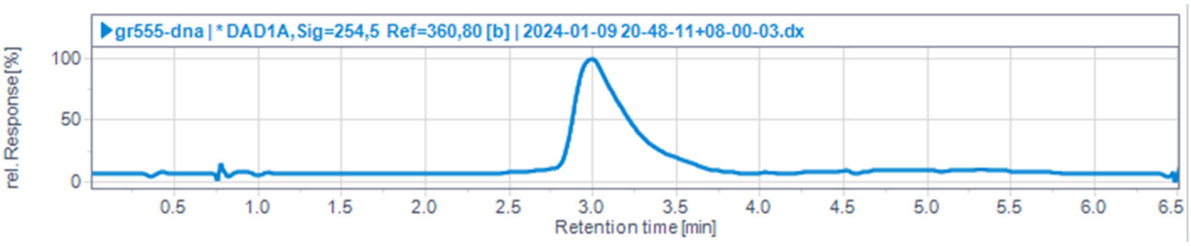


[1]H NMR spectrum and LC/MS trace of compound **15** (GR555-DNA)

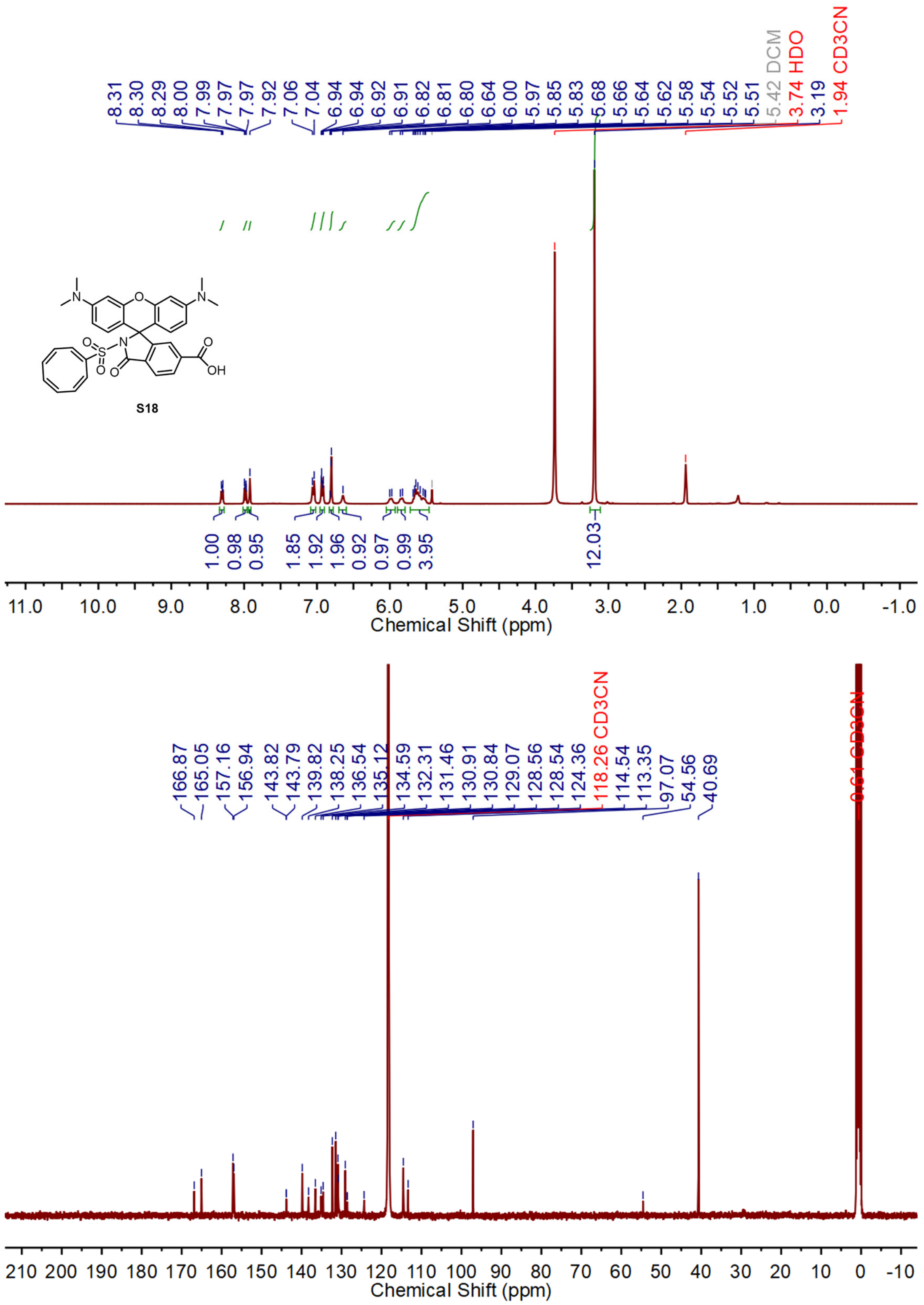


[1]H NMR and [13]C NMR spectra of compound **S18**

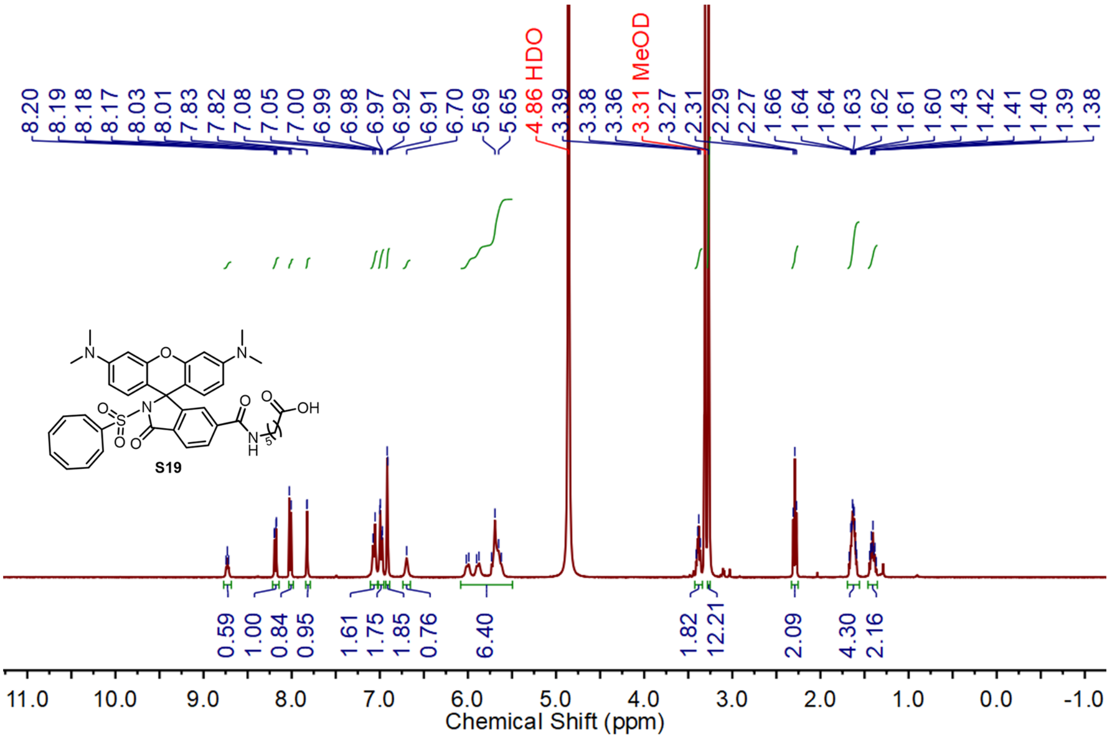


$^1$H NMR spectrum of compound **S19**

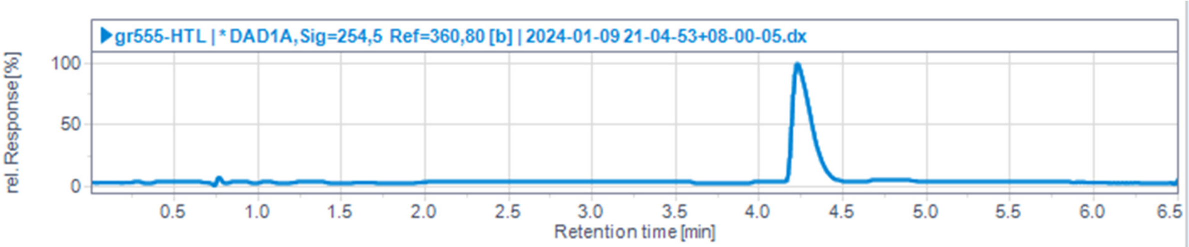


LC/MS trace of compound **19** (GR555-HTL)

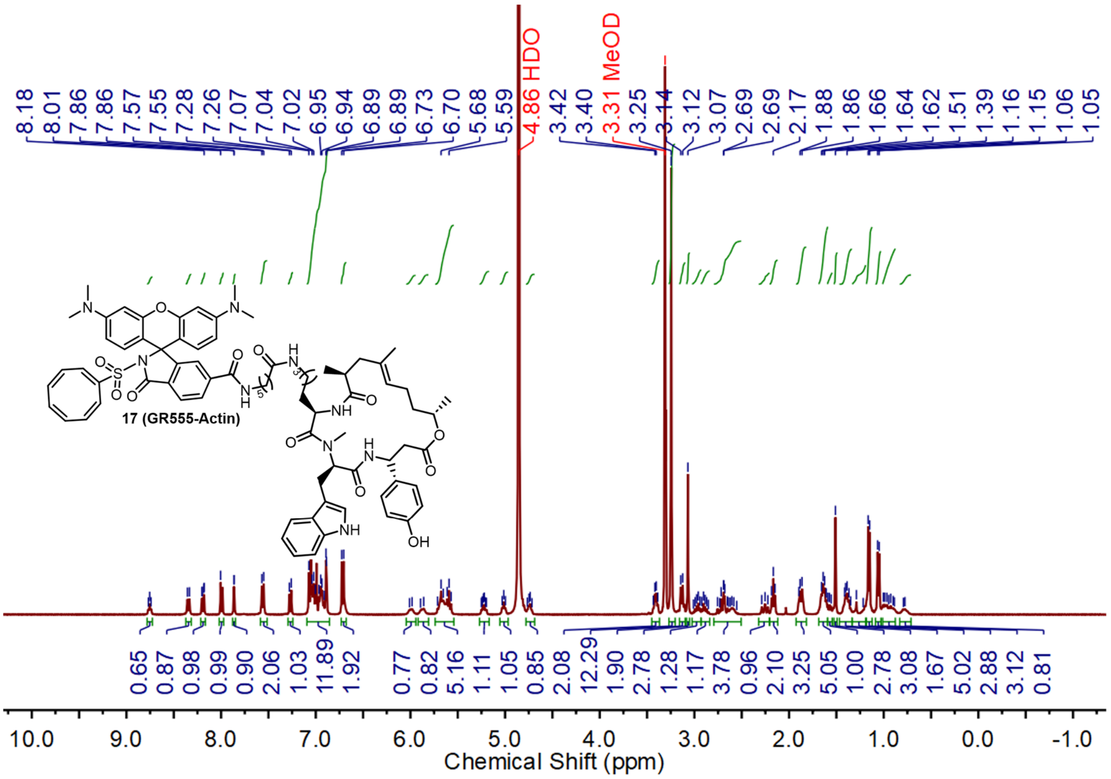

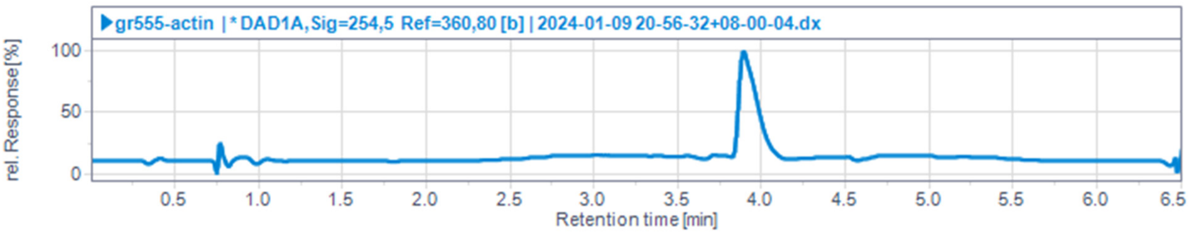

[1]H NMR spectrum and LC/MS trace of compound **17** (GR555-Actin)

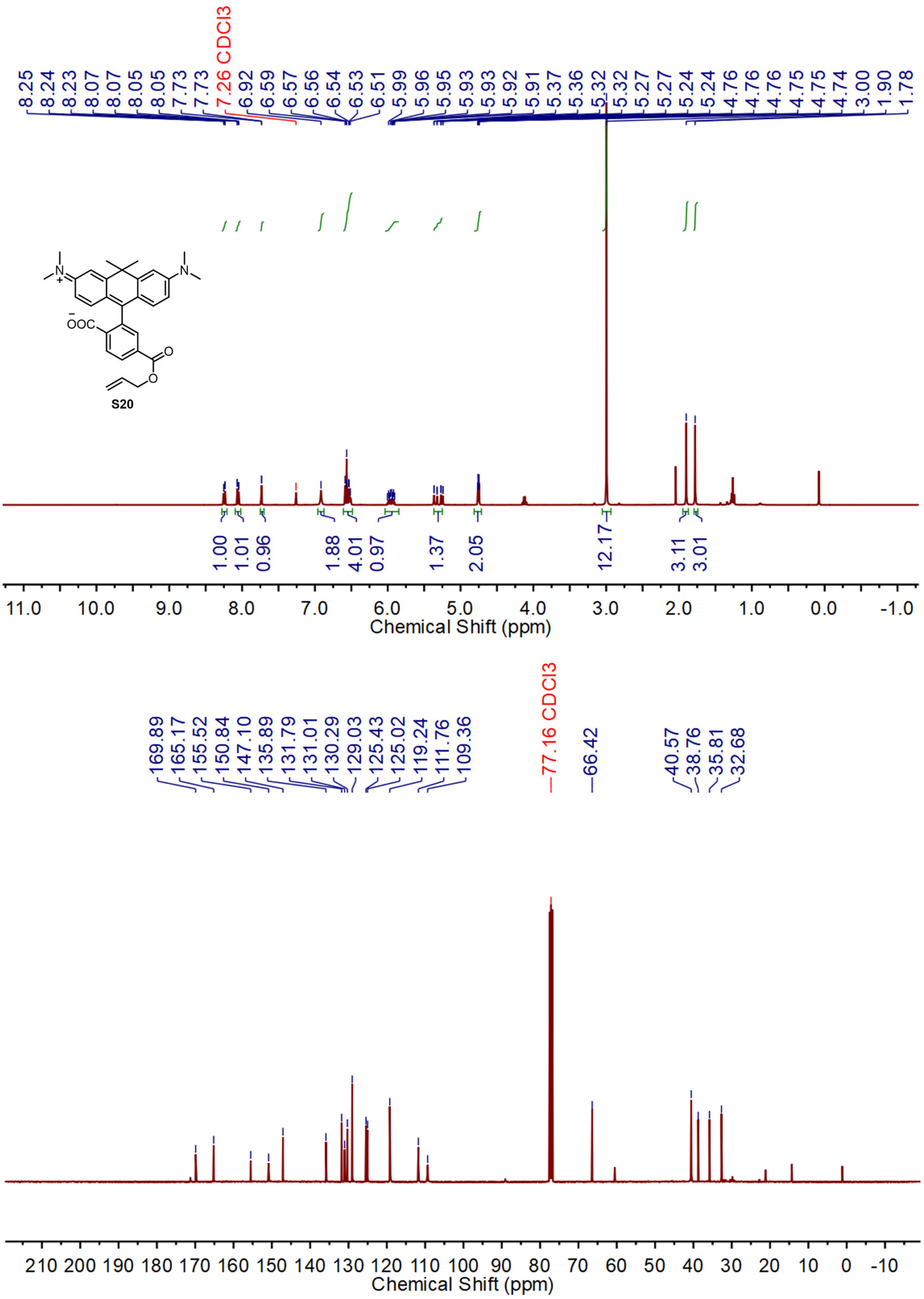


[1]H NMR and [13]C NMR spectra of compound **S20**

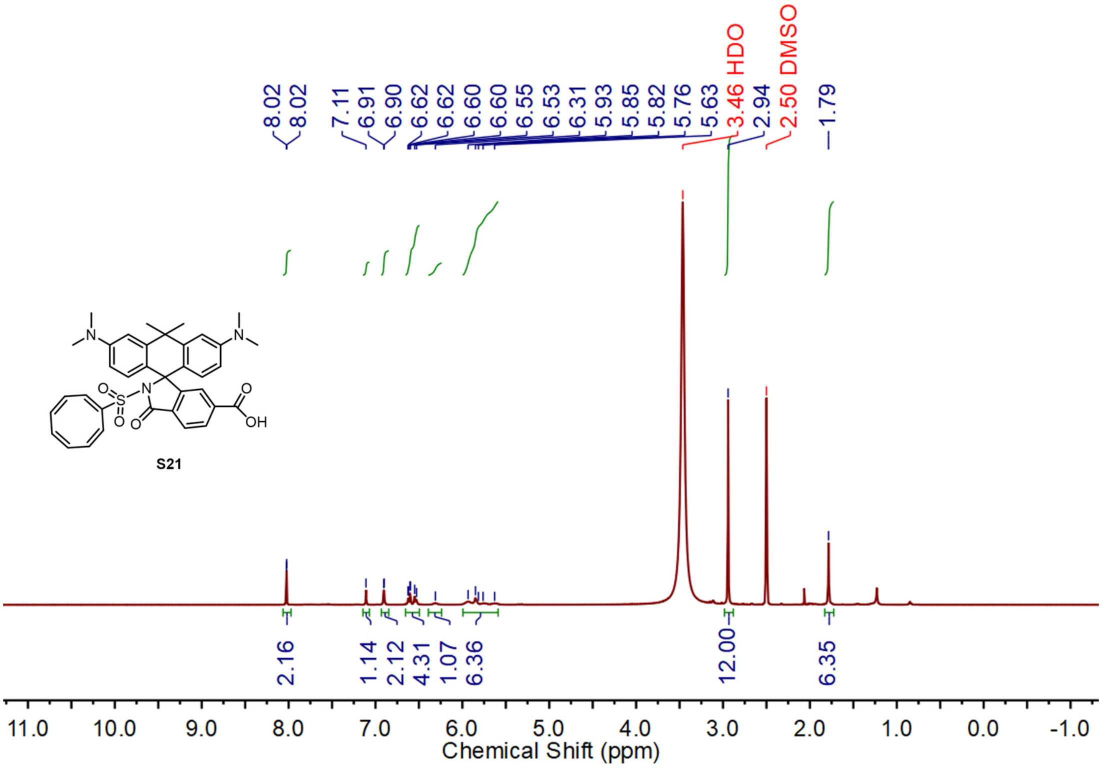


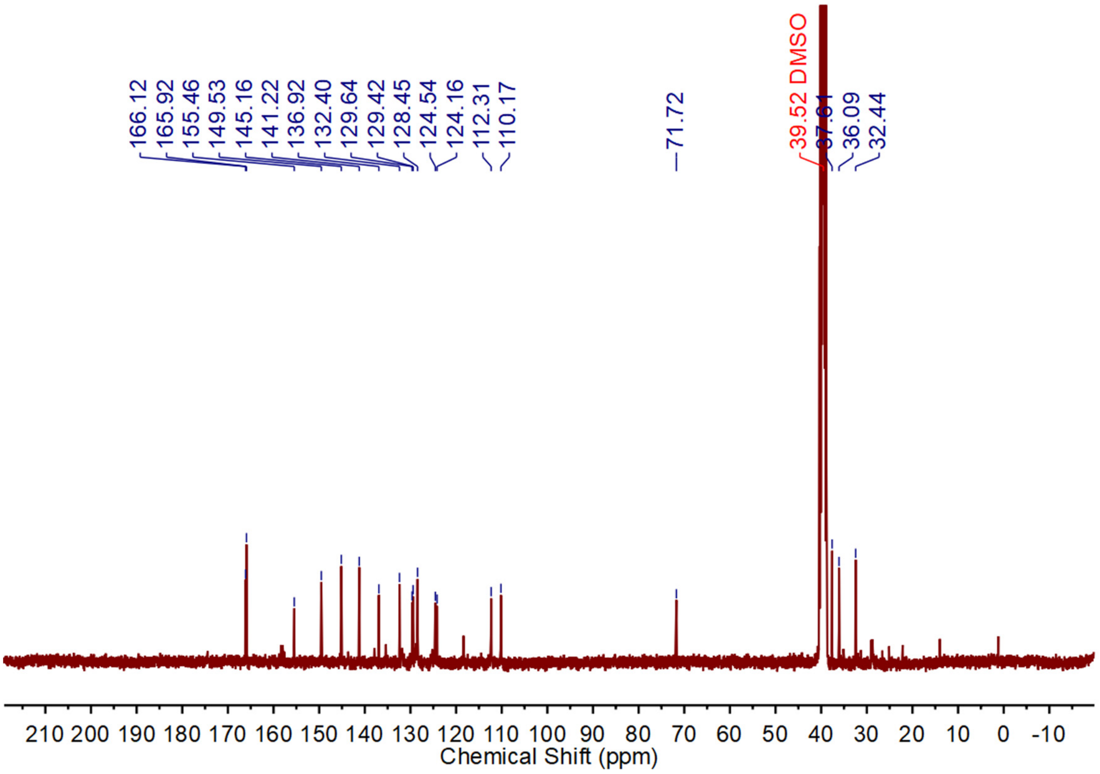


[1]H NMR and [13]C NMR spectra of compound **S21**

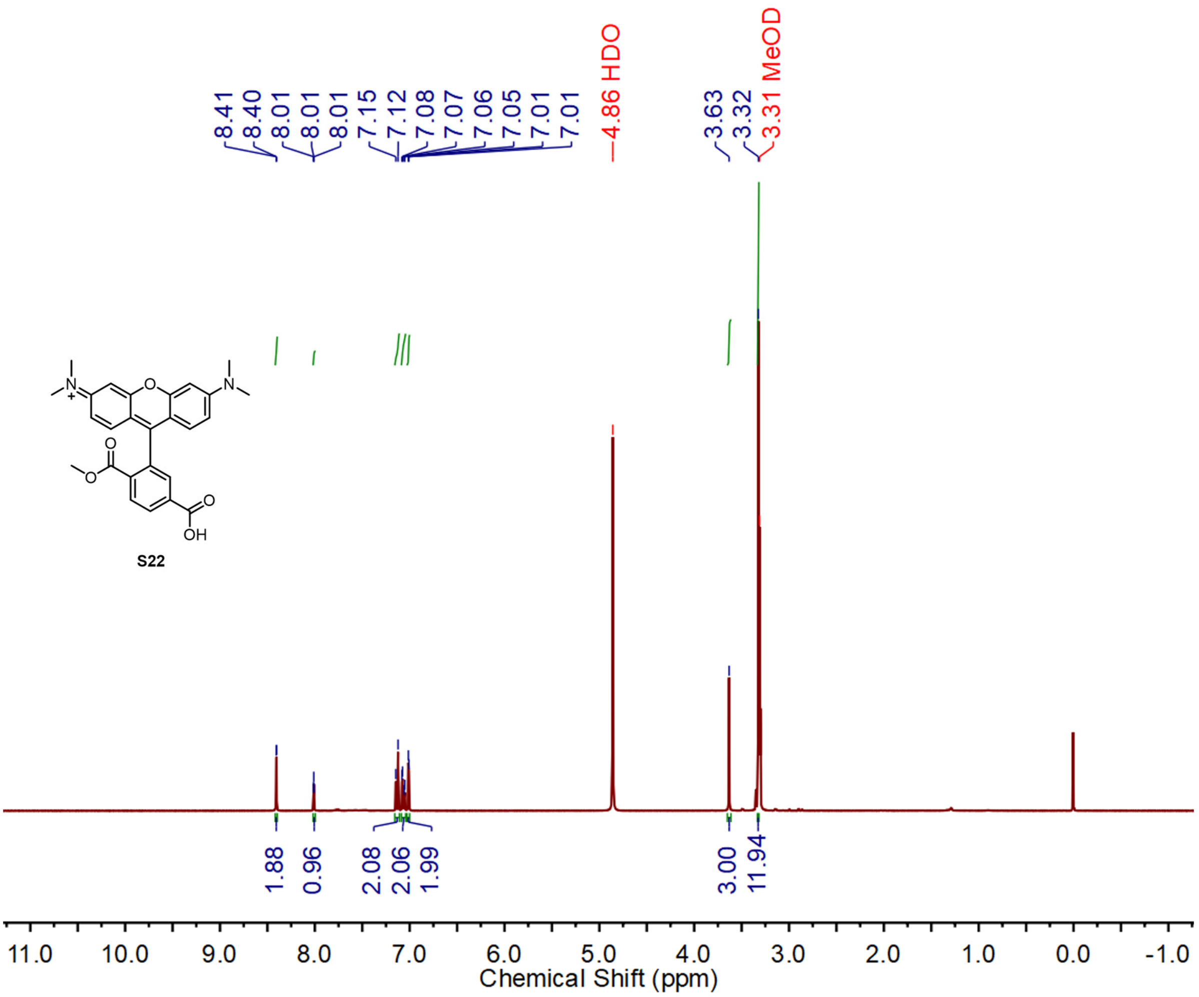


**S22**

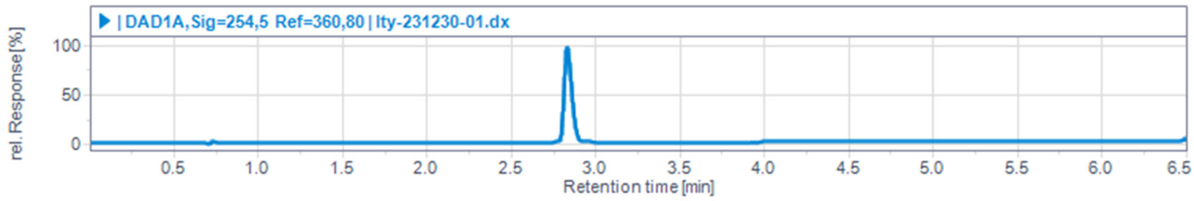


$^{1}$H NMR spectrum and LC/MS trace of compound **S22**

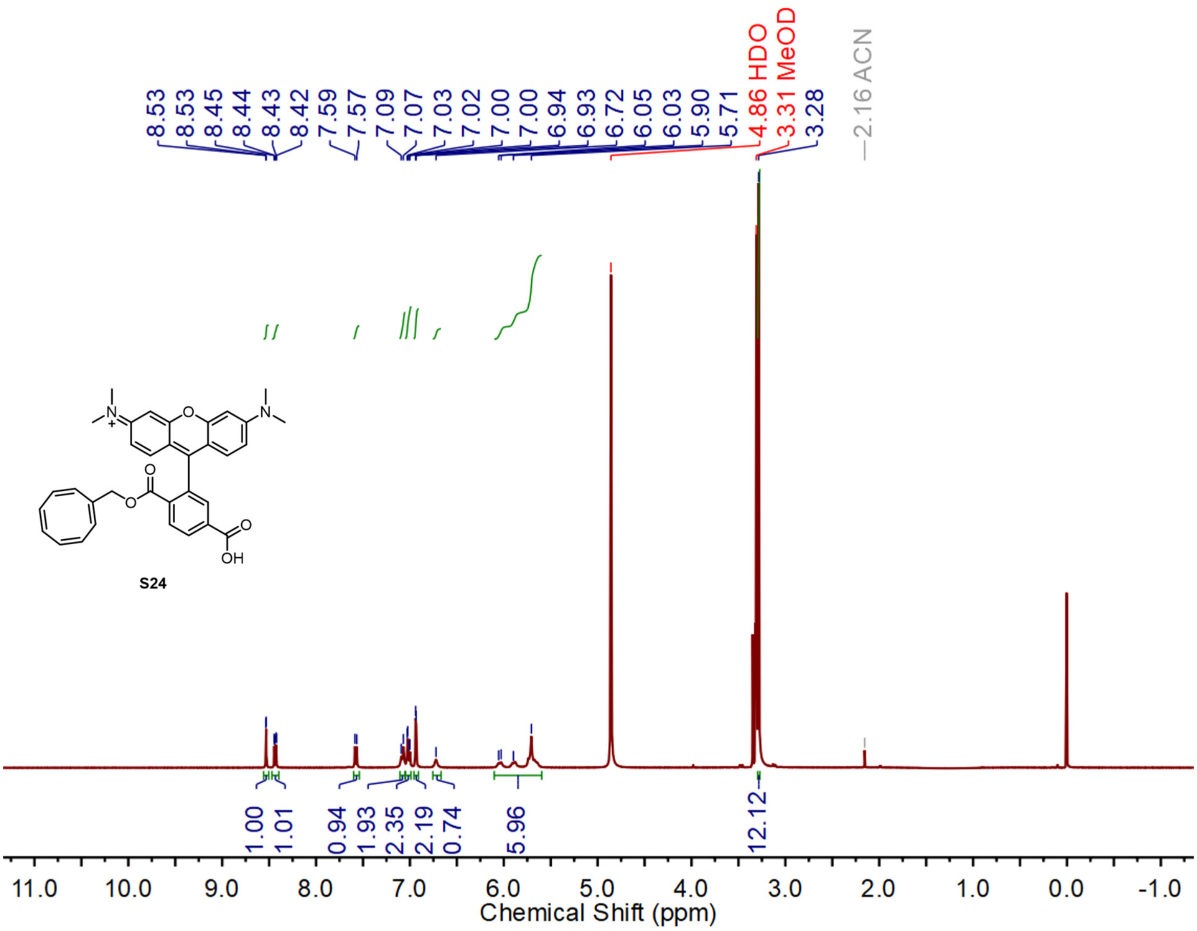


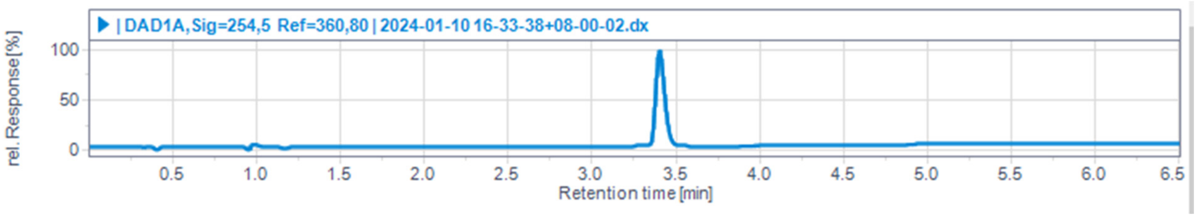


[1]H NMR spectrum and HPLC trace of compound **S24**

### Appendix

| | |
|---|---|
| aq | aqueous |
| oC | degrees Celsius |
| δ H | chemical shift in parts per million downfield from tetramethylsilane |
| DIPEA | N, N-diisopropylethylamine |
| DMF | dimethylformamide |
| eq | equivalent |
| ESI | electrospray ionization |
| Et | ethyl |
| g | gram(s) |
| HTL | HaloTag Ligand |
| HPLC | high-performance liquid chromatography |
| Hz | hertz |
| J | coupling constant (in NMR spectrometry) |
| LCMS | liquid chromatography mass spectrometry |
| μ | micro |
| m | multiplet (spectral); meter(s); milli |
| $M^+$ | parent molecular ion |
| Me | methyl |
| MHz | megahertz |
| min | minute(s) |
| mol | mole(s); molecular (as in mol wt) |
| mL | milliliter |
| MS | mass spectrometry |
| N | normal (equivalents per liter) |
| nm | nanometer(s) |
| NMR | nuclear magnetic resonance |
| pH | potential of hydrogen; a measure of the acidity or basicity of an aqueous solution |
| r.t. | room temperature |
| s | singlet (spectral) |
| t | triplet (spectral) |
| THF | tetrahydrofuran |

### Reference


1 Schindelin, J. *et al.* Fiji: an open-source platform for biological-image analysis. *Nat. Methods* **9**, 676-682 (2012).

2 Redmond, R. W. & Gamlin, J. N. A Compilation of Singlet Oxygen Yields from Biologically Relevant Molecules. *Photochem. Photobiol.* **70**, 391-475 (1999).

3 Battley, E. H. Escherichia Coli and Salmonella Typhimurium. Cellular and Molecular Biology, Volume 1; Volume 2. Frederick C. Neidhardt , John L. Ingraham , Boris Magasanik , K. Brooks Low , Moselio Schaechter , H. Edwin Umbarger. *Q.Rev.Biol.* **63**, 463-464 (1988).

4 Wang, L. *et al.* A general strategy to develop cell permeable and fluorogenic probes for multicolour nanoscopy. *Nat. Chem.* **12**, 165-172 (2020).

5 Lukinavičius, G. *et al.* SiR–Hoechst is a far-red DNA stain for live-cell nanoscopy. *Nat. Commun.* **6**, 8497 (2015).

6 Critchfield, F. E., Gibson, J. A., Jr. & Hall, J. L. Dielectric Constant and Refractive Index from 20 to 35° and Density at 25° for the System Tetrahydrofuran—Water1. *J. Am. Chem. Soc.* **75**, 6044-6045 (1953).

7 Kompa, J. *et al.* Exchangeable HaloTag Ligafnds for Super-Resolution Fluorescence Microscopy. *J. Am. Chem. Soc.***145**, 3075-3083 (2023).

8 Zolotukhin, S. *et al.* Production and purification of serotype 1, 2, and 5 recombinant adeno-associated viral vectors. *Methods* **28**, 158-167 (2002).

9 Casini, A., Storch, M., Baldwin, G. S. & Ellis, T. Bricks and blueprints: methods and standards for DNA assembly. *Nat. Rev. Mol. Cell Biol.* **16**, 568-576 (2015).

10 Mo, J. *et al.* Third-Generation Covalent TMP-Tag for Fast Labeling and Multiplexed Imaging of Cellular Proteins. *Angew. Chem. Int. Ed.* **61**, e202207905 (2022).

11 Liu, T. *et al.* Multi-color live-cell STED nanoscopy of mitochondria with a gentle inner membrane stain. *Proc. Natl. Acad. Sci. U.S.A* **119**, e2215799119(2022).

12 Li, M. A. *et al.* The piggyBac Transposon Displays Local and Distant Reintegration Preferences and Can Cause Mutations at Noncanonical Integration Sites. *Mol. Cell. Biol.* **33**, 1317-1330 (2013).

13 Frei, M. S. *et al.* Photoactivation of silicon rhodamines via a light-induced protonation. *Nat. Commun.* **10**, 4580 (2019).

14 Dolomanov, O. V., Bourhis, L. J., Gildea, R. J., Howard, J. A. K. & Puschmann, H. OLEX2: a complete structure solution, refinement and analysis program. *J. Appl. Crystallogr.* **42**, 339-341 (2009).

15 Sheldrick, G. Crystal structure refinement with SHELXL. **71**, 3-8 (2015).

16 Redmond, R. W. & Gamlin, J. N. A compilation of singlet oxygen yields from biologically relevant molecules. *Photochem. Photobiol.* **70**, 391-475 (1999).

17 Echegoyen, L., Maldonado, R., Nieves, J. & Alegria, A. Spin distributions in bridged bis(cyclooctatetraene) anion radicals. Dicyclooctatetraenylmethane and dicyclooctatetraenyldimethylsilane. *J. Am. Chem. Soc.* **106**, 7692-7695 (1984).

18 Obukhova, E. N. *et al.* Absorption, fluorescence, and acid-base equilibria of rhodamines in micellar media of sodium dodecyl sulfate. *Spectrochim. Acta A Mol. Biomol. Spectrosc.* **170**, 138-144 (2017).

19 Xu, S., Ma, W., Bai, Y. & Liu, H. Ultrasensitive Ambient Mass Spectrometry Immunoassays: Multiplexed Detection of Proteins in Serum and on Cell Surfaces. *J. Am. Chem. Soc.* **141**, 72-75 (2019).

20 Calitree, B. & Detty, M. Novel Rhodamine Dyes via Suzuki Coupling of Xanthone Triflates with Arylboroxins. *Synlett* **2010**, 89-92 (2010).

21 Yang, J. *et al.* Silicon-substituted rhodamine dyes and dye conjugates. WO2020033681 (2020).

22 Grimm, J. B. *et al.* A general method to improve fluorophores for live-cell and single-molecule microscopy. *Nat. Methods* **12**, 244-250 (2015).

23 Davies, T. Q., Tilby, M. J., Skolc, D., Hall, A. & Willis, M. C. Primary Sulfonamide Synthesis Using the Sulfinylamine Reagent N-Sulfinyl-O-(tert-butyl)hydroxylamine, t-BuONSO. *Org. Lett.* **22**, 9495-9499 (2020).

24 Lukinavičius, G. *et al.* A near-infrared fluorophore for live-cell super-resolution microscopy of

cellular proteins. *Nat. Chem.* **5**, 132-139 (2013).
